# Distinct Working Memory for Near and Far in a T-Maze Delayed Alternation Task: Evidence for Dual-Process Dynamics in Hippocampal-Prefrontal Coordination

**DOI:** 10.64898/2026.08.06.743185

**Authors:** Masatoshi Takita, Yukio Ichitani

## Abstract

We recently reported that rats performed better at a task distance of 2 m than at 0 m in a T-maze delayed alternation paradigm using a movable home cage in the longer-delay condition (Takita & Ichitani, 2026). We simultaneously recorded local field potentials from the bilateral prefrontal cortex, intermediate hippocampus, and ventral hippocampus. Across task epochs, coherence and two cross-frequency measures (phase-locking value and modulation index [MI]) revealed differences between correct and error trials in prefrontal interactions with hippocampal subregions. Among these measures, only MI was affected by task distance during the pre-task delay epoch. MI was highest in 2-m error trials and lowest in correct trials. In 0-m error trials, MI transiently increased during arm entry to levels comparable to those in 2-m error trials before declining toward the levels observed in correct trials during the later post-task delay. These MI dynamics appeared to be consistent with distance-dependent differences in behavioral performance. In addition, normalized Correct-Error Indices calculated for each electrophysiological measure revealed differential contributions of prefrontal coupling with the intermediate and ventral hippocampus across task distances. These findings suggest the existence of distinct near and far working memory states underlying distance-dependent behavioral differences, with distinct yet complementary contributions of the intermediate and ventral hippocampus to prefrontal interactions.

## Introduction

Working memory is known to depend on the prefrontal cortex and dopaminergic transmission in both monkeys and rats, suggesting that similar mechanisms are also present in humans (D’Esposito & Postle, 2015). However, differences in the duration of working memory have been observed across species. In humans and monkeys, working memory has typically been reported to persist for only a few minutes (Baddeley, 2012), whereas in rats it has been reported to extend from tens of minutes to several hours (Olton & Samuelson, 1976). Thus, despite the presumed common neural basis, substantial interspecies differences are observed at the behavioral level. Nevertheless, delayed alternation tasks in operant chambers have shown that rats can also perform working memory tasks with much shorter delays, on the order of several tens of seconds (Izaki et al., 2008).

Although delay duration is generally considered a key determinant of working memory performance, we previously found that, in a T-maze delayed alternation task, performance was unexpectedly better at a task distance of 2 m than at 0 m under longer-delay condition (Takita & Ichitani, 2026). This finding suggests that working memory performance may be influenced not only by delay duration but also by task-related demands associated with distance.

These behavioral effects of delay duration and task distance may, at least in part, reflect differences in the underlying neural circuitry. In rats, widespread regions of the posterior hippocampus project convergently to the prefrontal cortex (Takita et al., 2013). Evidence suggests that intermediate and ventral hippocampal projections to the prefrontal cortex make distinct contributions to working memory, although their roles depend on task demands. Specifically, disconnection of the intermediate (posterior dorsal) hippocampus-prefrontal cortex pathway impaired performance in an operant delayed alternation task (Izaki et al., 2008), whereas disruption of the ventral hippocampus-prefrontal cortex pathway impaired delayed alternation performance in maze tasks (Floresco et al., 1997; Wang & Cai, 2006). Evoked potential studies in the prefrontal cortex further suggest a hierarchical interaction between hippocampal inputs, in which ventral hippocampal input facilitates responses to intermediate hippocampal input, but not vice versa (Kawashima et al., 2006). Determining whether this functional asymmetry between the two pathways is engaged during behavior is essential for understanding how hippocampal inputs regulate prefrontal cortical function.

In the present study, we examined the effects of delay duration and task distance on working memory performance (Takita & Ichitani, 2026). Using electrophysiological recordings during a working memory task, we investigated how task distance influences prefrontal interactions with the intermediate and ventral hippocampus.

## Results

### Behavioral comparison of Naïve and surgically implanted Electrophysiology groups

Behavioral performance was compared between a Naïve group from our previous preprint study (Takita & Ichitani, 2026) and a surgically implanted Electrophysiology group (Fig. 1). In the Electrophysiology group, electrodes were implanted bilaterally into six brain regions: the prefrontal cortex (PC), intermediate hippocampus (iHP), and ventral hippocampus (vHP) (Fig. S1). Surgery was performed after animals had reached the 80% training criterion, and behavioral testing resumed following postoperative recovery. As shown in Fig. 1, both groups exhibited a similar behavioral pattern, with a greater reduction in performance under the 0 m condition than under the 2 m condition.

**Figure 1.**
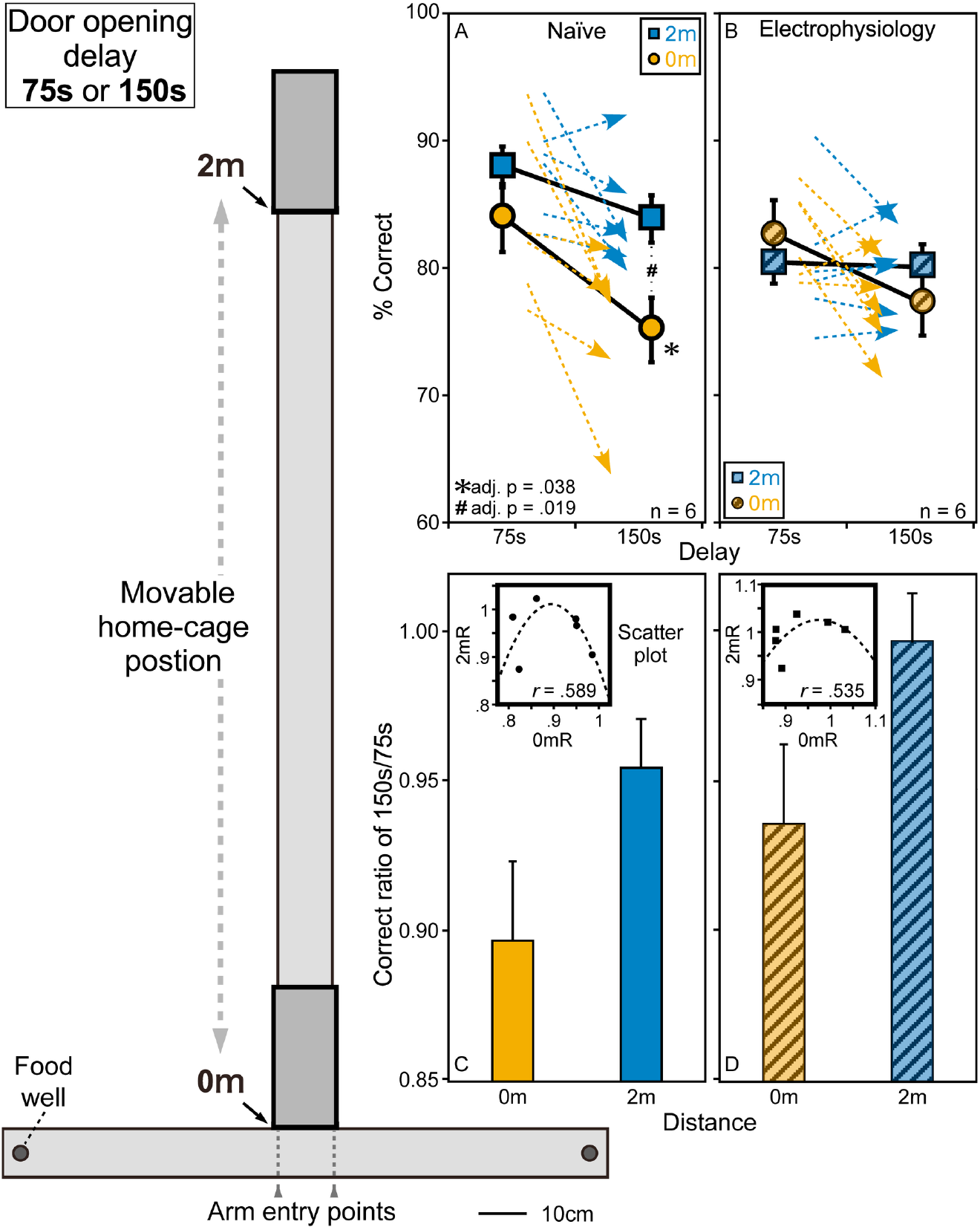
Comparison of behavioral accuracy between the Naïve and Electrophysiology groups. A schematic diagram of the T-maze used is also presented. Dashed vertical lines indicate the predefined arm entry points. Data from the “Naïve” group (A)—sourced from a previous study (Takita & Ichitani, 2026)—were used as reference data for comparison with the groups subjected to electrophysiological analysis (n = 6 per group). Dashed arrows (orange, 0 m; cyan, 2 m) indicate within-subject changes in correct responses from 75 s to 150 s. Circles and squares represent the 0 m and 2 m conditions, respectively; filled and hatched symbols denote the Naïve (A) and Electrophysiology (B) groups. The Naïve group showed significant main effects of delay and task distance without interaction, whereas no significant effects were detected in the Electrophysiology group. Under the 0 m condition, performance tended to decline with increasing delay, whereas no comparable tendency was observed under the 2 m condition. Bar graphs compare correct-response ratios (150 s/75 s) between the two conditions (C, D). Insets show an inverted-U relationship between the correct-response ratios, modeled by quadratic regression (see Results).

To evaluate the reduction in correct-response rate between the 75-s and 150-s retention intervals, a two-way mixed analysis of variance (ANOVA) was performed with task distance (0 m vs. 2 m) as a within-subject factor and group (Naïve vs. electrophysiology) as a between-subject factor. A significant main effect of task distance was observed, indicating that the reduction in correct-response rate was greater under the 0 m condition than under the 2 m condition (F(1,10) = 6.18, p = 0.032). Neither the main effect of group nor the interaction between task distance and group was significant (interaction: F(1,10) = 0.007, p = 0.94). Consistent with these findings, a direct comparison of the distance effect (0 m vs. 2 m) revealed no significant difference between the Naïve group (M = 4.74, 95% CI [−3.88, 13.37]) and the Electrophysiology group (M = 5.07, 95% CI [−0.27, 10.40]) (Welch’s t-test: t(9.73) = −0.08, p = 0.94, Cohen’s d = −0.05) (Fig. 1A,B).

Bar plots of the correct-response ratio (150 s/75 s) further demonstrated that both groups tended to exhibit higher ratios under the 2 m condition than under the 0 m condition (Fig. 1C,D). Scatter plots of the correct-response ratios under the two task distances (Fig. 1C,D, insets) revealed an inverted U-shaped relationship in both groups, indicating that the relationship between the correct-response ratios under the 0 m and 2 m conditions deviated from a simple linear correspondence in a similar manner across groups. Quadratic regression yielded the following equations: Naïve group, y = −11.79x² + 21.06x − 8.39 (R² = 0.35, RMSE = 0.041); Electrophysiology group, y = −5.73x² + 11.20x − 4.45 (R² = 0.29, RMSE = 0.031). The difference between the fitted curves was small (RMSE = 0.097), indicating similar overall curve shapes. Consistent with this observation, comparison of the quadratic regression coefficients revealed no significant difference between groups (F(2,6) = 2.31, p = 0.159). Together, these findings indicate that the relationship between the correct-response ratios under the 0 m and 2 m conditions did not differ significantly between the Naïve and Electrophysiology groups. Based on these results, we next examined neural activity associated with working memory performance.

### Local field potential (LFP) power during the 150-s delay period in the home cage

To characterize LFP signals in the prefrontal cortex (PC), intermediate hippocampus (iHP), and ventral hippocampus (vHP), LFP power was analyzed during the 150-s delay period in the home cage. For analysis, the delay period was divided into two consecutive 75-s periods, designated D1 (first 75 s) and D2 (second 75 s). Although a 75-s delay condition was included in the behavioral experiment, it was not analyzed separately; instead, D1 from the 150-s delay trials was used for comparison. LFP power (1-300 Hz) was calculated using 1-s time windows and then averaged into consecutive 5-s bins. These values were subsequently used for all statistical analyses. The data were classified by trial outcome (correct or error), task distance (0 m or 2 m), arm entry (left or right arm) and hemisphere (left: Fig. S2; right: Fig. S3), and visualized separately for each brain region. Across all three brain regions in both hemispheres, power spectra consistently exhibited a prominent peak at approximately 8-10 Hz. Accordingly, subsequent analyses focused on the 8-10 Hz and 51-100 Hz frequency bands.

### Power changes at 8-10 Hz and 51-100 Hz during the 150-s delay period in the home cage

Repeated-measures ANOVAs were performed according to the analytical framework summarized in Table S1. Across all six recording sites, significant effects of time were frequently detected in both the 8-10 Hz and 51-100 Hz bands. In contrast, significant main effects of trial outcome, task distance, or arm direction were not observed, although significant interactions involving these behavioral factors were occasionally detected.

Figure S4 illustrates temporal changes in LFP power throughout the recording period. Although these changes did not reach statistical significance, the post hoc time-course plots suggested a similar temporal pattern across all six regions in the 8-10 Hz band. Specifically, LFP power transiently decreased for approximately 30 s immediately after home-cage entry and subsequently increased in the hippocampal regions around home-cage exit, whereas no comparable increase was observed in the PC. In contrast, in the 51-100 Hz band, LFP power in the PC gradually decreased during the first approximately 30 s after home-cage entry, whereas no obvious temporal changes were observed in the hippocampal regions.

### Coherence changes at 8-10 Hz and 51-100 Hz between the PC and iHP/vHP during the 150-s delay period

Using the analytical framework summarized in Table S2, repeated-measures ANOVAs were performed to examine coherence in the 8-10 Hz and 51-100 Hz bands between the PC and iHP and between the PC and vHP in both hemispheres. Significant main effects and interactions involving behavioral factors, including trial outcome (correct/error [C/E]) and task distance (0 m/2 m), were detected only occasionally.

Figure S5 illustrates temporal changes in coherence across the entire 170-s recording period, including the 150-s home-cage delay epoch. Post hoc comparisons (Bonferroni-Dunn method) were performed, where appropriate, separately for the first and second 75-s periods. Significant differences between correct and error trials were detected in the 8-10 Hz and/or 51-100 Hz bands for the LPC-LiHP, LPC-LvHP, RPC-RiHP, and RPC-RvHP pairs. Across these region pairs, coherence was consistently lower during correct than during error trials, with the difference most pronounced during the first 75-s period.

### Phase-locking value (PLV) and modulation index (MI) changes between iHP/vHP (8-10 Hz) and PC (51-100 Hz) during the 150-s delay period

Repeated-measures ANOVAs were performed according to the analytical framework summarized in Table S3. Significant main effects of trial outcome were consistently observed, whereas significant main effects of task distance were not.

Figure 2 illustrates temporal changes in PLV and MI during a 170-s home-cage recording. Quantitative analyses were performed using the 150-s interval indicated in the figure. Post hoc comparisons (Bonferroni-Dunn method) were performed separately for D1 and D2. Significant differences between correct and error trials were detected for both PLV and MI in the LiHP-LPC, LvHP-LPC, RiHP-RPC, and RvHP-RPC pairs. Across all four pairs, both PLV and MI were consistently lower during correct than during error trials, and these differences persisted throughout the 150-s delay period. Of the two indices, only MI showed significant effects of task distance across all four PC-hippocampal pairs, with lower MI at 0 m than at 2 m during D2.

**Figure 2.**
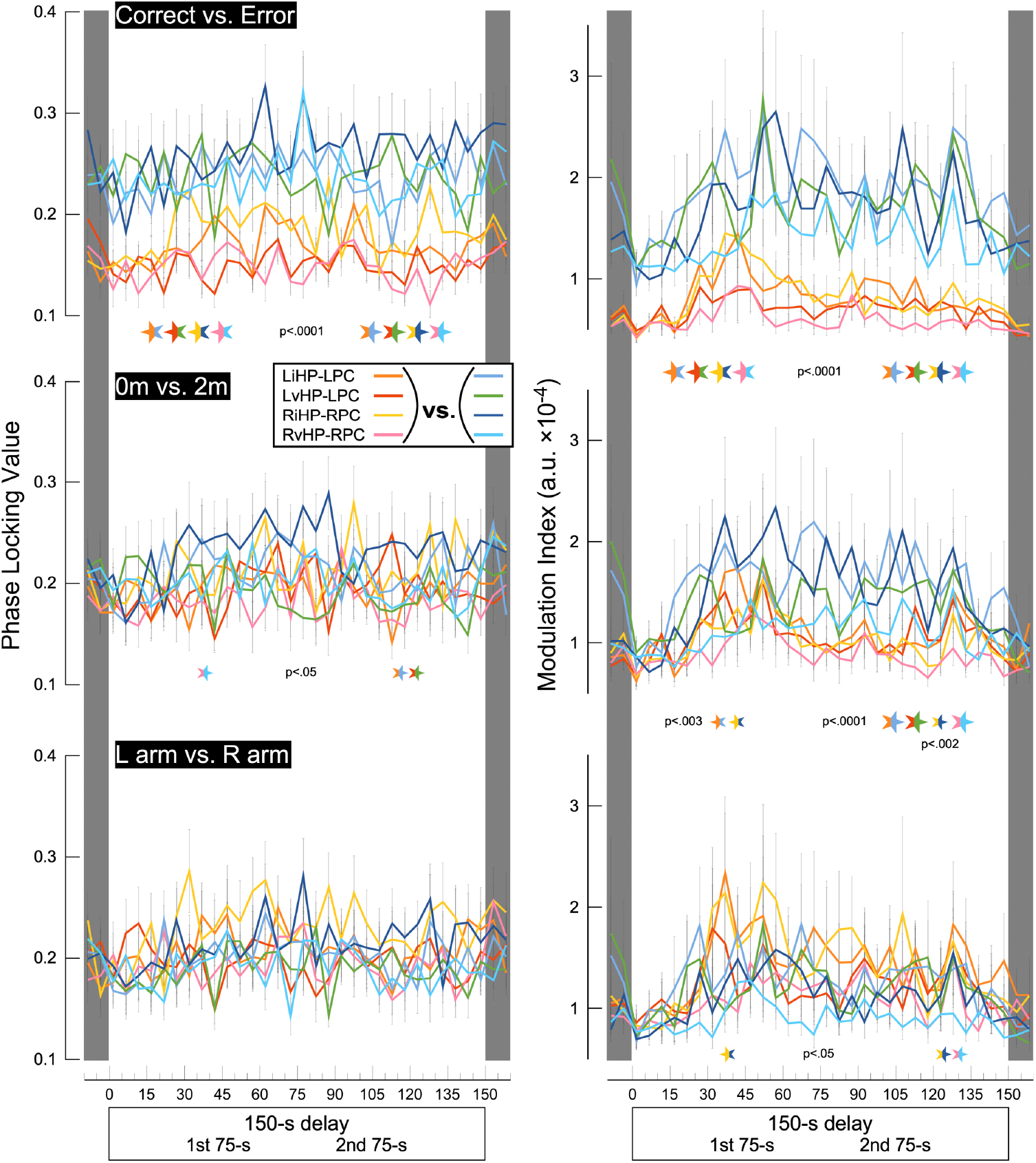
Time course of PLV and MI between iHP/vHP and PC in both hemispheres during a 150-s home-cage recording. Data are shown for PLV and MI between iHP/vHP (8-10 Hz) and PC (51-100 Hz) across three experimental factors (trial outcome: correct vs. error; task distance: 0 m vs. 2 m; and arm entry: L vs. R), using the same color coding as Figure S4 (see insets). Statistical analysis (Table S3) incorporated consecutive 5-s bins and two periods corresponding to the first and second 75-s periods. Post hoc comparisons between the two levels of each factor were performed separately for the first and second 75-s periods using the Bonferroni-Dunn test. The size of the significance markers is the same as in Figure S5. Four hippocampal (iHP/vHP)-PC MI pairs showed significant effects of task distance (0 m vs. 2 m), primarily during the second 75-s period. Among the electrophysiological measures analyzed directly (i.e., not CEIs), only MI showed significant effects of task distance across all four hippocampal-PC pairs.

### LFP power changes around T-maze arm entry

Neural activity during maze performance was analyzed over a 10-s period surrounding arm entry, divided into two consecutive 5-s epochs: A1 (5 s before arm entry) and A2 (5 s after arm entry). Repeated-measures ANOVAs were performed according to the analytical framework summarized in Table S4. Significant effects in both the 8-10 Hz and 51-100 Hz bands were detected predominantly as interactions between behavioral and temporal factors rather than as corresponding main effects.

Figure S6 illustrates changes in LFP power around arm entry. Although these changes did not reach statistical significance, the post hoc time-course plots suggested a similar pattern across all six regions in the 8-10 Hz band, characterized by a transient decrease lasting approximately 1 s immediately after correct-arm entry. In contrast, in the 51-100 Hz band, LFP power in the PC gradually decreased following correct-arm entry, whereas no apparent temporal changes were observed in the hippocampal regions.

### Coherence changes at 8-10 Hz and 51-100 Hz between the PC and iHP/vHP around T-maze arm entry

Repeated-measures ANOVAs were performed according to the analytical framework summarized in Table S5. Significant main effects and interactions involving behavioral factors were detected only occasionally.

Figure S7 illustrates temporal changes in coherence around arm entry. Post hoc comparisons (Bonferroni-Dunn method) were performed separately for A1 and A2. Significant differences between correct and error trials were detected in several region pairs in both the 8-10 Hz and 51-100 Hz bands. Across these region pairs, coherence was consistently lower during correct than during error trials, and significant differences were observed more frequently in the 51-100 Hz band than in the 8-10 Hz band.

### PLV and MI changes between iHP/vHP (8-10 Hz) and PC (51-100 Hz) around T-maze arm entry

Repeated-measures ANOVAs were performed according to the analytical framework summarized in Table S6. Significant main effects of trial outcome were consistently observed, whereas significant main effects of task distance were not.

Figure S8 illustrates temporal changes in PLV and MI around arm entry. Post hoc comparisons (Bonferroni-Dunn method) were performed separately for the A1 and A2 periods. Significant differences between correct and error trials were detected for both PLV and MI in all four hippocampal-prefrontal pairs. In every pair, both PLV and MI were consistently lower in correct than in error trials. The time-course plots revealed no obvious change in PLV around arm entry, whereas MI displayed a transient elevation around arm entry across all four hippocampal-PC pairs under both 0 m and 2 m conditions.

### Interaction effects of trial outcome and task distance on MI in iHP/vHP-PC coupling (8-10 Hz hippocampal phase, 51-100 Hz prefrontal amplitude) across task epochs

For MI during the 150-s pre-task (home-cage) delay epoch (Fig. 2), no significant hemispheric differences were observed between LiHP-LPC and RiHP-RPC (F(1,94)=0.096, p=0.76) or between LvHP-LPC and RvHP-RPC (F(1,94)=1.041, p=0.31). Therefore, subsequent MI analyses were conducted using pooled data across hemispheres. To enable comparisons across the three task epochs (pre-task delay, arm entry, and post-task delay), the same analyses were also performed for the arm entry epoch (Fig. S8) and the post-task delay epoch.

A repeated-measures ANOVA revealed no significant main effect of coupling (iHP-PC vs. vHP-PC) or coupling-related interactions in any epoch (all p > 0.05), indicating comparable coupling dynamics between the two hippocampal-prefrontal couplings. In contrast, significant main effects of trial outcome were consistently observed across the pre-task (F(1,184)=33.64, p<0.0001), arm entry (F(1,184)=33.71, p<0.0001), and post-task (F(1,184)=45.87, p<0.0001) epochs. A significant main effect of task distance was detected only during the pre-task epoch (F(1,184)=5.51, p=0.020), together with a significant trial outcome × distance interaction (F(1,184)=4.27, p=0.040). Because this interaction represented the principal distance-dependent effect observed across all electrophysiological analyses, the temporal evolution of MI was examined in detail.

Figure 3 summarizes the temporal dynamics of this study by illustrating how hippocampal-prefrontal MI changed across task epochs under different task distances and trial outcomes. Significant effects within the temporal subdivisions (D1/D2 and A1/A2) were further examined using Bonferroni-Dunn post hoc multiple comparisons. For both iHP-PC and vHP-PC couplings, MI was significantly higher in error than in correct trials throughout all three task epochs. Furthermore, during error trials, MI was significantly higher at 2 m than at 0 m. In the 0-m condition, error trials exhibited a distinctive temporal profile: before arm entry, MI values were comparable to those in correct trials but increased approximately twofold during the post-entry epoch, approaching the levels observed in the 2-m error condition. They then decreased significantly from D1 to D2 during the post-task delay epoch. These temporal profiles appeared to reflect the behavioral effects of task distance.

**Figure 3.**
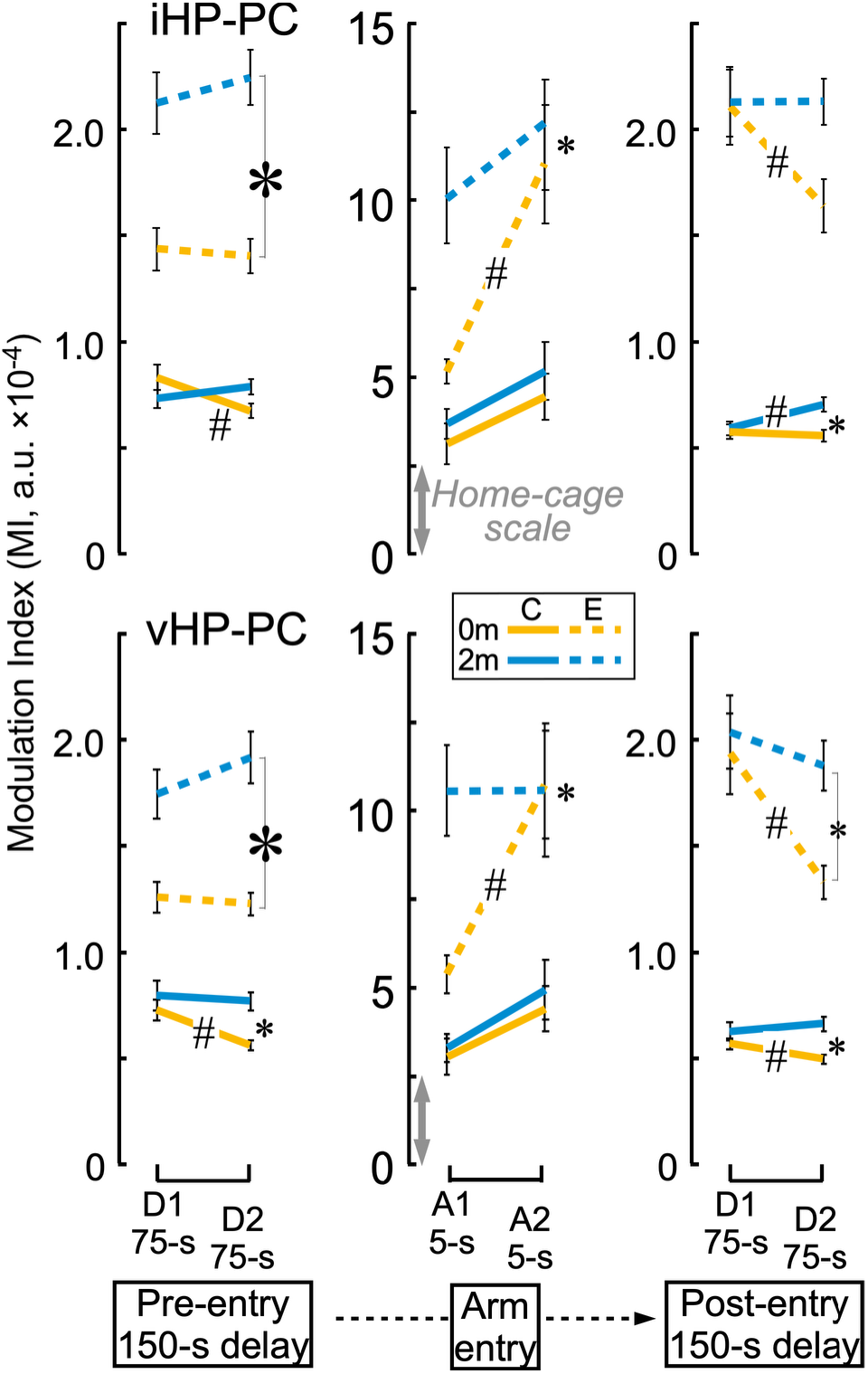
The temporal dynamics of MI in hippocampal-prefrontal coupling across task periods under different task distances and trial outcomes. D1/D2 and A1/A2 indicate the two delay periods and two arm-entry periods, respectively. Based on the ANOVA results, MI values for iHP-PC and vHP-PC coupling were plotted for each temporal subdivision, followed by Bonferroni-Dunn post hoc comparisons. The y-axis scale for the arm-entry epoch is sixfold larger than that for the delay epochs. Asterisks indicate significant differences between the 0-m and 2-m conditions (large: p < 0.001; small: p = 0.0005-0.046), and # indicates significant differences between D1 and D2 or between A1 and A2 (p = 0.0007-0.032).

### Differential temporal dynamics of normalized Correct-Error Indices (CEIs) derived from coherence, PLV, and MI in iHP/vHP-PC coupling across task epochs and task distances

To further characterize differences between iHP-PC and vHP-PC coupling, Correct-Error Index (CEI) values, calculated as (correct − error)/(correct + error), were derived for coherence (COH8-10 and COH51-100), PLV, and MI. Using the same temporal subdivisions (D1/D2 and A1/A2) as in Fig. 3, CEI values were analyzed by repeated-measures ANOVA.

For COH51-100, significant main effects of coupling (iHP-PC vs. vHP-PC) were observed during both the task epoch (F(1,92)=16.62, p<0.0001) and the post-task epoch (F(1,92)=5.39, p=0.022), with post hoc comparisons revealing significant differences between iHP-PC and vHP-PC at both distances (0 m and 2 m; Fig. 4). During the pre-task epoch, Bonferroni-Dunn post hoc comparisons revealed a significant difference between iHP-PC and vHP-PC coupling at 2 m (p=0.0352), but not at 0 m. COH8-10 also exhibited a similar pattern to that observed for COH51-100, although the effects were more limited.

**Figure 4.**
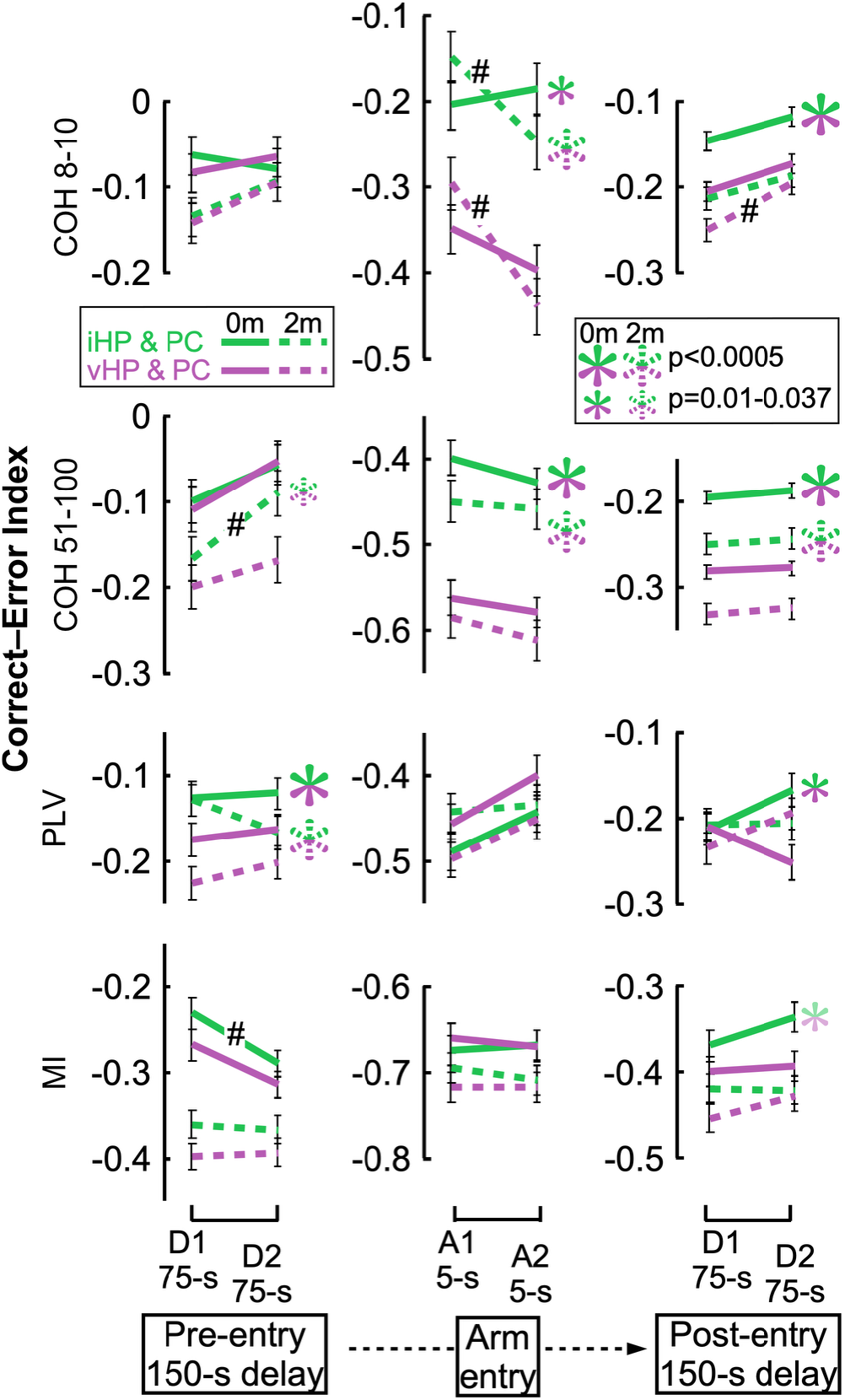
Temporal dynamics of the Correct-Error Index (CEI) for coherence (8-10 Hz and 51-100 Hz; COH8-10 and COH51-100), PLV, and MI. CEI values are shown using the same temporal subdivisions and Bonferroni-Dunn post hoc comparisons as in Figure 3. Negative CEI values indicate stronger interactions during error than during correct trials. Insets show iHP-PC and vHP-PC coupling in green and purple, respectively, with solid and dashed lines indicating the 0-m and 2-m conditions, respectively. The second inset summarizes differences between iHP-PC and vHP-PC coupling at each task distance. Significant differences between D1 and D2 or between A1 and A2 are indicated by # (p = 0.0001-0.036). A faint asterisk indicates the exploratory Bonferroni-Dunn comparison between iHP-PC and vHP-PC coupling for MI at 0 m during the post-task epoch (adjusted p = 0.0107).

For PLV, a significant main effect of coupling was observed during the pre-task epoch (F(1,92)=6.87, p=0.010). Bonferroni-Dunn post hoc comparisons confirmed significant differences between iHP-PC and vHP-PC coupling at both 0 m (p=0.0158) and 2 m (p=0.0005) (Fig. 4). During the epoch, neither the main effect of coupling nor the coupling × distance interaction reached significance (both p > 0.05). During the post-task epoch, a significant coupling × distance interaction was detected (F(1,92)=4.61, p=0.034), and subsequent post hoc comparisons revealed a significant difference between the two couplings at 0 m (p=0.0365), but not at 2 m (Fig. 4).

For MI, significant main effects of distance were observed during the pre-task epoch (F(1,92)=6.73, p=0.011) and the task epoch (F(1,92)=3.96, p=0.050). No significant main effect of coupling or coupling-related interactions was detected in any epoch (all p > 0.05). Although no significant omnibus effect involving coupling was detected, an exploratory Bonferroni-Dunn comparison suggested a difference between iHP-PC and vHP-PC coupling at 0 m during the post-task epoch (adjusted p = 0.0107; Fig. 4).

To further characterize significant temporal effects, Bonferroni-Dunn post hoc multiple comparisons were performed within each temporal subdivision (D1/D2 and A1/A2). The resulting temporal profiles are presented in Fig. 4 for the pre-task delay (D1/D2), arm entry (A1/A2), and post-task delay (D1/D2) epochs.

Overall, the CEI analysis demonstrated that iHP-PC and vHP-PC coupling exhibited distinct patterns of correct-error modulation. The differences between the two coupling pathways were not uniform across conditions but varied depending on task distance and temporal epoch, indicating differential regulation of hippocampal-prefrontal coupling across task contexts.

## Discussion

### Behavioral validation and overview of neural dynamics

Behavioral performance in the Electrophysiology group was broadly consistent with that of the Naïve group (Takita & Ichitani, 2026), preserving the characteristic inverted U-shaped behavioral relationship across task distances despite some quantitative differences (Fig. 1). The preservation of this inverted U-shaped relationship suggests that the behavioral organization underlying successful performance was maintained despite the surgical procedures. Notably, the non-linear relationship between the 0 m and 2 m conditions indicates that performance in the 2 m task cannot be explained as a simple quantitative extension of performance in the 0 m task. Rather, it suggests that increasing task distance may require the recruitment of a qualitatively different behavioral strategy. Because this characteristic behavioral relationship was preserved in the Electrophysiology group, this cohort provided an appropriate model for investigating the neural mechanisms supporting this distance-dependent behavioral adaptation. We therefore recorded local field potentials from the PC, intermediate hippocampus, and ventral hippocampus (Fig. S1). We examined oscillatory activity, interregional coordination, and cross-frequency interactions using analyses of power, coherence, phase-locking value (PLV), and modulation index (MI) (Figs. 2-4 and S4-S8; Tables S1-S6).

### Neural dynamics during the home-cage delay epoch

During the 150-s delay period, band-limited power showed distinct temporal patterns across regions. In the 8-10 Hz band, power exhibited a similar temporal profile across recording sites, whereas in the 51-100 Hz band, changes were more pronounced in the PC (Table S1, Fig. S4).

Although behavioral variables produced relatively few effects on band-limited power, analyses of functional interactions revealed clear differences between correct and error trials. Coherence within the 8-10 Hz and 51-100 Hz bands was lower during correct trials across multiple PC-hippocampal pairs (Table S2, Fig. S5). Likewise, both the phase-locking value (PLV) and modulation index (MI) between hippocampal 8-10 Hz signals and prefrontal 51-100 Hz signals were lower during correct trials (Table S3, Fig. 2). These findings indicate that successful performance was associated with reduced functional coupling both within frequency bands and across hippocampal-prefrontal frequency bands. Among these measures, only MI consistently reflected task distance during the second half of the delay period (Fig. 2, right column). This selective sensitivity prompted further examination of the temporal dynamics of MI across task epochs.

### Neural dynamics around arm entry

Band-limited power around arm entry shared several features with those observed during the home-cage delay epoch, while also exhibiting task-related transient changes (Table S4, Fig. S6). Power in the 8-10 Hz band transiently decreased after arm entry across multiple regions, whereas 51-100 Hz power gradually decreased predominantly in the PC.

Coherence between the PC and iHP/vHP differed between correct and error trials in both frequency bands, with more frequent differences observed in the 51-100 Hz band (Table S5, Fig. S7).

Around arm entry, MI showed a transient increase that was not observed for PLV, suggesting that amplitude-phase interactions were selectively enhanced during behavioral execution.

### Distance-dependent dynamics of hippocampal-prefrontal coordination

Differences in band-limited power between correct and error trials were modest (Tables S1 and S4; Figs. S2-S4 and S6). In contrast, functional interaction measures more clearly distinguished correct and error trials across both task epochs (Figs. 2 and 3).

Among these measures, MI showed additional sensitivity to task distance. In the 0-m error condition, MI increased around arm entry to a level comparable to that in the 2-m error condition, then gradually declined during the post-task period (Fig. 3). Notably, although MI was generally higher during error trials, the 2-m error condition was associated with better behavioral performance. These results indicate that hippocampal-prefrontal phase-amplitude coupling is not simply related to error processing or performance degradation, but instead reflects task-dependent coordination that varies with behavioral demands and temporal engagement. Importantly, this dissociation between MI and behavioral performance further suggests that these interactions cannot be interpreted solely in terms of behavioral success.

This interpretation is consistent with previous studies showing that hippocampal-prefrontal interactions contribute to cognitive control during memory retrieval (Ranganath et al., 2003; Ranganath, 2010; Rugg & Vilberg, 2013) and to context-dependent memory processing (Place et al., 2016; Eichenbaum, 2017). It is also compatible with the view that hippocampal-prefrontal interactions vary according to learning and behavioral demands (Takehara-Nishiuchi, 2020; Sun & Takehara-Nishiuchi, 2024). Similar observations have also been reported in humans, where hippocampal-prefrontal connectivity was not directly associated with behavioral performance (Iliopoulos et al., 2025).

Taken together, the present findings suggest that hippocampal-prefrontal MI reflects task-dependent interactions within the hippocampal-prefrontal network rather than behavioral success per se. This interpretation is also consistent with our previous proposal that task-intrinsic factors shape working memory processing (Takita & Ichitani, 2026). From the perspective of Cognitive Load Theory (Sweller, 1988), such task-intrinsic factors may represent properties of the task that contribute to differences in the cognitive demands imposed on the working memory system. Although the present study was not designed to directly test this framework, this perspective provides a useful conceptual account of how task distance may influence working memory processing.

### Correct-Error Index (CEI) Reveals Measure-Specific Functional Differences

CEI analysis revealed that synchronization measures captured distinct aspects of hippocampal-prefrontal network organization. In particular, CEI values derived from high-frequency coherence (COH51-100) showed the clearest differentiation between iHP-PC and vHP-PC during task execution under both distance conditions, whereas low-frequency coherence (COH8-10) also exhibited significant coupling-related differences. These findings suggest that different frequency components of coherence may capture distinct aspects of functional differentiation between iHP-PC and vHP-PC coupling during spatial choice. The persistence of coherence-related CEI differences into the post-task delay further suggests that task-related network states may be maintained after behavioral completion. In contrast, CEI values for PLV showed no coupling-specific differences during task execution but differed during pre-task and post-task delay epochs, suggesting that phase synchronization may be more closely related to network states established before task initiation and maintained after completion than to online processing during behavior. CEI analysis of MI provided limited evidence for coupling-specific effects, with only one adjusted post hoc comparison reaching significance in the 0 m post-task delay condition. Therefore, MI-related differences should be interpreted cautiously.

### Functional Implications of Differential Hippocampal-Prefrontal Coupling

Previous lesion studies have demonstrated that distinct hippocampal-prefrontal pathways contribute to working memory depending on behavioral demands. The iHP-PC pathway has been implicated in operant delayed alternation (Izaki et al., 2008), whereas disruption of the vHP-PC pathway impairs maze-based working memory performance (Floresco et al., 1997; Wang & Cai, 2006). Anatomical and physiological studies further support interactions between these pathways, showing convergent projections from intermediate and ventral hippocampal regions to the prefrontal cortex (Takita et al., 2013), and functional interactions between hippocampal subregions (Kawashima et al., 2006). Together, these findings suggest that iHP-PC and vHP-PC pathways are components of an integrated hippocampal-prefrontal network whose relative contribution varies according to behavioral context.

The present CEI findings extend previous lesion studies by demonstrating dynamic changes in functional coupling during maze-based working memory. Whereas lesion studies identify pathways necessary for successful performance, the present synchronization analysis reveals how hippocampal-prefrontal interactions are adaptively organized during behavior. The present results indicate that functional differentiation between iHP-PC and vHP-PC coupling depends on task distance and behavioral epoch. Specifically, greater differentiation during the 0 m condition suggests stronger functional specialization of iHP-PC and vHP-PC coupling under conditions requiring immediate spatial processing, whereas the 2 m condition showed a more balanced involvement of the two pathways. These findings suggest that maze-based working memory is supported not only by the vHP-PC pathway but also by task-dependent recruitment of the iHP-PC pathway.

These results further support our previous proposal that task distance functions as a task-intrinsic factor influencing working memory processing (Takita & Ichitani, 2026). The near-far distinction may represent different modes of working memory operation characterized by differential engagement of intermediate and ventral hippocampal-prefrontal pathways. Such differences may reflect adaptive reorganization of hippocampal-prefrontal interactions in response to task demands. This interpretation is consistent with Cognitive Load Theory.

### Limitations

Several methodological limitations should be considered when interpreting the present findings. First, surrogate-based validation was not performed for either the phase-locking value (PLV) or phase-amplitude coupling (PAC) analyses. Although relatively long recording epochs, amplitude normalization, and artifact exclusion procedures were employed to reduce potential analytical biases, residual influences arising from signal characteristics, waveform shape, and filtering procedures cannot be completely excluded. Because identical preprocessing and analytical procedures were applied across all experimental conditions, any such methodological influences would be expected to affect all conditions similarly. Therefore, the present findings should be interpreted primarily as reflecting relative differences between experimental conditions rather than absolute estimates of coupling strength.

Second, measures that are less sensitive to zero-lag interactions, such as imaginary coherence, were not included. Consequently, the potential influence of volume conduction or common-source activity cannot be completely excluded. Accordingly, the observed coherence, PLV, and PAC results should be interpreted as indices of functional synchronization rather than definitive evidence of direct neuronal coupling or causal interactions.

Finally, the present PLV analysis was intended to quantify the temporal stability of cross-frequency phase relationships between hippocampal 8-10 Hz signals and prefrontal 51-100 Hz signals.

## Conclusion

This study demonstrates that task distance is associated with distinct patterns of hippocampal-prefrontal coordination during working memory. The coordination patterns involving the intermediate and ventral hippocampus were observed within the common anatomical framework of convergent posterior hippocampal projections to the prefrontal cortex. These coordination patterns were not simply related to behavioral accuracy but reflected task-dependent functional organization within this neural system. Together, these findings indicate that task distance shapes the functional organization of hippocampal-prefrontal coordination across near and far task conditions.

## Materials and methods

### Animals

All experimental procedures were approved by the Animal Care and Use Committee of National Institute of Advanced Industrial Science and Technology (approval no. LS-00000799, LS-00000968, LS-00001276, LS-00001533, LS-00001894) and conducted in accordance with the NIH (1996) and Science Council of Japan (2006) guidelines. All experimental data were analyzed at the University of Electro-Communications.

Nine 6-week-old male Sprague-Dawley rats (CLEA, Tokyo, Japan) were purchased and housed individually. After a 3-4-week acclimation period, body weights ranged from 285 to 356 g at the start of behavioral training. Three rats were used in a preliminary study to validate the recording and analysis procedures and were excluded from the final analysis. The remaining six rats were used for all experiments. Rats were fed standard laboratory chow (CE-7, CLEA, Tokyo, Japan) with ad libitum access to water and were housed in a temperature-controlled room (24°C) under a 12-h light/dark cycle. During behavioral training, body weight was maintained at ≥85% of the free-feeding weight.

### Behavioral Task

The experimental procedures were based on a previously reported T-maze delayed alternation task protocol (Takita & Ichitani, 2026). Briefly, during training, the distance from the movable home-cage exit to the T-junction was set to 1 m, and 45-mg precision pellets (Bio-Serv, Flemington, NJ, USA; F0021-J) were used as rewards. The delay duration was gradually increased across sessions, starting at 30 s, until rats reached a criterion of ≥80% correct choices over multiple sessions at a delay of 113 s (approximately 1-2 weeks). Each training session began with an initial free-choice trial and continued until the rat completed 20 correct responses.

During testing, rats performed two sessions per day. In each session, combinations of distance (0 and 2 m) and delay (75 and 150 s) were presented in a pseudo-random order, with constraints preventing consecutive presentation of similar conditions (see Fig. 2 of Takita & Ichitani, 2026). Electrophysiological recordings were obtained over 2-5 recording cycles, each consisting of 8 consecutive days.

### Behavioral events

Arm entry was operationally defined as crossing the predefined arm entry point in the arm where the rat ultimately reached the food well (Fig. 1). This criterion was applied uniformly to all trials, including rare instances of vicarious trial-and-error behavior, in which rats briefly entered one arm but reversed before reaching the food well in that arm and subsequently reached the food well in the opposite arm. The same criterion was used to align all electrophysiological analyses to arm entry.

### Surgery and Electrode Implantation

Anesthesia was induced with pentobarbital (10-15 mg/kg, intraperitoneal injection) and maintained with sevoflurane (1.5-2%; NARCOBIT-E vaporizer, Natsume Seisakusho Co., Ltd., Japan). Animals were secured in a stereotaxic apparatus, and adequate anesthetic depth was confirmed by the absence of a tail-pinch reflex. Body temperature was maintained at 37°C using a feedback-controlled heating pad with rectal temperature monitoring.

Recording electrodes were constructed from PFA-coated platinum-iridium wires (A-M Systems, catalog no. 776000; bare wire diameter, 0.002 inch; coated diameter, 0.004 inch). Two wires were twisted together and reinforced with nail polish, resulting in a final diameter of approximately 0.008-0.01 inch. Electrode impedance ranged from 4 to 10 kΩ at 1 kHz in 0.9% NaCl at room temperature. Wire electrodes were stereotaxically implanted bilaterally into the medial prefrontal cortex (PC), intermediate hippocampus (iHP), and ventral hippocampus (vHP) according to Paxinos and Watson (1982). Stereotaxic coordinates (mm) were as follows: PC (AP +2.7 to +3.7, ML 0.6-1.0, DV 3.0-4.0), iHP (AP −4.8 to −6.3, ML 4.5-5.5, DV 2.5-4.0), and vHP (AP −6.0 to −6.7, ML 5.0-6.0, DV 5.0-7.0). Ipsilateral inter-electrode distances were approximately 9.7 mm (PC-iHP), 10.9 mm (PC-vHP), and 2.9 mm (iHP-vHP).

A 12-channel connector (Omnetics, A79018-001, Minneapolis, MN, USA) was attached to the recording electrodes and secured to the skull with anchor screws placed in the frontal and occipital bones (approximately AP +5 mm and AP −9 mm, respectively) using dental cement. While housed in the home cage, the connector was protected with a ∼5 g dust cover. After a 1-week recovery period, the dust cover was replaced with a wireless headstage transmitter (W16, Triangle BioSystems International, Durham, NC, USA; system gain, ∼800; band-pass filter, 0.8-7 kHz; weight, 4.5 g).

### Data Acquisition and Preprocessing

Local field potentials (LFPs) were recorded wirelessly at a sampling rate of 20 kHz using a common reference formed by electrically shorting two skull screws. Signals were downsampled to 1 kHz for subsequent analyses.

LFP signals were inspected for artifacts. Segments containing non-physiological noise (e.g., movement artifacts or signal saturation) were identified based on visual inspection and amplitude criteria. Brief transient artifacts exceeding ±2 standard deviations from the mean of each trial were corrected by linear interpolation when limited in duration.

After artifact correction, trials whose mean signal amplitudes exceeded ±2 standard deviations from the mean for each subject within each experimental condition were considered outliers and excluded from further analyses.

To reduce inter-trial variability and minimize spurious phase-amplitude coupling arising from broadband power fluctuations, LFP amplitudes were normalized by z-score transformation across all trials within each subject for each recording day before spectral and phase-amplitude coupling analyses.

### Frequency Bands and Analysis Epochs

Power spectral density was first computed over the frequency range of 1-300 Hz for each task distance (0 and 2 m), separately for the first and second 75 s of the 150 s delay period, correct and error trials, and the left and right iHP, vHP, and PC (Figs. S2 and S3). Based on the resulting spectral profiles, the 8-10 Hz and 51-100 Hz frequency ranges were interpreted as theta-range and high-gamma-range activity, respectively. To examine temporal changes in frequency-specific activity, analyses were performed during the home-cage delay period and around T-maze arm entry. The 150 s home-cage delay period was divided into two consecutive 75 s intervals, designated D1 (first 75 s) and D2 (second 75 s). Likewise, the 10 s period surrounding arm entry was divided into two consecutive 5 s intervals, designated A1 (−5 to 0 s before arm entry) and A2 (0 to +5 s after arm entry). Unless otherwise specified, electrophysiological measures were quantified separately for each epoch.

### Power Analysis

Power spectral density was estimated using Welch’s method with 1-s windows and 50% overlap. During the home-cage delay epoch, power values were averaged over consecutive 5-s time bins within D1 and D2. During the period, power values were averaged over consecutive 1-s time bins within A1 and A2. Band-limited power was calculated for the 8-10 Hz and 51-100 Hz frequency bands for each trial and normalized across trials within each recording session.

### Coherence Analysis

Functional connectivity between regions was assessed using magnitude-squared coherence. Coherence was computed for predefined intrahemispheric electrode pairs using Welch’s method. Coherence values were quantified using the same epoch definitions (D1/D2 and A1/A2) described above.

### Phase-locking value (PLV)

Cross-frequency phase relationships between hippocampal 8-10 Hz oscillations and prefrontal 51-100 Hz oscillations were quantified using the PLV. Hippocampal local field potentials (LFPs) were band-pass filtered in the 8-10 Hz range, whereas prefrontal LFPs were filtered in the 51-100 Hz range using zero-phase finite impulse response (FIR) filters. Instantaneous phase time series were extracted using the Hilbert transform. Cross-frequency PLV was calculated from the consistency of instantaneous phase relationships between hippocampal 8-10 Hz and prefrontal 51-100 Hz oscillations within each analysis epoch. Larger PLV values indicate greater temporal stability of the phase relationship across the epoch.

### Phase-Amplitude Coupling (PAC): Modulation Index (MI)

Phase-amplitude coupling was quantified using the modulation index (MI) described by Tort et al. (2010). The phase of hippocampal LFPs in the 8-10 Hz range and the amplitude envelope of prefrontal LFPs in the 51-100 Hz range were extracted using zero-phase finite impulse response (FIR) filters followed by the Hilbert transform.

### Statistical Analysis

Behavioral performance was evaluated as the percentage of correct responses for each condition, separately for the first and second halves of each daily session. Electrophysiological data were analyzed using repeated-measures ANOVAs with factor structures appropriate for each analysis (Tables S1-S6). For the home-cage delay analyses, repeated-measures ANOVAs included trial outcome (correct/error), task distance (0 m/2 m), arm direction (left/right), period (D1/D2), and within-period time as repeated factors. For the maze-task analyses, repeated-measures ANOVAs included trial outcome (correct/error), task distance (0 m/2 m), retention interval (75 s/150 s), arm direction (left/right), period (A1/A2), and within-period time as repeated factors. Thus, each analysis incorporated the appropriate combination of trial outcome, task distance, retention interval (analyses only), arm direction, and temporal factors according to the analytical design. Significant effects were followed by Bonferroni-Dunn multiple comparisons where appropriate. Welch’s t test was used for pairwise comparisons when the assumption of equal variances was not met. All statistical analyses were performed using StatView J-5 (Hulinks Inc., Tokyo, Japan). Statistical significance was defined as *P* < 0.05. Data are presented as means ± SE.

### Histology

Electrode locations were verified histologically after completion of the experiments by passing direct current through the electrodes. Brains were sectioned into 70-μm-thick serial sections and stained with thionin to confirm electrode placement.

## Acknowledgment

We thank the late Dr. Yoshinori Izaki (St. Marianna University School of Medicine), Drs. Hiroshi Yokoi and Yoshiko Yabuki (The University of Electro-Communications), for their valuable discussions, and Ms. Atsuko Yamashita (National Institute of Advanced Industrial Science and Technology) for her technical assistance. This work was supported by JSPS KAKENHI (15K04202 and 18K03196 to M.T.), and in part by an AIST grant for neurorehabilitation research (M.T.). Part of this work was conducted while M.T. was affiliated with National Institute of Advanced Industrial Science and Technology. We further express our appreciation to Dr. Sei-Etsu Fujiwara (St. Marianna University School of Medicine) for their dedicated and sincere efforts in electrophysiological data analysis. We are grateful for their continued collaboration on methodological development and additional analyses.

