## Supplementary Information for "Distinct Working Memory for Near and Far in a T-Maze Delayed Alternation Task: Evidence for Dual-Process Dynamics in Hippocampal-Prefrontal Coordination"

### Supplementary Methods S1-S7

#### S1. Apparatus and experimental setup

The behavioral apparatus was identical to that described previously (Takita & Ichitani, 2026). Briefly, a matte-black acrylic T-maze (12 cm corridor width, 1 cm wall thickness, 4 cm wall height) was installed at a height of 50 cm above the floor. The internal lengths of the T-maze corridors were 134 cm for the crossbar and 230 cm for the stem extending from the center of the crossbar. A movable home cage allowed adjustment of the task distance from 0 m to 2 m depending on the task condition.

Experiments were conducted under dim indirect illumination (9-10 lux) with continuous white noise (60-65 dB). An LED light (~150 lux) mounted above the home cage was illuminated at trial onset as a behavioral cue.
Behavior was continuously monitored using an overhead CCD camera.

#### S2. Detailed behavioral procedures

Each trial began by opening the door of the home cage and simultaneously turning on the LED above it. The rat then exited the cage and navigated the T-maze toward a food well. After reaching the food well, the LED was turned off, and the opposite arm was closed. The rat then returned to the cage, and the door was closed to initiate the delay period before the next trial.

To account for the rare instances of vicarious trial-and-error behavior, arm entry was operationally defined as crossing the predefined arm entry point in the arm where the rat ultimately reached the food well (Fig. 1).

**S3. Electrode construction and stereotaxic implantation details**

Recording electrodes were constructed from PFA-coated platinum-iridium wire (A-M Systems, USA; catalog no. 776000). Two wires were twisted together and reinforced to obtain a final diameter of approximately 0.008-0.01 inches. Electrode impedance ranged from 4-10 kΩ at 1 kHz in 0.9% saline.

Stereotaxic coordinates were as follows (in mm), based on the rat brain atlas (Paxinos & Watson, 1982): Prefrontal cortex (PC): AP +2.7 to +3.7, ML 0.6-1.0, DV 3.0-4.0; Intermediate hippocampus (iHP): AP −4.8 to −6.3, ML 4.5-5.5, DV 2.5-4.0; Ventral hippocampus (vHP): AP −6.0 to −6.7, ML 5.0-6.0, DV 5.0-7.0. Approximate ipsilateral inter-electrode distances were as follows (in mm): PC-iHP: 9.7, PC-vHP: 10.9, iHP-vHP: 2.9.

Electrodes were connected to a multi-channel connector (Omnetics, USA). Of the available channels, 12 were used for recording electrodes and 2 for skull screw electrodes. The assembly was fixed using dental cement and skull anchor screws and protected by a dust cover during recovery.

**S4. Data preprocessing**

LFP signals were visually inspected to identify movement artifacts, signal saturation, and non-physiological noise. Outlier handling was performed in two steps. First, within each trial, time points exceeding ±2 standard deviations from the trial mean were identified as transient artifacts and corrected using linear interpolation when brief.

Second, at the trial level, trials whose mean signal magnitudes exceeded ±2 standard deviations from the mean within each condition for each subject were excluded as outliers. Trials containing excessive noise were also excluded from subsequent analyses. Amplitude normalization was performed using z-score transformation within each subject for each recording day across all trials to reduce inter-trial variability and minimize the influence of broadband power differences, which can artificially inflate phase-amplitude coupling measures. This approach accounts for potential day-to-day variability in signal amplitude.

### S5. Spectral and connectivity analyses

#### Power spectral density

Power spectral density was estimated using Welch’s method with 1-s windows and 50% overlap. A broad frequency range (1-300 Hz) was first examined to characterize the overall spectral profiles (Figures S2-S3). Based on the observed spectral features, the 8-10 Hz and 51-100 Hz frequency ranges were interpreted as theta-range and high-gamma-range activity, respectively.

#### Coherence

Functional connectivity between brain regions was quantified using magnitude-squared coherence for predefined intrahemispheric electrode pairs between PC and iHP/vHP.

$$C_{xy}(f) = |S_{xy}(f)|^2 / (S_{xx}(f)S_{yy}(f))$$

where $S_{xy}(f)$ represents cross-spectral density and $S_{xx}(f)$, $S_{yy}(f)$ represent auto-spectral densities.

**Phase-locking value (PLV)**

Cross-frequency phase relationships between hippocampal 8-10 Hz oscillations and prefrontal 51-100 Hz activity were quantified using the phase-locking value (PLV). Hippocampal LFPs were filtered in the 8-10 Hz band, whereas prefrontal LFPs were filtered in the 51-100 Hz band using zero-phase finite impulse response (FIR) filters. Instantaneous phase time series were extracted using the Hilbert transform. For each analysis epoch, PLV was calculated from the consistency of the instantaneous phase relationship throughout the epoch, yielding a single PLV value for each recording segment. PLV values range from 0 to 1, with larger values indicating greater temporal consistency of the phase relationship.

**Phase-Amplitude Coupling (PAC): Modulation Index (MI)**

PAC between the phase of hippocampal 8-10 Hz oscillations and the amplitude of prefrontal 51-100 Hz activity was quantified using the modulation index (MI) based on Tort et al. (2010). The instantaneous phase of the hippocampal 8-10 Hz oscillation was divided into 18 bins (20° each), and the mean 51-100 Hz amplitude within each phase bin was used for MI estimation.

$MI = DKL(P \| U) / \log(N)$

where DKL represents Kullback-Leibler divergence between the observed distribution and the uniform distribution.

**S6. Time-resolved analysis around arm entry**

Neural activity was analyzed from −5 s to +5 s relative to arm entry using 1-s bins. Band-limited power, coherence, PLV, and MI were quantified separately for each time bin and compared across correct/error trials, distance conditions, delay conditions, and arm entry direction. This analysis was used to characterize dynamic neural coordination around arm entry (Figures S6-S8, Tables S4-S6).

**S7. Additional notes**

No significant baseline differences in spectral profiles were observed between correct and error trials. All analyses were performed using custom LabVIEW scripts (National Instruments) together with commercial statistical software (StatView J-5; Hulinks, Tokyo, Japan).

**Supplementary Figures S1-S8**

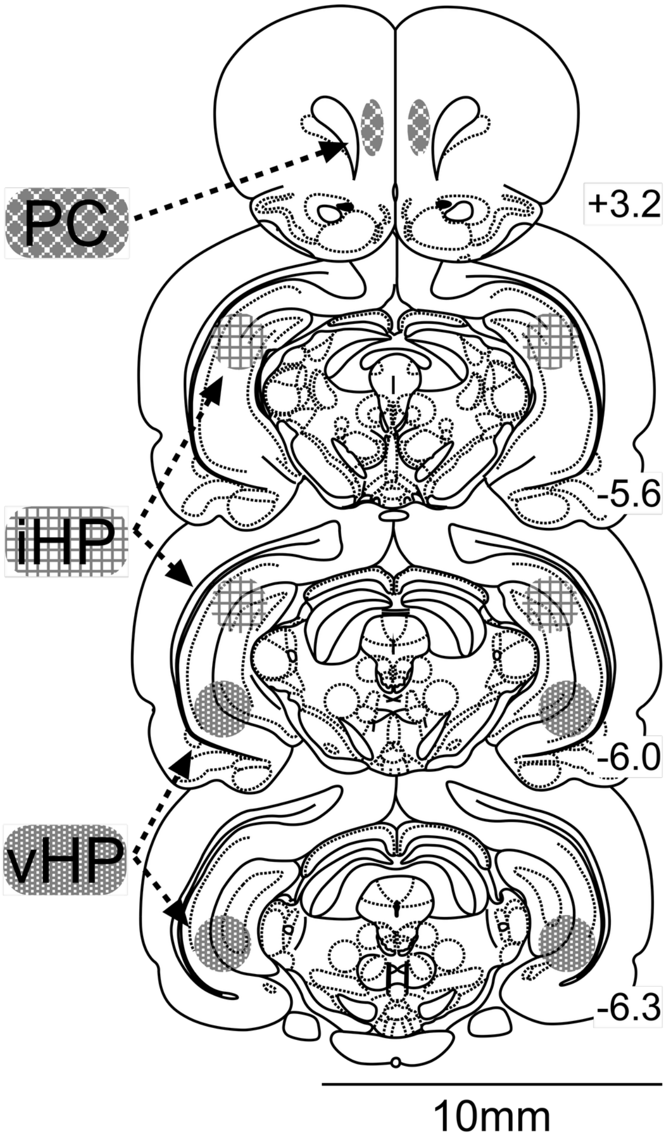

**Figure S1. Schematic of electrode placements for simultaneous bilateral recordings at six sites in the prefrontal cortex (PC), intermediate hippocampus (iHP), and ventral hippocampus (vHP).** Coronal brain sections show the corresponding electrode locations, with dashed arrows indicating the relationship between the labeled brain regions and their electrode locations (Paxinos & Watson, 1982).

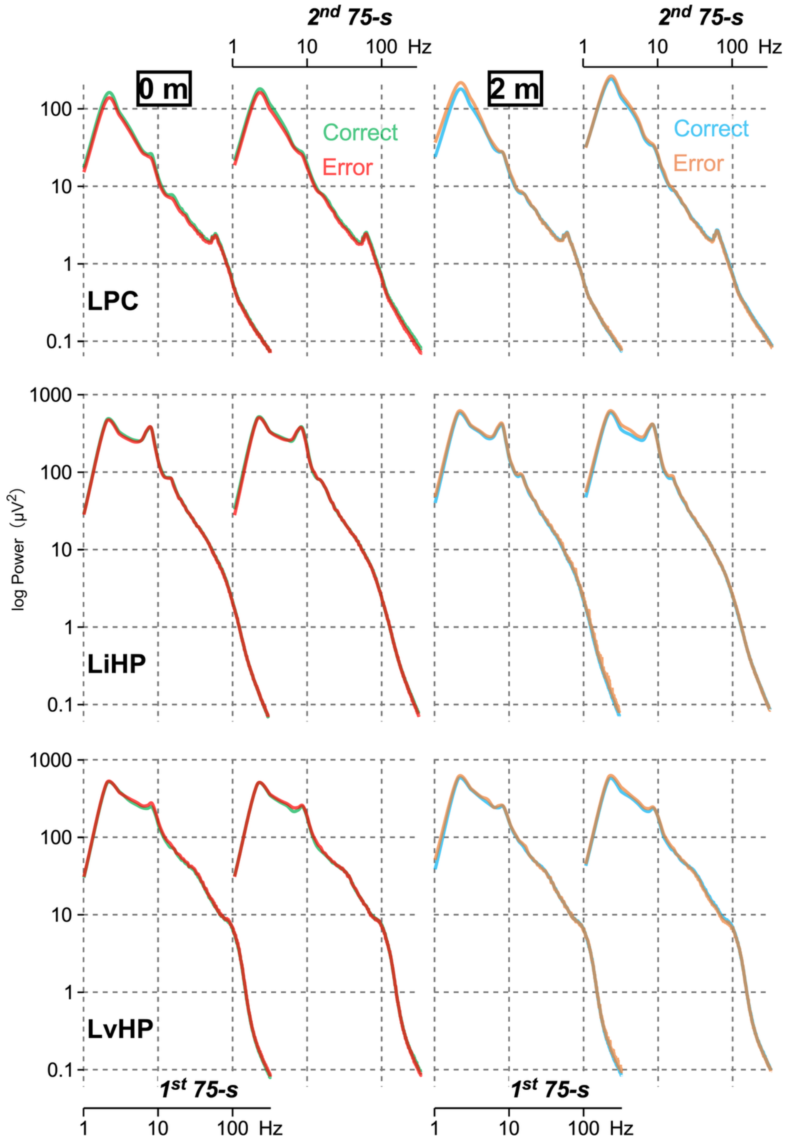

**Figure S2. Power spectral density (PSD) of left hemisphere local field potential (LFP; 1-300 Hz) during the 150-s delay (home cage) under 0-m and 2-m T-maze conditions.** Trials from the 75-s condition were excluded to avoid overlap with the 150-s delay analysis window. For visualization, the 150-s delay was divided into two 75-s periods (first and second halves). The two columns on the left correspond to the first and second 75-s periods of the 150-s delay under the 0-m condition, whereas the two columns on the right correspond to the first and second 75-s periods of the 150-s delay under the 2-m condition. Correct trials are shown in green (0 m) and cyan (2 m), and error trials are shown in red (0 m) and orange (2 m). The x-axis represents frequency (Hz, logarithmic scale), and the y-axis represents PSD (logarithmic scale). Recording regions in the left hemisphere were the medial prefrontal cortex (LPC), intermediate hippocampus (LiHP), and ventral hippocampus (LvHP).

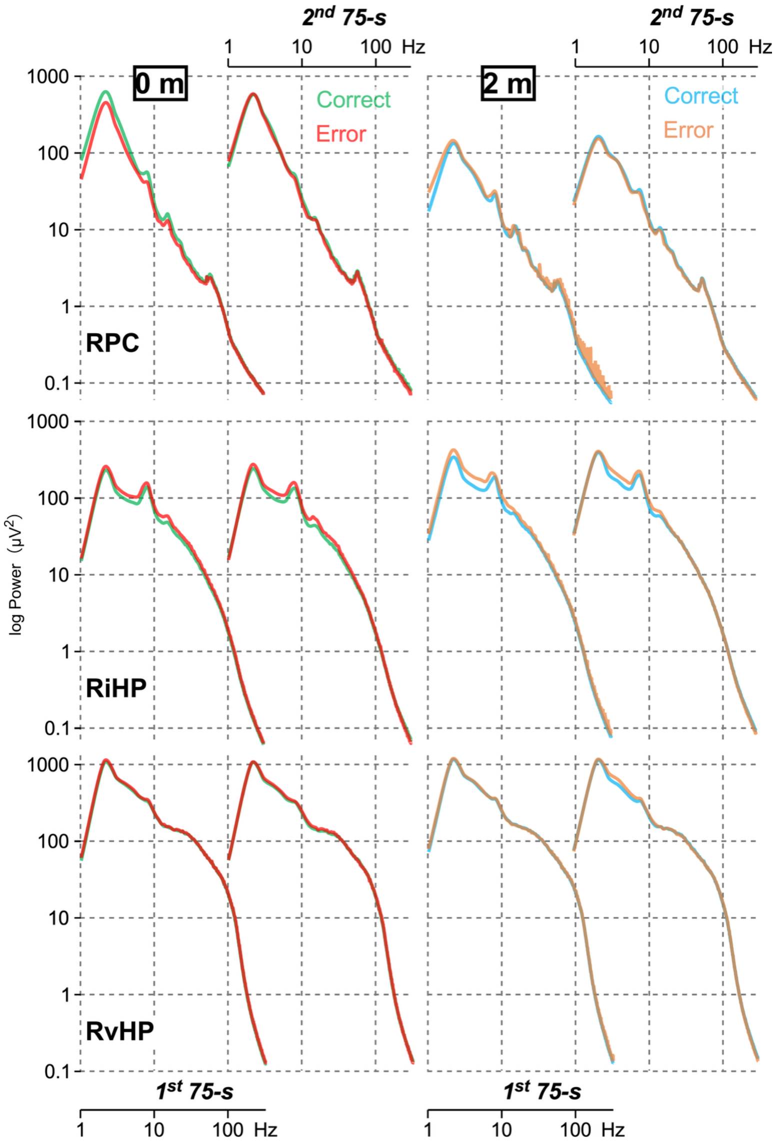

**Figure S3. PSD of right hemisphere LFP (1-300 Hz) during the 150-s delay (home cage) under 0-m and 2-m T-maze conditions.** The same inclusion criteria, period division, and trial color scheme as in Figure S2 were applied. Recording regions in the right hemisphere were RPC, RiHP, and RvHP, corresponding to the left hemisphere regions shown in Figure S2. Based on the observed spectral profiles, the 8-10 Hz and 51-100 Hz frequency ranges were interpreted as theta-range and high-gamma-range activity, respectively.

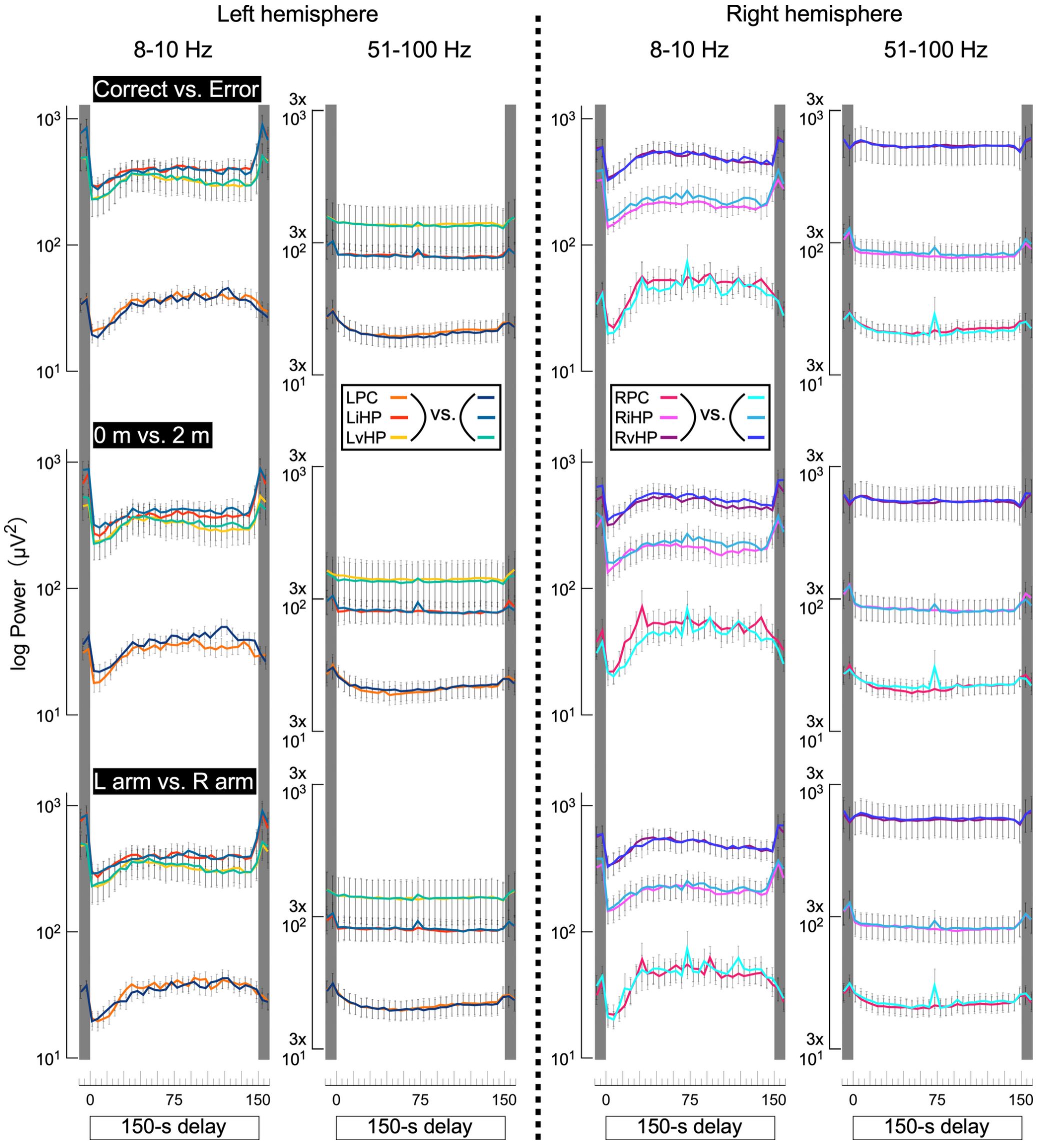

**Figure S4. Time course of log-transformed power in both hemispheres during the 150-s delay (home cage).** Power was analyzed in the 8-10 Hz and 51-100 Hz frequency bands across three experimental factors: trial outcome (correct vs. error), task distance (0 m vs. 2 m), and arm entry (L vs. R). Data are shown for each recording site (LPC/RPC, LiHP/RiHP, LvHP/RvHP; see insets for color coding). For statistical analysis, data were segmented into consecutive 5-s bins (15 bins total) and additionally grouped into two periods corresponding to the first and second 75-s periods (see Table S1). Consistent with the results of the repeated-measures ANOVA summarized in Table S1, no significant main effects or interactions of any factor were observed in either frequency band or region (F[1,40] = 0.00-0.68, p = 0.41-0.99), indicating stable power dynamics across conditions. Error bars represent mean ± SE.

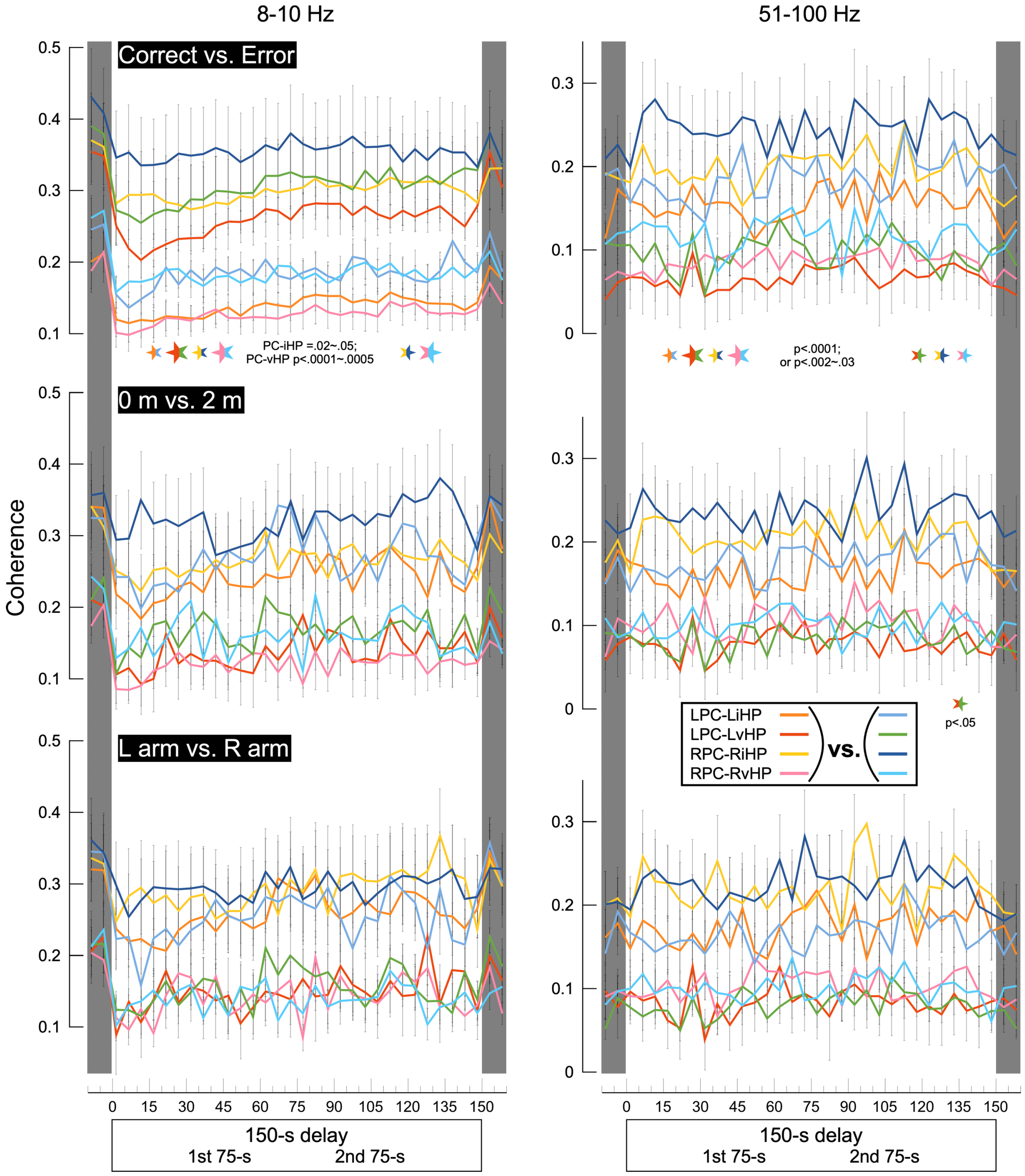

**Figure S5. Time course of hippocampal-prefrontal coherence during the 150-s home-cage recording.** Data are shown for each PC-iHP/vHP pair (e.g., LPC-LiHP, LPC-LvHP, RPC-RiHP, RPC-RvHP; see insets for color coding) across three experimental factors (trial outcome: correct vs. error; task distance: 0 m vs. 2 m; and arm entry: L vs. R). Statistical analysis (Table S2) incorporated consecutive 5-s bins and two periods corresponding to the first and second 75-s periods. Post hoc comparisons between the two levels of each factor were performed separately for the first and second 75-s periods using the Bonferroni-Dunn test. The size of the significance markers is the same as in Figure S4.

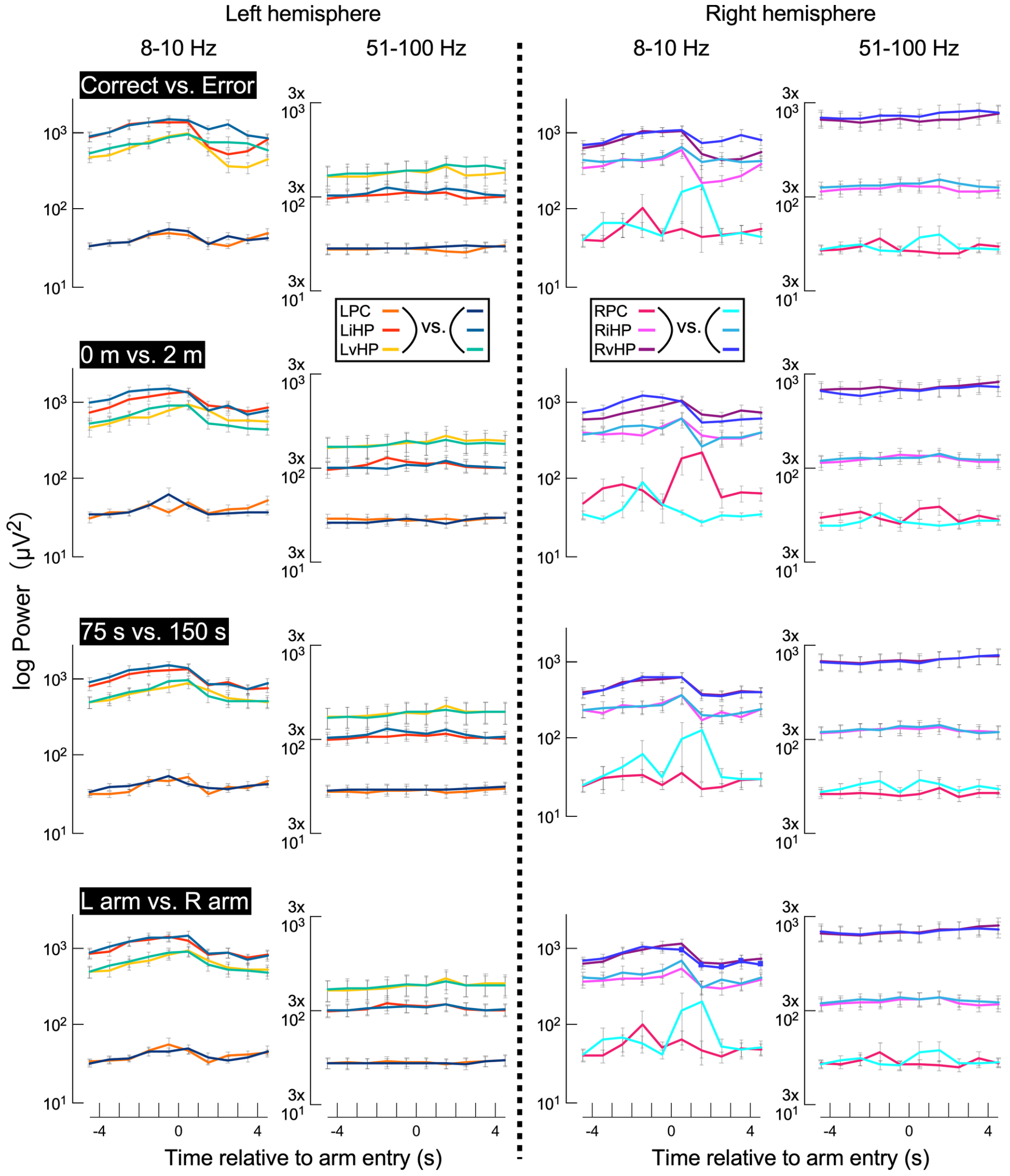

**Figure S6. Time course of log-transformed power in both hemispheres during the 10-s period surrounding arm entry.** Power was analyzed in the 8-10 Hz and 51-100 Hz frequency bands across four experimental factors: trial outcome (correct vs. error), task distance (0 m vs. 2 m), delay length (75 s vs. 150 s) and arm entry (L vs. R). Data are shown for each recording site (LPC/RPC, LiHP/RiHP, LvHP/RvHP; see insets for color coding). For statistical analysis, data were segmented into consecutive 1-s bins (5 bins total) and additionally grouped into two periods corresponding to the A1 (−5 to 0 s) and A2 (0 to +5 s) (see Table S4). Consistent with the results of the repeated-measures ANOVA summarized in Table S4, no significant main effects or interactions of any factor were observed in either frequency band or region ($F[1,80] = 0.00\text{-}2.01$, $p = 0.160\text{-}0.999$), indicating stable power dynamics across conditions. Error bars represent mean ± SE.

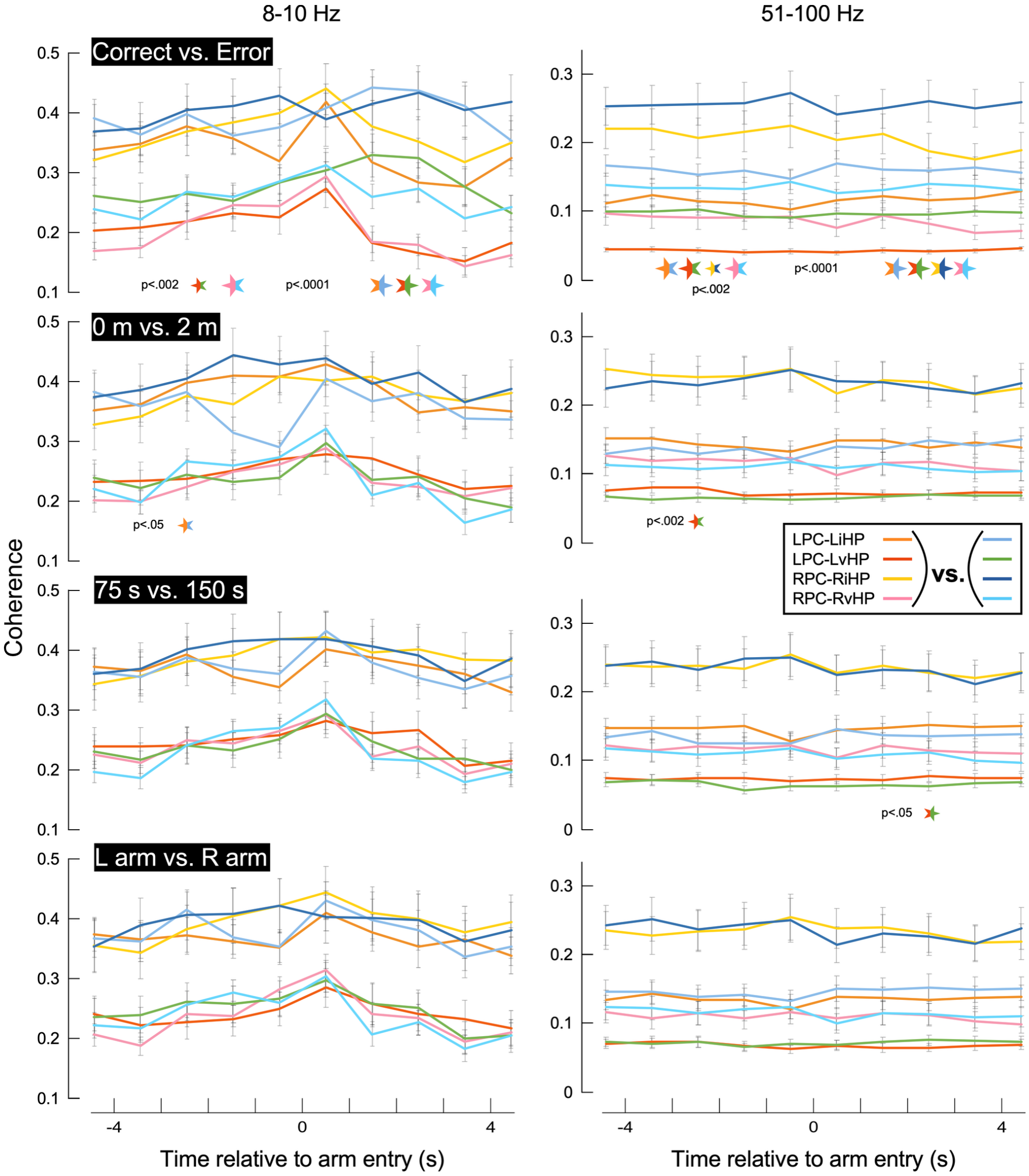

**Figure S7. Time course of coherence in the 8-10 Hz and 51-100 Hz frequency bands between PC and iHP/vHP in both hemispheres during the 10-s period surrounding arm entry.** Data are shown for each PC-iHP/vHP pair (e.g., LPC-LiHP, LPC-LvHP, RPC-RiHP, RPC-RvHP; see insets for color coding) across four experimental factors (trial outcome: correct vs. error; task distance: 0 m vs. 2 m; delay length: 75 s vs. 150 s; and arm entry: L vs. R). Statistical analysis (Table S5) incorporated consecutive 1-s bins and two periods corresponding to A1 (−5 to 0 s) and A2 (0 to +5 s). Post hoc comparisons between the two levels of each factor were performed separately for the A1 and A2 periods using the Bonferroni-Dunn test. The size of the significance markers is the same as in Figure S5.

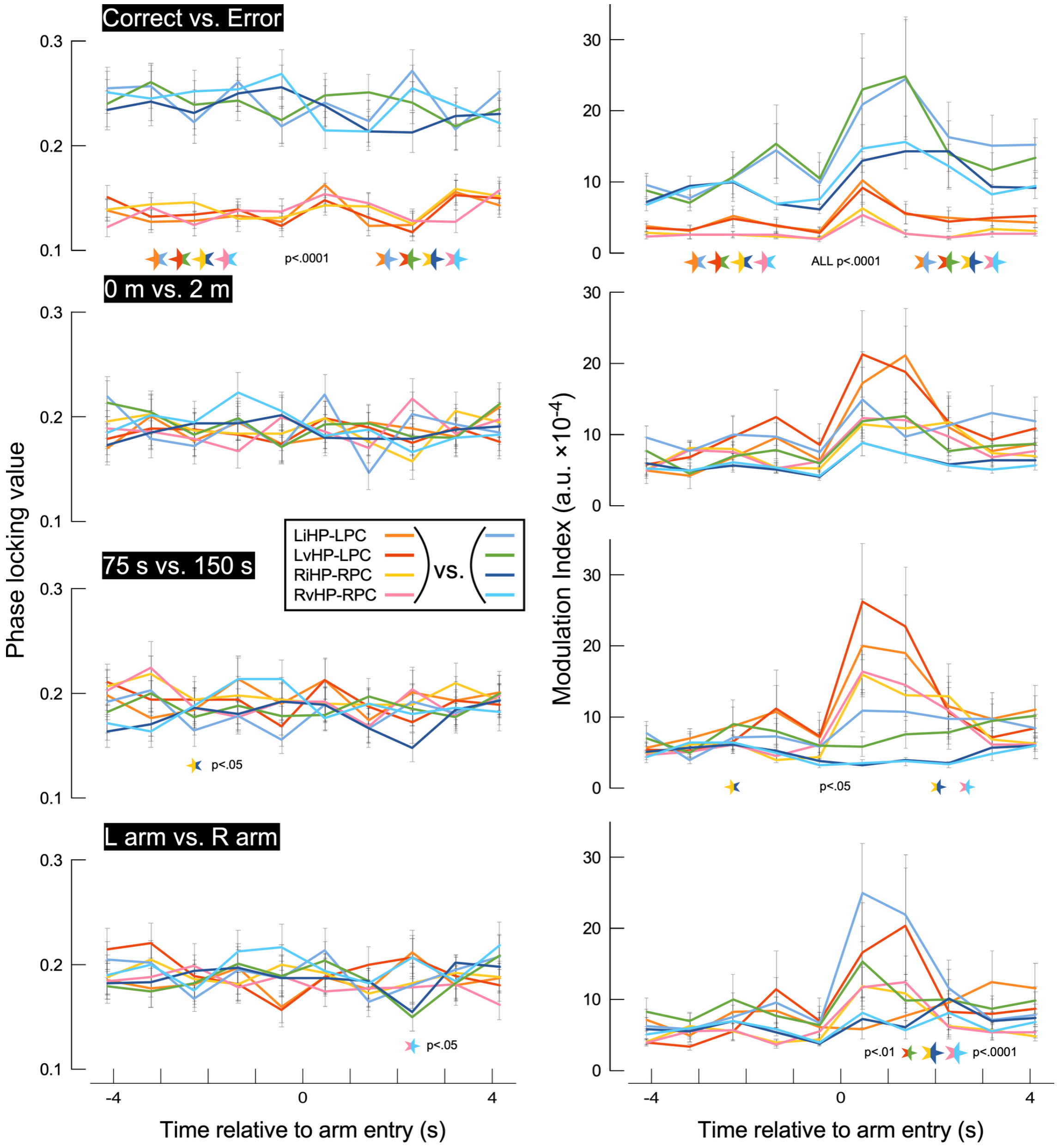

**Figure S8. Time course of hippocampal-prefrontal PLV and MI during the 10-s period surrounding arm entry.** Data are shown for PLV and MI between iHP/vHP (8-10 Hz) and PC (51-100 Hz) across four experimental factors (trial outcome: correct vs. error; task distance: 0 m vs. 2 m; delay length: 75 s vs. 150 s; and arm entry: L vs. R). Statistical analysis (Table S6) incorporated consecutive 1-s bins and two periods corresponding to A1 (−5 to 0 s) and A2 (0 to +5 s). Post hoc comparisons between the two levels of each factor were performed separately for the A1 and A2 periods using the Bonferroni-Dunn test. The size of the significance markers is the same as in Figure S7.

**Supplementary Tables S1-S6**

**Table S1. ANOVA table for spectral power in both hemispheres during a 150-s delay in the home cage.** Power in the 8-10 Hz and 51-100 Hz bands across brain regions was analyzed using a repeated-measures ANOVA. The model included four within-subject factors (accuracy [correct/error; C/E], distance [0 m/2 m], arm entry [L/R], and time [consecutive 5-s bins; 15 bins]) and one between-subject factor (period). In Tables S1-S3, period was defined as the first (D1) and second (D2) 75-s periods of the delay epoch. In Tables S4-S6, period was defined as the first (A1) and second (A2) 5-s periods around T-maze arm entry. Cells with p < 0.05 are highlighted in color in the final row. Colors indicate individual indices, and multiple colors denote interaction effects. When a within-subject factor shows a significant effect, main effects of time bins and period are not additionally highlighted by color, regardless of interaction effects. When a within-subject factor is not significant, significant main effects of time bins or period are indicated by color. The color-coding scheme in Tables S2 and S3 follows that used here. Abbreviations: LPC/RPC, left/right prefrontal cortex; LiHP/RiHP, left/right intermediate hippocampus; LvHP/RvHP, left/right ventral hippocampus.

| Power in home cage | df | | 8-10 Hz | | | 51-100 Hz | | | | 8-10 Hz | | | 51-100 Hz | | | | 8-10 Hz | | | 51-100 Hz | | |
|---|---|---|---|---|---|---|---|---|---|---|---|---|---|---|---|---|---|---|---|---|---|---|
| | | | F-value | p-value | Lambda | F-value | p-value | Lambda | | F-value | p-value | Lambda | F-value | p-value | Lambda | | F-value | p-value | Lambda | F-value | p-value | Lambda |
| C/E | 1 | | 0.032 | 0.8583 | 0.032 | 0.011 | 0.9152 | 0.011 | | 0.007 | 0.9359 | 0.007 | 0.001 | 0.9701 | 0.001 | | 0.002 | 0.9612 | 0.002 | 0.001 | 0.9745 | 0.001 |
| 0m/2m | 1 | | 0.679 | 0.4149 | 0.679 | 0.011 | 0.9176 | 0.011 | | 0.092 | 0.7628 | 0.092 | 0.003 | 0.9588 | 0.003 | | 0.007 | 0.9345 | 0.007 | 0.006 | 0.9367 | 0.006 |
| L/R | 1 | | 0.033 | 0.8566 | 0.033 | 0.003 | 0.9559 | 0.003 | | 2E-05 | 0.9962 | 2E-05 | 0.003 | 0.957 | 0.003 | | 0.0003 | 0.9873 | 0.0003 | 5E-05 | 0.9943 | 5E-05 |
| C/E x 0m/2m | 1 | | 0.039 | 0.8449 | 0.039 | 0.0001 | 0.9911 | 0.0001 | | 6E-06 | 0.998 | 6E-06 | 2E-06 | 0.9988 | 2E-06 | | 5E-13 | >.9999 | 5E-13 | 0.001 | 0.9783 | 0.001 |
| C/E x L/R | 1 | | 0.006 | 0.9407 | 0.006 | 0.002 | 0.9684 | 0.002 | | 0.0002 | 0.9882 | 0.0002 | 0.003 | 0.9563 | 0.003 | | 0.006 | 0.9404 | 0.006 | 0.0002 | 0.9896 | 0.0002 |
| 0m/2m x L/R | 1 | | 0.123 | 0.7276 | 0.123 | 0.005 | 0.9455 | 0.005 | | 0.002 | 0.9618 | 0.002 | 0.0004 | 0.9848 | 0.0004 | | 0.005 | 0.9414 | 0.005 | 8E-05 | 0.9931 | 8E-05 |
| C/E x 0m/2m x L/R | 1 | | 0.027 | 0.8695 | 0.027 | 0.001 | 0.9726 | 0.001 | | 0.01 | 0.9224 | 0.01 | 0.006 | 0.9409 | 0.006 | | 0.001 | 0.9744 | 0.001 | 2E-05 | 0.9965 | 2E-05 |
| Group | 40 | | | | | | | | | | | | | | | | | | | | | |
| D1/D2 | 1 | | 66.569 | <.0001 | 66.569 | 1.989 | 0.1662 | 1.989 | | 24.544 | <.0001 | 24.544 | 8.786 | 0.0051 | 8.786 | | 0.014 | 0.9053 | 0.014 | 3.027 | 0.0896 | 3.027 |
| D1/D2 x C/E | 1 | | 0.063 | 0.8036 | 0.063 | 0.367 | 0.5481 | 0.367 | | 0.049 | 0.8262 | 0.049 | 0.203 | 0.6545 | 0.203 | | 0.035 | 0.8525 | 0.035 | 8.657 | 0.0054 | 8.657 |
| D1/D2 x 0m/2m | 1 | | 4.703 | 0.0361 | 4.703 | 0.16 | 0.6909 | 0.16 | | 0.058 | 0.8111 | 0.058 | 1.618 | 0.2108 | 1.618 | | 1.341 | 0.2537 | 1.341 | 0.004 | 0.9497 | 0.004 |
| D1/D2 x L/R | 1 | | 0.131 | 0.719 | 0.131 | 0.442 | 0.5098 | 0.442 | | 0.944 | 0.337 | 0.944 | 0.037 | 0.848 | 0.037 | | 0.001 | 0.9816 | 0.001 | 0.113 | 0.7381 | 0.113 |
| D1/D2 x C/E x 0m/2m | 1 | | 0.0001 | 0.9907 | 0.0001 | 0.07 | 0.7922 | 0.07 | | 0.207 | 0.6515 | 0.207 | 0.021 | 0.8867 | 0.021 | | 0.309 | 0.5812 | 0.309 | 0.036 | 0.8514 | 0.036 |
| D1/D2 x C/E x L/R | 1 | | 0.147 | 0.7039 | 0.147 | 0.085 | 0.7724 | 0.085 | | 0.308 | 0.5821 | 0.308 | 0.3 | 0.5867 | 0.3 | | 0.115 | 0.7358 | 0.115 | 0.945 | 0.3368 | 0.945 |
| D1/D2 x 0m/2m x L/R | 1 | | 0.037 | 0.8477 | 0.037 | 0.863 | 0.3586 | 0.863 | | 0.058 | 0.8105 | 0.058 | 1.366 | 0.2494 | 1.366 | | 0.481 | 0.4921 | 0.481 | 0.149 | 0.7015 | 0.149 |
| D1/D2 x C/E x 0m/2m x L/R | 1 | | 0.061 | 0.8063 | 0.061 | 0.011 | 0.9183 | 0.011 | | 0.669 | 0.4182 | 0.669 | 1.44 | 0.2372 | 1.44 | | 0.065 | 0.7998 | 0.065 | 0.542 | 0.4659 | 0.542 |
| D1/D2 x Group | 40 | | | | | | | | | | | | | | | | | | | | | |
| 15bins | 14 | LPC | 10.63 | <.0001 | 148.82 | 15.127 | <.0001 | 211.77 | LiHP | 10.204 | <.0001 | 142.86 | 2.533 | 0.0016 | 35.457 | LvHP | 9.034 | <.0001 | 126.48 | 1.512 | 0.1016 | 21.168 |
| 15bins x C/E | 14 | | 0.732 | 0.743 | 10.243 | 0.33 | 0.9902 | 4.616 | | 0.249 | 0.9978 | 3.484 | 0.31 | 0.9927 | 4.347 | | 0.357 | 0.9853 | 5.003 | 0.815 | 0.6536 | 11.403 |
| 15bins x 0m/2m | 14 | | 1.496 | 0.1073 | 20.947 | 1.124 | 0.3326 | 15.742 | | 0.552 | 0.9018 | 7.724 | 1.217 | 0.258 | 17.041 | | 0.573 | 0.8864 | 8.028 | 1.197 | 0.2734 | 16.754 |
| 15bins x L/R | 14 | | 0.86 | 0.6027 | 12.045 | 0.482 | 0.943 | 6.744 | | 0.647 | 0.8256 | 9.061 | 0.584 | 0.8785 | 8.175 | | 0.656 | 0.8171 | 9.191 | 1.296 | 0.2049 | 18.14 |
| 15bins x C/E x 0m/2m | 14 | | 0.569 | 0.8895 | 7.968 | 0.319 | 0.9917 | 4.462 | | 0.218 | 0.9989 | 3.054 | 0.259 | 0.9972 | 3.619 | | 0.652 | 0.821 | 9.132 | 0.304 | 0.9935 | 4.257 |
| 15bins x C/E x L/R | 14 | | 0.718 | 0.7572 | 10.051 | 0.15 | 0.9999 | 2.095 | | 0.604 | 0.863 | 8.451 | 1.255 | 0.2314 | 17.567 | | 0.332 | 0.9898 | 4.647 | 0.572 | 0.8877 | 8.003 |
| 15bins x 0m/2m x L/R | 14 | | 1.392 | 0.1514 | 19.493 | 0.478 | 0.9448 | 6.693 | | 0.539 | 0.91 | 7.552 | 0.399 | 0.9752 | 5.588 | | 0.476 | 0.9459 | 6.663 | 1.034 | 0.417 | 14.478 |
| 15bins x C/E x 0m/2m x L/R | 14 | | 0.6 | 0.8657 | 8.405 | 0.366 | 0.9835 | 5.124 | | 0.594 | 0.8705 | 8.319 | 1.295 | 0.205 | 18.137 | | 0.915 | 0.5425 | 12.807 | 0.846 | 0.6191 | 11.839 |
| 15bins x Group | 560 | | | | | | | | | | | | | | | | | | | | | |
| D1/D2 x 15bins | 14 | | 12.378 | <.0001 | 173.29 | 20.9 | <.0001 | 292.61 | | 7.4 | <.0001 | 103.6 | 0.378 | 0.9808 | 5.289 | | 14.888 | <.0001 | 208.44 | 4.61 | <.0001 | 64.536 |
| D1/D2 x 15bins x C/E | 14 | | 1.048 | 0.4038 | 14.666 | 0.191 | 0.9995 | 2.679 | | 0.39 | 0.9778 | 5.454 | 0.364 | 0.9839 | 5.098 | | 0.442 | 0.9605 | 6.192 | 0.779 | 0.6926 | 10.905 |
| D1/D2 x 15bins x 0m/2m | 14 | | 1.356 | 0.1699 | 18.988 | 0.504 | 0.9314 | 7.054 | | 0.712 | 0.7634 | 9.966 | 0.775 | 0.6968 | 10.851 | | 0.311 | 0.9927 | 4.356 | 0.87 | 0.5918 | 12.183 |
| D1/D2 x 15bins x L/R | 14 | | 1.563 | 0.0852 | 21.88 | 0.15 | 0.9999 | 2.097 | | 0.344 | 0.9879 | 4.813 | 0.563 | 0.894 | 7.88 | | 0.389 | 0.9779 | 5.449 | 1.121 | 0.3354 | 15.697 |
| D1/D2 x 15bins x C/E x 0m/2m | 14 | | 0.656 | 0.8175 | 9.185 | 0.154 | 0.9999 | 2.159 | | 0.198 | 0.9994 | 2.767 | 0.331 | 0.99 | 4.634 | | 0.376 | 0.9813 | 5.261 | 0.624 | 0.8464 | 8.731 |
| D1/D2 x 15bins x C/E x L/R | 14 | | 0.934 | 0.5219 | 13.071 | 0.275 | 0.9962 | 3.849 | | 0.26 | 0.9972 | 3.633 | 0.883 | 0.5771 | 12.368 | | 0.391 | 0.9775 | 5.474 | 1.159 | 0.3035 | 16.223 |
| D1/D2 x 15bins x 0m/2m x L/R | 14 | | 0.847 | 0.6181 | 11.852 | 0.277 | 0.996 | 3.879 | | 0.469 | 0.949 | 6.57 | 0.232 | 0.9985 | 3.25 | | 0.306 | 0.9932 | 4.288 | 1.308 | 0.1976 | 18.307 |
| D1/D2 x 15bins x C/E x 0m/2m x L/R | 14 | | 1.011 | 0.4402 | 14.155 | 0.112 | >.9999 | 1.571 | | 0.407 | 0.9729 | 5.694 | 0.5 | 0.9333 | 7.006 | | 0.253 | 0.9976 | 3.537 | 0.901 | 0.5575 | 12.616 |
| D1/D2 x 15bins x Group | 560 | | | | | | | | | | | | | | | | | | | | | |
| C/E | 1 | | 0.079 | 0.7803 | 0.079 | 0.011 | 0.9169 | 0.011 | | 0.113 | 0.7386 | 0.113 | 0.02 | 0.8894 | 0.02 | | 0.002 | 0.9684 | 0.002 | 3E-05 | 0.9956 | 3E-05 |
| 0m/2m | 1 | | 0.2 | 0.657 | 0.2 | 0.032 | 0.8586 | 0.032 | | 0.083 | 0.7742 | 0.083 | 0.0001 | 0.9915 | 0.0001 | | 0.075 | 0.786 | 0.075 | 0.001 | 0.9747 | 0.001 |
| L/R | 1 | | 0.025 | 0.8741 | 0.025 | 0.044 | 0.8344 | 0.044 | | 0.023 | 0.8807 | 0.023 | 0.004 | 0.9482 | 0.004 | | 0.001 | 0.9755 | 0.001 | 0.003 | 0.957 | 0.003 |
| C/E x 0m/2m | 1 | | 0.019 | 0.8921 | 0.019 | 0.014 | 0.905 | 0.014 | | 0.002 | 0.9666 | 0.002 | 0.012 | 0.912 | 0.012 | | 2E-05 | 0.9962 | 2E-05 | 0.0002 | 0.9897 | 0.0002 |
| C/E x L/R | 1 | | 0.007 | 0.9357 | 0.007 | 0.009 | 0.9258 | 0.009 | | 0.007 | 0.9326 | 0.007 | 0.001 | 0.9729 | 0.001 | | 0.004 | 0.948 | 0.004 | 0.0002 | 0.9878 | 0.0002 |
| 0m/2m x L/R | 1 | | 0.04 | 0.8419 | 0.04 | 0.004 | 0.9486 | 0.004 | | 0.002 | 0.9665 | 0.002 | 0.0004 | 0.9846 | 0.0004 | | 0.028 | 0.8689 | 0.028 | 0.001 | 0.9759 | 0.001 |
| C/E x 0m/2m x L/R | 1 | | 0.024 | 0.8776 | 0.024 | 0.006 | 0.9372 | 0.006 | | 0.013 | 0.9082 | 0.013 | 0.001 | 0.9725 | 0.001 | | 2E-06 | 0.9989 | 2E-06 | 6E-07 | 0.9994 | 6E-07 |
| Group | 40 | | | | | | | | | | | | | | | | | | | | | |
| D1/D2 | 1 | | 10.026 | 0.003 | 10.026 | 0.018 | 0.8935 | 0.018 | | 6.827 | 0.0126 | 6.827 | 33.762 | <.0001 | 33.762 | | 0.044 | 0.8352 | 0.044 | 18.111 | 0.0001 | 18.111 |
| D1/D2 x C/E | 1 | | 0.11 | 0.7421 | 0.11 | 0.798 | 0.3771 | 0.798 | | 0.056 | 0.8134 | 0.056 | 0.883 | 0.3529 | 0.883 | | 0.119 | 0.7323 | 0.119 | 4.217 | 0.0466 | 4.217 |
| D1/D2 x 0m/2m | 1 | | 1.229 | 0.2742 | 1.229 | 1.001 | 0.3231 | 1.001 | | 2.393 | 0.1298 | 2.393 | 0.385 | 0.5385 | 0.385 | | 0.099 | 0.7542 | 0.099 | 0.517 | 0.4765 | 0.517 |
| D1/D2 x L/R | 1 | | 0.333 | 0.5674 | 0.333 | 0.218 | 0.643 | 0.218 | | 0.669 | 0.4182 | 0.669 | 0.167 | 0.6848 | 0.167 | | 0.208 | 0.6506 | 0.208 | 0.009 | 0.9248 | 0.009 |
| D1/D2 x C/E x 0m/2m | 1 | | 0.462 | 0.5006 | 0.462 | 0.186 | 0.6683 | 0.186 | | 0.028 | 0.8672 | 0.028 | 0.036 | 0.8506 | 0.036 | | 0.014 | 0.9048 | 0.014 | 0.189 | 0.666 | 0.189 |
| D1/D2 x C/E x L/R | 1 | | 0.162 | 0.6895 | 0.162 | 0.2 | 0.6568 | 0.2 | | 0.025 | 0.8759 | 0.025 | 0.99 | 0.3258 | 0.99 | | 0.0002 | 0.9893 | 0.0002 | 0.1 | 0.7534 | 0.1 |
| D1/D2 x 0m/2m x L/R | 1 | | 2.513 | 0.1208 | 2.513 | 0.009 | 0.9232 | 0.009 | | 0.113 | 0.7388 | 0.113 | 1.526 | 0.2239 | 1.526 | | 0.263 | 0.6108 | 0.263 | 0.343 | 0.5612 | 0.343 |
| D1/D2 x C/E x 0m/2m x L/R | 1 | | 0.813 | 0.3728 | 0.813 | 0.394 | 0.5339 | 0.394 | | 0.02 | 0.8896 | 0.02 | 1.356 | 0.2511 | 1.356 | | 0.044 | 0.8354 | 0.044 | 1.003 | 0.3226 | 1.003 |
| D1/D2 x Group | 40 | | | | | | | | | | | | | | | | | | | | | |
| 15bins | 14 | RPC | 4.382 | <.0001 | 61.347 | 3.31 | <.0001 | 46.341 | RiHP | 12.58 | <.0001 | 176.13 | 5.365 | <.0001 | 75.111 | RvHP | 10.757 | <.0001 | 150.6 | 6.399 | <.0001 | 89.587 |
| 15bins x C/E | 14 | | 0.576 | 0.8843 | 8.066 | 0.705 | 0.7704 | 9.868 | | 0.416 | 0.97 | 5.822 | 0.544 | 0.9071 | 7.613 | | 0.939 | 0.516 | 13.148 | 0.344 | 0.9878 | 4.821 |
| 15bins x 0m/2m | 14 | | 1.014 | 0.4377 | 14.19 | 0.868 | 0.5947 | 12.146 | | 0.515 | 0.9249 | 7.214 | 1.158 | 0.3039 | 16.217 | | 0.879 | 0.5817 | 12.31 | 1.556 | 0.0871 | 21.789 |
| 15bins x L/R | 14 | | 0.844 | 0.6204 | 11.823 | 0.819 | 0.6483 | 11.471 | | 0.556 | 0.8987 | 7.788 | 0.754 | 0.7197 | 10.553 | | 0.574 | 0.8862 | 8.032 | 0.323 | 0.9912 | 4.517 |
| 15bins x C/E x 0m/2m | 14 | | 0.57 | 0.8891 | 7.976 | 0.721 | 0.7536 | 10.1 | | 0.398 | 0.9756 | 5.566 | 0.638 | 0.8336 | 8.936 | | 0.567 | 0.8913 | 7.934 | 0.476 | 0.9457 | 6.667 |
| 15bins x C/E x L/R | 14 | | 1.058 | 0.3933 | 14.817 | 0.847 | 0.6174 | 11.86 | | 0.223 | 0.9988 | 3.127 | 0.965 | 0.488 | 13.513 | | 0.656 | 0.8171 | 9.191 | 0.241 | 0.9981 | 3.371 |
| 15bins x 0m/2m x L/R | 14 | | 1.216 | 0.2589 | 17.023 | 0.959 | 0.4941 | 13.432 | | 0.671 | 0.8039 | 9.389 | 0.722 | 0.7529 | 10.11 | | 0.454 | 0.956 | 6.349 | 0.579 | 0.8821 | 8.109 |
| 15bins x C/E x 0m/2m x L/R | 14 | | 0.922 | 0.5347 | 12.907 | 1.193 | 0.2762 | 16.702 | | 0.583 | 0.8793 | 8.161 | 0.977 | 0.4756 | 13.677 | | 0.645 | 0.8275 | 9.031 | 0.441 | 0.961 | 6.175 |
| 15bins x Group | 560 | | | | | | | | | | | | | | | | | | | | | |
| D1/D2 x 15bins | 14 | | 7.358 | <.0001 | 103.02 | 3.117 | 0.0001 | 43.638 | | 8.723 | <.0001 | 122.13 | 2.85 | 0.0004 | 39.906 | | 26.466 | <.0001 | 370.53 | 5.827 | <.0001 | 81.573 |
| D1/D2 x 15bins x C/E | 14 | | 0.48 | 0.9437 | 6.725 | 0.617 | 0.8522 | 8.635 | | 0.537 | 0.9116 | 7.517 | 0.628 | 0.843 | 8.786 | | 0.288 | 0.9951 | 4.029 | 0.574 | 0.8857 | 8.04 |
| D1/D2 x 15bins x 0m/2m | 14 | | 1.022 | 0.4292 | 14.307 | 1.047 | 0.4043 | 14.659 | | 1.211 | 0.2628 | 16.95 | 0.318 | 0.9918 | 4.448 | | 0.442 | 0.9606 | 6.191 | 0.514 | 0.9257 | 7.194 |
| D1/D2 x 15bins x L/R | 14 | | 0.589 | 0.8749 | 8.242 | 0.769 | 0.7033 | 10.768 | | 0.154 | 0.9999 | 2.153 | 0.821 | 0.646 | 11.5 | | 0.474 | 0.9466 | 6.642 | 0.386 | 0.9787 | 5.409 |
| D1/D2 x 15bins x C/E x 0m/2m | 14 | | 0.851 | 0.6127 | 11.919 | 0.866 | 0.5965 | 12.123 | | 0.468 | 0.9497 | 6.548 | 0.428 | 0.9658 | 5.996 | | 0.239 | 0.9982 | 3.34 | 0.506 | 0.9302 | 7.083 |
| D1/D2 x 15bins x C/E x L/R | 14 | | 0.682 | 0.7928 | 9.551 | 1.19 | 0.2786 | 16.66 | | 0.293 | 0.9946 | 4.106 | 0.649 | 0.8241 | 9.084 | | 0.29 | 0.9949 | 4.062 | 0.551 | 0.902 | 7.719 |
| D1/D2 x 15bins x 0m/2m x L/R | 14 | | 0.554 | 0.9 | 7.76 | 0.793 | 0.6769 | 11.107 | | 0.487 | 0.9403 | 6.821 | 0.597 | 0.8681 | 8.362 | | 0.594 | 0.871 | 8.312 | 0.267 | 0.9967 | 3.737 |
| D1/D2 x 15bins x C/E x 0m/2m x L/R | 14 | | 0.46 | 0.953 | 6.444 | 0.94 | 0.515 | 13.161 | | 0.422 | 0.9679 | 5.91 | 0.512 | 0.9266 | 7.173 | | 0.193 | 0.9995 | 2.695 | 0.215 | 0.999 | 3.007 |
| D1/D2 x 15bins x Group | 560 | | | | | | | | | | | | | | | | | | | | | |

Cells are shaded for p < 0.05: CE | 0m2m | L/R | bins/period

**Table S2. ANOVA table of coherence between PC and iHP/vHP in both hemispheres during a 150-s delay in the home cage.** Coherence between PC and iHP/vHP in the 8-10 Hz and 51-100 Hz bands was analyzed using the same repeated-measures ANOVA as in Table S1. See Table S1 for details on factors, significance criteria, and abbreviations.

| Coherence in home cage | df | | 8-10 Hz | | | 51-100 Hz | | | | 8-10 Hz | | | 51-100 Hz | | |
|---|---|---|---|---|---|---|---|---|---|---|---|---|---|---|---|
| | | | F-value | p-value | Lam-bda | F-value | p-value | Lam-bda | | F-value | p-value | Lam-bda | F-value | p-value | Lam-bda |
| C/E | 1 | LPC_LiHP | 0.161 | 0.6903 | 0.161 | 0.313 | 0.5791 | 0.313 | RPC_RIHP | 0.425 | 0.518 | 0.425 | 0.537 | 0.4679 | 0.537 |
| 0m/2m | 1 | | 0.002 | 0.9609 | 0.002 | 0.076 | 0.7838 | 0.076 | | 0.15 | 0.701 | 0.15 | 0.222 | 0.6404 | 0.222 |
| L/R | 1 | | 0.006 | 0.9368 | 0.006 | 0.028 | 0.8688 | 0.028 | | 0.001 | 0.9765 | 0.001 | 0.007 | 0.9348 | 0.007 |
| C/E x 0m/2m | 1 | | 0.002 | 0.9679 | 0.002 | 0.049 | 0.8261 | 0.049 | | 0.004 | 0.9528 | 0.004 | 0.009 | 0.9267 | 0.009 |
| C/E x L/R | 1 | | 0.004 | 0.9511 | 0.004 | 0.048 | 0.827 | 0.048 | | 0.008 | 0.9294 | 0.008 | 0.012 | 0.9131 | 0.012 |
| 0m/2m x L/R | 1 | | 0.176 | 0.6774 | 0.176 | 0.008 | 0.9309 | 0.008 | | 0.003 | 0.9575 | 0.003 | 0.01 | 0.9217 | 0.01 |
| C/E x 0m/2m x L/R | 1 | | 0.043 | 0.8374 | 0.043 | 0.008 | 0.9303 | 0.008 | | 3E-05 | 0.9953 | 3E-05 | 0.01 | 0.9226 | 0.01 |
| Group | 40 | | | | | | | | | | | | | | |
| D1/D2 | 1 | | 10.541 | 0.0024 | 10.541 | 7.877 | 0.0077 | 7.877 | | 5.643 | 0.0224 | 5.643 | 1.433 | 0.2383 | 1.433 |
| D1/D2 x C/E | 1 | | 1.469 | 0.2327 | 1.469 | 0.03 | 0.8629 | 0.03 | | 0.59 | 0.447 | 0.59 | 0.503 | 0.4823 | 0.503 |
| D1/D2 x 0m/2m | 1 | | 0.396 | 0.5326 | 0.396 | 0.041 | 0.84 | 0.041 | | 1.113 | 0.2977 | 1.113 | 0.199 | 0.6582 | 0.199 |
| D1/D2 x L/R | 1 | | 0.385 | 0.5387 | 0.385 | 0.0004 | 0.9833 | 0.0004 | | 2.306 | 0.1367 | 2.306 | 0.148 | 0.7029 | 0.148 |
| D1/D2 x C/E x 0m/2m | 1 | | 1.047 | 0.3124 | 1.047 | 0.013 | 0.9089 | 0.013 | | 3.547 | 0.067 | 3.547 | 0.015 | 0.9029 | 0.015 |
| D1/D2 x C/E x L/R | 1 | | 0.172 | 0.6807 | 0.172 | 0.164 | 0.6874 | 0.164 | | 0.439 | 0.5112 | 0.439 | 2.252 | 0.1413 | 2.252 |
| D1/D2 x 0m/2m x L/R | 1 | | 8.129 | 0.0069 | 8.129 | 0.87 | 0.3565 | 0.87 | | 0.999 | 0.3235 | 0.999 | 1.677 | 0.2027 | 1.677 |
| D1/D2 x C/E x 0m/2m x L/R | 1 | | 0.926 | 0.3418 | 0.926 | 4.656 | 0.037 | 4.656 | | 0.846 | 0.3631 | 0.846 | 2.788 | 0.1028 | 2.788 |
| D1/D2 x Group | 40 | | | | | | | | | | | | | | |
| 15bins | 14 | | 1.994 | 0.0164 | 27.916 | 1.041 | 0.4106 | 14.569 | | 0.745 | 0.7291 | 10.429 | 1.038 | 0.4132 | 14.532 |
| 15bins x C/E | 14 | | 1.297 | 0.2039 | 18.162 | 1.496 | 0.1075 | 20.939 | | 0.594 | 0.8708 | 8.314 | 1.074 | 0.3781 | 15.042 |
| 15bins x 0m/2m | 14 | | 1.146 | 0.3144 | 16.04 | 0.762 | 0.7109 | 10.668 | | 0.885 | 0.575 | 12.395 | 0.833 | 0.6328 | 11.666 |
| 15bins x L/R | 14 | | 0.789 | 0.6812 | 11.053 | 1.285 | 0.2116 | 17.989 | | 0.648 | 0.8246 | 9.077 | 1.062 | 0.3898 | 14.868 |
| 15bins x C/E x 0m/2m | 14 | | 1.314 | 0.1939 | 18.392 | 1.246 | 0.2371 | 17.451 | | 1.567 | 0.0838 | 21.945 | 1.429 | 0.1346 | 19.999 |
| 15bins x C/E x L/R | 14 | | 0.826 | 0.6414 | 11.558 | 0.525 | 0.9188 | 7.355 | | 1.051 | 0.4 | 14.72 | 1.355 | 0.1706 | 18.97 |
| 15bins x 0m/2m x L/R | 14 | | 1.168 | 0.2958 | 16.356 | 0.543 | 0.9079 | 7.596 | | 0.648 | 0.8249 | 9.072 | 0.758 | 0.715 | 10.615 |
| 15bins x C/E x 0m/2m x L/R | 14 | | 2.001 | 0.016 | 28.02 | 0.918 | 0.5384 | 12.859 | | 1.084 | 0.3689 | 15.178 | 1.083 | 0.3705 | 15.155 |
| 15bins x Group | 560 | | | | | | | | | | | | | | |
| D1/D2 x 15bins | 14 | | 2.885 | 0.0003 | 40.397 | 0.884 | 0.5764 | 12.377 | | 1.38 | 0.1575 | 19.321 | 1.239 | 0.2421 | 17.35 |
| D1/D2 x 15bins x C/E | 14 | | 0.939 | 0.5156 | 13.152 | 0.806 | 0.6625 | 11.29 | | 1.031 | 0.4199 | 14.437 | 0.405 | 0.9734 | 5.672 |
| D1/D2 x 15bins x 0m/2m | 14 | | 1.273 | 0.2193 | 17.821 | 0.709 | 0.7667 | 9.92 | | 0.846 | 0.6189 | 11.842 | 0.771 | 0.7011 | 10.795 |
| D1/D2 x 15bins x L/R | 14 | | 1.171 | 0.2937 | 16.393 | 0.746 | 0.7284 | 10.439 | | 0.726 | 0.7489 | 10.164 | 2.106 | 0.0103 | 29.478 |
| D1/D2 x 15bins x C/E x 0m/2m | 14 | | 0.986 | 0.4663 | 13.8 | 0.494 | 0.9369 | 6.913 | | 0.865 | 0.5981 | 12.103 | 1.865 | 0.0275 | 26.116 |
| D1/D2 x 15bins x C/E x L/R | 14 | | 1.094 | 0.3596 | 15.321 | 1.37 | 0.1624 | 19.187 | | 0.595 | 0.8699 | 8.331 | 1.113 | 0.3424 | 15.587 |
| D1/D2 x 15bins x 0m/2m x L/R | 14 | | 0.763 | 0.7097 | 10.684 | 0.828 | 0.6385 | 11.595 | | 0.65 | 0.8234 | 9.095 | 0.626 | 0.8441 | 8.768 |
| D1/D2 x 15bins x C/E x 0m/2m x L/R | 14 | | 0.883 | 0.5771 | 12.369 | 0.464 | 0.9513 | 6.5 | | 0.929 | 0.5274 | 13 | 1.459 | 0.1217 | 20.424 |
| D1/D2 x 15bins x Group | 560 | | | | | | | | | | | | | | |
| C/E | 1 | LPC_LvHP | 1.359 | 0.2507 | 1.359 | 4.469 | 0.0408 | 4.469 | RPC_RvHP | 1.485 | 0.2301 | 1.485 | 1.346 | 0.2529 | 1.346 |
| 0m/2m | 1 | | 0.002 | 0.9667 | 0.002 | 0.266 | 0.6086 | 0.266 | | 0.004 | 0.952 | 0.004 | 0.014 | 0.905 | 0.014 |
| L/R | 1 | | 0.068 | 0.7959 | 0.068 | 0.052 | 0.8214 | 0.052 | | 0.008 | 0.9279 | 0.008 | 0.016 | 0.9015 | 0.016 |
| C/E x 0m/2m | 1 | | 0.12 | 0.7305 | 0.12 | 1.609 | 0.2119 | 1.609 | | 0.037 | 0.8493 | 0.037 | 0.001 | 0.9786 | 0.001 |
| C/E x L/R | 1 | | 1E-05 | 0.9972 | 1E-05 | 0.106 | 0.746 | 0.106 | | 0.0005 | 0.983 | 0.0005 | 0.009 | 0.9258 | 0.009 |
| 0m/2m x L/R | 1 | | 0.086 | 0.7703 | 0.086 | 0.077 | 0.7827 | 0.077 | | 0.032 | 0.8596 | 0.032 | 0.044 | 0.8351 | 0.044 |
| C/E x 0m/2m x L/R | 1 | | 0.031 | 0.8608 | 0.031 | 0.291 | 0.5924 | 0.291 | | 0.043 | 0.8364 | 0.043 | 0.02 | 0.8885 | 0.02 |
| Group | 40 | | | | | | | | | | | | | | |
| D1/D2 | 1 | | 5.253 | 0.0273 | 5.253 | 1.13 | 0.2941 | 1.13 | | 0.836 | 0.366 | 0.836 | 0.021 | 0.8842 | 0.021 |
| D1/D2 x C/E | 1 | | 3.001 | 0.0909 | 3.001 | 2.031 | 0.1619 | 2.031 | | 0.151 | 0.6996 | 0.151 | 1.278 | 0.2651 | 1.278 |
| D1/D2 x 0m/2m | 1 | | 0.13 | 0.7201 | 0.13 | 2.707 | 0.1077 | 2.707 | | 0.835 | 0.3663 | 0.835 | 1.944 | 0.1709 | 1.944 |
| D1/D2 x L/R | 1 | | 1.276 | 0.2654 | 1.276 | 0.872 | 0.356 | 0.872 | | 0.303 | 0.5852 | 0.303 | 0.018 | 0.8932 | 0.018 |
| D1/D2 x C/E x 0m/2m | 1 | | 0.1 | 0.7537 | 0.1 | 0.763 | 0.3875 | 0.763 | | 1.146 | 0.2909 | 1.146 | 0.347 | 0.559 | 0.347 |
| D1/D2 x C/E x L/R | 1 | | 0.256 | 0.6158 | 0.256 | 0.649 | 0.4252 | 0.649 | | 0.343 | 0.5612 | 0.343 | 1.228 | 0.2745 | 1.228 |
| D1/D2 x 0m/2m x L/R | 1 | | 0.688 | 0.4119 | 0.688 | 2.898 | 0.0965 | 2.898 | | 0.819 | 0.3708 | 0.819 | 0.35 | 0.5573 | 0.35 |
| D1/D2 x C/E x 0m/2m x L/R | 1 | | 0.471 | 0.4964 | 0.471 | 0.13 | 0.7203 | 0.13 | | 0.044 | 0.8345 | 0.044 | 0.509 | 0.4796 | 0.509 |
| D1/D2 x Group | 40 | | | | | | | | | | | | | | |
| 15bins | 14 | | 1.242 | 0.2405 | 17.384 | 0.688 | 0.787 | 9.635 | | 1.575 | 0.0815 | 22.056 | 1.24 | 0.2418 | 17.357 |
| 15bins x C/E | 14 | | 0.491 | 0.9384 | 6.873 | 0.496 | 0.9356 | 6.947 | | 0.718 | 0.7569 | 10.054 | 1.091 | 0.3628 | 15.271 |
| 15bins x 0m/2m | 14 | | 2.87 | 0.0003 | 40.182 | 0.688 | 0.7868 | 9.637 | | 2.318 | 0.0042 | 32.458 | 1.594 | 0.0764 | 22.311 |
| 15bins x L/R | 14 | | 0.669 | 0.8051 | 9.371 | 0.912 | 0.5452 | 12.773 | | 0.398 | 0.9756 | 5.566 | 1.123 | 0.3342 | 15.718 |
| 15bins x C/E x 0m/2m | 14 | | 1.038 | 0.4136 | 14.527 | 0.855 | 0.6084 | 11.974 | | 0.636 | 0.8357 | 8.903 | 0.862 | 0.6014 | 12.062 |
| 15bins x C/E x L/R | 14 | | 1.051 | 0.4009 | 14.708 | 0.501 | 0.9329 | 7.016 | | 1.321 | 0.1895 | 18.496 | 1.378 | 0.1587 | 19.29 |
| 15bins x 0m/2m x L/R | 14 | | 1.207 | 0.2656 | 16.897 | 1.662 | 0.0596 | 23.27 | | 0.889 | 0.5711 | 12.444 | 1.044 | 0.4074 | 14.614 |
| 15bins x C/E x 0m/2m x L/R | 14 | | 0.576 | 0.8846 | 8.062 | 0.836 | 0.6303 | 11.698 | | 0.918 | 0.5393 | 12.848 | 1.095 | 0.3589 | 15.33 |
| 15bins x Group | 560 | | | | | | | | | | | | | | |
| D1/D2 x 15bins | 14 | | 1.478 | 0.1141 | 20.692 | 1.849 | 0.0293 | 25.88 | | 1.879 | 0.0261 | 26.3 | 1.071 | 0.3814 | 14.993 |
| D1/D2 x 15bins x C/E | 14 | | 0.447 | 0.9586 | 6.259 | 1.119 | 0.3375 | 15.664 | | 0.556 | 0.8989 | 7.783 | 0.612 | 0.8558 | 8.575 |
| D1/D2 x 15bins x 0m/2m | 14 | | 0.477 | 0.9453 | 6.678 | 0.506 | 0.9302 | 7.084 | | 0.659 | 0.8148 | 9.226 | 0.824 | 0.6437 | 11.529 |
| D1/D2 x 15bins x L/R | 14 | | 1.194 | 0.2752 | 16.72 | 0.49 | 0.939 | 6.855 | | 1.544 | 0.091 | 21.615 | 0.448 | 0.9582 | 6.275 |
| D1/D2 x 15bins x C/E x 0m/2m | 14 | | 1.291 | 0.2077 | 18.075 | 1.104 | 0.3512 | 15.449 | | 0.503 | 0.9317 | 7.046 | 0.77 | 0.7027 | 10.775 |
| D1/D2 x 15bins x C/E x L/R | 14 | | 1.14 | 0.319 | 15.964 | 0.945 | 0.51 | 13.225 | | 1.164 | 0.299 | 16.301 | 0.727 | 0.748 | 10.176 |
| D1/D2 x 15bins x 0m/2m x L/R | 14 | | 1.195 | 0.2745 | 16.734 | 0.991 | 0.4613 | 13.869 | | 0.934 | 0.522 | 13.069 | 1.37 | 0.1626 | 19.183 |
| D1/D2 x 15bins x C/E x 0m/2m x L/R | 14 | | 0.942 | 0.5126 | 13.191 | 1.061 | 0.3907 | 14.856 | | 1.33 | 0.1846 | 18.616 | 1.185 | 0.2824 | 16.591 |
| D1/D2 x 15bins x Group | 560 | | | | | | | | | | | | | | |

**Table S3. ANOVA table of PLV and MI between iHP/vHP and PC in both hemispheres during a 150-s delay in the home cage.** PLV and MI between iHP/vHP (8-10 Hz) and PC (51-100 Hz) were analyzed using the same repeated-measures ANOVA as in Table S1. See Table S1 for details on factors, significance criteria, and abbreviations.

| PLV and MI in home cage | df | PLV: LiHP 8-10Hz_LPC 51-100Hz F-value | p-value | Lambda | PLV: RiHP 8-10Hz_RPC 51-100Hz F-value | p-value | Lambda | MI: LiHP 8-10Hz_LPC 51-100Hz F-value | p-value | Lambda | MI: RiHP 8-10Hz_RPC 51-100Hz F-value | p-value | Lambda |
|---|---|---|---|---|---|---|---|---|---|---|---|---|---|
| C/E | 1 | 12.394 | 0.0011 | 12.394 | 6.589 | 0.0141 | 6.589 | 6.749 | 0.0131 | 6.749 | 4.862 | 0.0333 | 4.862 |
| 0m/2m | 1 | 0.832 | 0.3672 | 0.832 | 0.472 | 0.4961 | 0.472 | 1.405 | 0.2429 | 1.405 | 1.747 | 0.1937 | 1.747 |
| L/R | 1 | 0.118 | 0.7333 | 0.118 | 0.273 | 0.6043 | 0.273 | 0.115 | 0.7367 | 0.115 | 0.624 | 0.4343 | 0.624 |
| C/E x 0m/2m | 1 | 0.332 | 0.5678 | 0.332 | 0.029 | 0.8657 | 0.029 | 1.01 | 0.3209 | 1.01 | 0.458 | 0.5026 | 0.458 |
| C/E x L/R | 1 | 0.159 | 0.692 | 0.159 | 0.161 | 0.6908 | 0.161 | 0.203 | 0.655 | 0.203 | 0.53 | 0.471 | 0.53 |
| 0m/2m x L/R | 1 | 0.231 | 0.6337 | 0.231 | 0.252 | 0.6184 | 0.252 | 0.036 | 0.8506 | 0.036 | 0.436 | 0.5128 | 0.436 |
| C/E x 0m/2m x L/R | 1 | 0.002 | 0.9666 | 0.002 | 0.334 | 0.5667 | 0.334 | 0.186 | 0.6687 | 0.186 | 0.094 | 0.7605 | 0.094 |
| Group | 40 | | | | | | | | | | | | |
| D1/D2 | 1 | 0.015 | 0.9021 | 0.015 | 2.857 | 0.0988 | 2.857 | 0.154 | 0.6965 | 0.154 | 0.209 | 0.6499 | 0.209 |
| D1/D2 x C/E | 1 | 0.228 | 0.6355 | 0.228 | 4.62 | 0.0377 | 4.62 | 0.734 | 0.3966 | 0.734 | 1.579 | 0.2162 | 1.579 |
| D1/D2 x 0m/2m | 1 | 0.092 | 0.7635 | 0.092 | 0.014 | 0.9064 | 0.014 | 1.742 | 0.1944 | 1.742 | 0.724 | 0.3998 | 0.724 |
| D1/D2 x L/R | 1 | 0.214 | 0.6461 | 0.214 | 1.09 | 0.3028 | 1.09 | 0.341 | 0.5625 | 0.341 | 0.066 | 0.7988 | 0.066 |
| D1/D2 x C/E x 0m/2m | 1 | 8.3 | 0.0063 | 8.3 | 1.414 | 0.2414 | 1.414 | 0.004 | 0.9495 | 0.004 | 0.163 | 0.6882 | 0.163 |
| D1/D2 x C/E x L/R | 1 | 2.707 | 0.1078 | 2.707 | 1.301 | 0.2608 | 1.301 | 0.088 | 0.7678 | 0.088 | 0.004 | 0.9525 | 0.004 |
| D1/D2 x 0m/2m x L/R | 1 | 1.838 | 0.1827 | 1.838 | 0.0005 | 0.9826 | 0.0005 | 0.189 | 0.6661 | 0.189 | 0.792 | 0.3789 | 0.792 |
| D1/D2 x C/E x 0m/2m x L/R | 1 | 0.054 | 0.8172 | 0.054 | 1.371 | 0.2485 | 1.371 | 1.479 | 0.2311 | 1.479 | 3.599 | 0.0651 | 3.599 |
| D1/D2 x Group | 40 | | | | | | | | | | | | |
| 15bins | 14 | 0.925 | 0.5312 | 12.952 | 1.359 | 0.1684 | 19.028 | 2.021 | 0.0147 | 28.298 | 2.766 | 0.0005 | 38.727 |
| 15bins x C/E | 14 | 0.787 | 0.6838 | 11.019 | 0.756 | 0.7172 | 10.586 | 1.056 | 0.3955 | 14.786 | 1.285 | 0.2113 | 17.996 |
| 15bins x 0m/2m | 14 | 0.641 | 0.8308 | 8.98 | 1.244 | 0.2387 | 17.419 | 1.06 | 0.3914 | 14.846 | 0.853 | 0.6112 | 11.939 |
| 15bins x L/R | 14 | 1.013 | 0.4386 | 14.177 | 0.87 | 0.5923 | 12.177 | 0.774 | 0.6978 | 10.839 | 0.947 | 0.5078 | 13.253 |
| 15bins x C/E x 0m/2m | 14 | 0.798 | 0.6713 | 11.179 | 0.991 | 0.461 | 13.872 | 1.363 | 0.1662 | 19.086 | 0.424 | 0.9672 | 5.939 |
| 15bins x C/E x L/R | 14 | 0.523 | 0.9203 | 7.322 | 0.758 | 0.7149 | 10.617 | 0.685 | 0.79 | 9.592 | 0.481 | 0.9435 | 6.731 |
| 15bins x 0m/2m x L/R | 14 | 1.063 | 0.3893 | 14.876 | 2.067 | 0.0121 | 28.94 | 1.222 | 0.2544 | 17.111 | 1.172 | 0.2927 | 16.41 |
| 15bins x C/E x 0m/2m x L/R | 14 | 0.703 | 0.7721 | 9.845 | 0.92 | 0.5372 | 12.874 | 0.801 | 0.6685 | 11.215 | 0.755 | 0.7182 | 10.572 |
| 15bins x Group | 560 | | | | | | | | | | | | |
| D1/D2 x 15bins | 14 | 1.689 | 0.0539 | 23.651 | 2.122 | 0.0096 | 29.712 | 1.856 | 0.0285 | 25.98 | 3.195 | <.0001 | 44.728 |
| D1/D2 x 15bins x C/E | 14 | 1.134 | 0.3241 | 15.88 | 0.855 | 0.6089 | 11.967 | 0.746 | 0.7276 | 10.449 | 0.848 | 0.6161 | 11.877 |
| D1/D2 x 15bins x 0m/2m | 14 | 1.316 | 0.1928 | 18.418 | 1.331 | 0.1841 | 18.629 | 0.939 | 0.516 | 13.147 | 1.216 | 0.2588 | 17.027 |
| D1/D2 x 15bins x L/R | 14 | 0.599 | 0.8666 | 8.388 | 0.938 | 0.517 | 13.134 | 0.908 | 0.5499 | 12.713 | 0.93 | 0.5262 | 13.016 |
| D1/D2 x 15bins x C/E x 0m/2m | 14 | 1.79 | 0.0369 | 25.055 | 0.903 | 0.5556 | 12.641 | 0.554 | 0.9003 | 7.754 | 0.724 | 0.751 | 10.136 |
| D1/D2 x 15bins x C/E x L/R | 14 | 0.543 | 0.9079 | 7.596 | 0.893 | 0.5661 | 12.507 | 0.577 | 0.8838 | 8.076 | 0.547 | 0.9046 | 7.665 |
| D1/D2 x 15bins x 0m/2m x L/R | 14 | 1.911 | 0.023 | 26.75 | 1.112 | 0.344 | 15.561 | 0.485 | 0.9411 | 6.797 | 0.619 | 0.8507 | 8.66 |
| D1/D2 x 15bins x C/E x 0m/2m x L/R | 14 | 1.254 | 0.2323 | 17.549 | 0.637 | 0.8347 | 8.918 | 0.215 | 0.999 | 3.008 | 0.67 | 0.8045 | 9.38 |
| D1/D2 x 15bins x Group | 560 | | | | | | | | | | | | |

| PLV and MI in home cage | df | PLV: LvHP 8-10Hz_LPC 51-100Hz F-value | p-value | Lambda | PLV: RvHP 8-10Hz_RPC 51-100Hz F-value | p-value | Lambda | MI: LvHP 8-10Hz_LPC 51-100Hz F-value | p-value | Lambda | MI: RvHP 8-10Hz_RPC 51-100Hz F-value | p-value | Lambda |
|---|---|---|---|---|---|---|---|---|---|---|---|---|---|
| C/E | 1 | 25.271 | <.0001 | 25.271 | 25.218 | <.0001 | 25.218 | 9.332 | 0.004 | 9.332 | 13.676 | 0.0007 | 13.676 |
| 0m/2m | 1 | 0.031 | 0.8611 | 0.031 | 0.981 | 0.3279 | 0.981 | 0.718 | 0.4019 | 0.718 | 1.001 | 0.3231 | 1.001 |
| L/R | 1 | 0.214 | 0.6463 | 0.214 | 0.028 | 0.8679 | 0.028 | 0.009 | 0.925 | 0.009 | 0.751 | 0.3915 | 0.751 |
| C/E x 0m/2m | 1 | 0.285 | 0.5961 | 0.285 | 0.201 | 0.6566 | 0.201 | 1.058 | 0.3099 | 1.058 | 0.778 | 0.383 | 0.778 |
| C/E x L/R | 1 | 0.036 | 0.851 | 0.036 | 0.235 | 0.6306 | 0.235 | 0.115 | 0.7361 | 0.115 | 1.513 | 0.2258 | 1.513 |
| 0m/2m x L/R | 1 | 0.009 | 0.9252 | 0.009 | 0.813 | 0.3727 | 0.813 | 0.29 | 0.5932 | 0.29 | 0.788 | 0.38 | 0.788 |
| C/E x 0m/2m x L/R | 1 | 0.002 | 0.9641 | 0.002 | 0.005 | 0.9431 | 0.005 | 0.465 | 0.4994 | 0.465 | 0.122 | 0.7291 | 0.122 |
| Group | 40 | | | | | | | | | | | | |
| D1/D2 | 1 | 0.221 | 0.6407 | 0.221 | 0.006 | 0.9394 | 0.006 | 0.142 | 0.7083 | 0.142 | 0.015 | 0.902 | 0.015 |
| D1/D2 x C/E | 1 | 0.946 | 0.3366 | 0.946 | 0.201 | 0.656 | 0.201 | 0.203 | 0.6545 | 0.203 | 0.816 | 0.3718 | 0.816 |
| D1/D2 x 0m/2m | 1 | 5.941 | 0.0193 | 5.941 | 0.56 | 0.4585 | 0.56 | 1.895 | 0.1763 | 1.895 | 1.955 | 0.1697 | 1.955 |
| D1/D2 x L/R | 1 | 0.181 | 0.6726 | 0.181 | 0.589 | 0.4471 | 0.589 | 0.003 | 0.954 | 0.003 | 0.087 | 0.7693 | 0.087 |
| D1/D2 x C/E x 0m/2m | 1 | 0.415 | 0.5231 | 0.415 | 0.013 | 0.9093 | 0.013 | 0.345 | 0.5602 | 0.345 | 0.001 | 0.9798 | 0.001 |
| D1/D2 x C/E x L/R | 1 | 2.211 | 0.1449 | 2.211 | 0.053 | 0.8195 | 0.053 | 0.685 | 0.4128 | 0.685 | 0.331 | 0.5684 | 0.331 |
| D1/D2 x 0m/2m x L/R | 1 | 3.182 | 0.082 | 3.182 | 2E-06 | 0.9989 | 2E-06 | 1.012 | 0.3204 | 1.012 | 0.528 | 0.4719 | 0.528 |
| D1/D2 x C/E x 0m/2m x L/R | 1 | 1.135 | 0.2932 | 1.135 | 0.471 | 0.4966 | 0.471 | 0.378 | 0.5422 | 0.378 | 0.774 | 0.3843 | 0.774 |
| D1/D2 x Group | 40 | | | | | | | | | | | | |
| 15bins | 14 | 0.924 | 0.5322 | 12.939 | 0.98 | 0.4728 | 13.715 | 2.317 | 0.0042 | 32.439 | 1.37 | 0.1626 | 19.181 |
| 15bins x C/E | 14 | 1.247 | 0.237 | 17.454 | 1.303 | 0.2002 | 18.245 | 1.958 | 0.019 | 27.415 | 1.691 | 0.0536 | 23.674 |
| 15bins x 0m/2m | 14 | 1.041 | 0.4099 | 14.579 | 1.347 | 0.1752 | 18.851 | 0.701 | 0.7743 | 9.814 | 0.511 | 0.9275 | 7.15 |
| 15bins x L/R | 14 | 0.455 | 0.9555 | 6.364 | 1.026 | 0.4249 | 14.367 | 0.631 | 0.8401 | 8.832 | 0.99 | 0.4615 | 13.865 |
| 15bins x C/E x 0m/2m | 14 | 1.027 | 0.4236 | 14.385 | 1.341 | 0.1784 | 18.77 | 0.707 | 0.7678 | 9.904 | 0.474 | 0.9469 | 6.631 |
| 15bins x C/E x L/R | 14 | 0.342 | 0.9882 | 4.788 | 0.642 | 0.8303 | 8.988 | 0.92 | 0.5366 | 12.882 | 0.901 | 0.5579 | 12.612 |
| 15bins x 0m/2m x L/R | 14 | 0.563 | 0.8941 | 7.878 | 1.631 | 0.0668 | 22.834 | 1.729 | 0.0464 | 24.212 | 0.94 | 0.5147 | 13.164 |
| 15bins x C/E x 0m/2m x L/R | 14 | 0.396 | 0.976 | 5.55 | 1.039 | 0.4125 | 14.542 | 1.342 | 0.1776 | 18.79 | 1.166 | 0.2976 | 16.324 |
| 15bins x Group | 560 | | | | | | | | | | | | |
| D1/D2 x 15bins | 14 | 1.43 | 0.1341 | 20.017 | 0.871 | 0.591 | 12.193 | 1.18 | 0.2862 | 16.523 | 3.218 | <.0001 | 45.056 |
| D1/D2 x 15bins x C/E | 14 | 1.063 | 0.3891 | 14.878 | 0.908 | 0.5495 | 12.718 | 0.8 | 0.6694 | 11.204 | 2.072 | 0.0119 | 29.007 |
| D1/D2 x 15bins x 0m/2m | 14 | 2.001 | 0.016 | 28.012 | 0.397 | 0.9759 | 5.551 | 0.473 | 0.9472 | 6.622 | 0.713 | 0.7627 | 9.976 |
| D1/D2 x 15bins x L/R | 14 | 1.5 | 0.1058 | 21.004 | 0.709 | 0.7665 | 9.922 | 1.089 | 0.3645 | 15.245 | 0.94 | 0.5147 | 13.164 |
| D1/D2 x 15bins x C/E x 0m/2m | 14 | 0.909 | 0.5491 | 12.722 | 1.422 | 0.1377 | 19.904 | 0.39 | 0.9777 | 5.463 | 0.886 | 0.574 | 12.408 |
| D1/D2 x 15bins x C/E x L/R | 14 | 1.005 | 0.4467 | 14.066 | 1.336 | 0.1808 | 18.709 | 1.201 | 0.2698 | 16.819 | 0.799 | 0.6707 | 11.187 |
| D1/D2 x 15bins x 0m/2m x L/R | 14 | 1.638 | 0.0652 | 22.926 | 0.947 | 0.5069 | 13.265 | 0.694 | 0.7808 | 9.723 | 0.701 | 0.7746 | 9.81 |
| D1/D2 x 15bins x C/E x 0m/2m x L/R | 14 | 0.875 | 0.5865 | 12.249 | 0.647 | 0.8258 | 9.058 | 0.778 | 0.6938 | 10.891 | 1.24 | 0.2416 | 17.36 |
| D1/D2 x 15bins x Group | 560 | | | | | | | | | | | | |

**Table S4. ANOVA table of spectral power in both hemispheres during the 10-s period surrounding arm entry.** Power in the 8-10 Hz and 51-100 Hz bands across brain regions was analyzed using a repeated-measures ANOVA. The model included five within-subject factors in the task matrix (accuracy [C/E] × distance [0m/2m] × delay length [75 s/150 s] × arm entry [L/R] × time [consecutive 1-s bins; 5bins]) and one between-subject factor (period [A1 vs. A2] within the arm entry epoch). Cells with p < 0.05 are highlighted in color in the final row. Colors indicate individual indices, and multiple colors denote interaction effects. The color-coding scheme follows that described in Table S1, including rules for assignment of main effects and interaction effects. See Table S1 for details on abbreviations.

| Power around T-maze arm entry | df | | 8-10 Hz | | | 51-100 Hz | | | | 8-10 Hz | | | 51-100 Hz | | | | 8-10 Hz | | | 51-100 Hz | | |
|---|---|---|---|---|---|---|---|---|---|---|---|---|---|---|---|---|---|---|---|---|---|---|
| | | | F-value | p-value | Lam-bda | F-value | p-value | Lam-bda | | F-value | p-value | Lam-bda | F-value | p-value | Lam-bda | | F-value | p-value | Lam-bda | F-value | p-value | Lam-bda |
| C/E | 1 | | 0.055 | 0.815 | 0.055 | 0.014 | 0.906 | 0.014 | | 0.622 | 0.433 | 0.622 | 0.075 | 0.785 | 0.075 | | 0.508 | 0.478 | 0.508 | 0.026 | 0.873 | 0.026 |
| 0m/2m | 1 | | 0.013 | 0.909 | 0.013 | 0.038 | 0.846 | 0.038 | | 0.268 | 0.606 | 0.268 | 0.019 | 0.889 | 0.019 | | 0.004 | 0.951 | 0.004 | 0.010 | 0.923 | 0.010 |
| 75s/150s | 1 | | 0.094 | 0.760 | 0.094 | 0.071 | 0.790 | 0.071 | | 0.170 | 0.681 | 0.170 | 0.096 | 0.758 | 0.096 | | 0.011 | 0.915 | 0.011 | 0.002 | 0.964 | 0.002 |
| L/R | 1 | | 0.073 | 0.788 | 0.073 | 0.009 | 0.926 | 0.009 | | 0.027 | 0.869 | 0.027 | 0.000 | 0.997 | 0.000 | | 0.001 | 0.982 | 0.001 | 0.000 | 0.993 | 0.000 |
| C/E x 0m/2m | 1 | | 0.002 | 0.966 | 0.002 | 0.000 | 0.995 | 0.000 | | 0.017 | 0.898 | 0.017 | 0.002 | 0.963 | 0.002 | | 0.007 | 0.934 | 0.007 | 0.002 | 0.968 | 0.002 |
| C/E x 75s/150s | 1 | | 0.033 | 0.857 | 0.033 | 0.005 | 0.942 | 0.005 | | 0.026 | 0.873 | 0.026 | 0.022 | 0.883 | 0.022 | | 0.005 | 0.944 | 0.005 | 0.000 | 0.999 | 0.000 |
| C/E x L/R | 1 | | 0.687 | 0.410 | 0.687 | 0.000 | 0.986 | 0.000 | | 0.001 | 0.969 | 0.001 | 0.000 | 0.983 | 0.000 | | 0.020 | 0.888 | 0.020 | 0.001 | 0.970 | 0.001 |
| 0m/2m x 75s/150s | 1 | | 0.330 | 0.567 | 0.330 | 0.000 | 0.986 | 0.000 | | 0.002 | 0.964 | 0.002 | 0.009 | 0.924 | 0.009 | | 0.004 | 0.949 | 0.004 | 0.000 | 0.997 | 0.000 |
| 0m/2m x L/R | 1 | | 0.002 | 0.965 | 0.002 | 0.006 | 0.941 | 0.006 | | 0.001 | 0.974 | 0.001 | 0.016 | 0.901 | 0.016 | | 0.002 | 0.969 | 0.002 | 0.001 | 0.977 | 0.001 |
| 75s/150s x L/R | 1 | | 0.472 | 0.494 | 0.472 | 0.029 | 0.865 | 0.029 | | 0.014 | 0.905 | 0.014 | 0.001 | 0.970 | 0.001 | | 0.012 | 0.912 | 0.012 | 0.000 | 0.989 | 0.000 |
| C/E x 0m/2m x 75s/150s | 1 | | 0.017 | 0.897 | 0.017 | 0.000 | 0.996 | 0.000 | | 0.005 | 0.945 | 0.005 | 0.016 | 0.900 | 0.016 | | 0.004 | 0.952 | 0.004 | 0.000 | 0.984 | 0.000 |
| C/E x 0m/2m x L/R | 1 | | 0.034 | 0.855 | 0.034 | 0.004 | 0.952 | 0.004 | | 0.021 | 0.884 | 0.021 | 0.036 | 0.850 | 0.036 | | 0.003 | 0.959 | 0.003 | 0.000 | 0.992 | 0.000 |
| C/E x 75s/150s x L/R | 1 | | 0.879 | 0.351 | 0.879 | 0.011 | 0.917 | 0.011 | | 0.005 | 0.945 | 0.005 | 0.002 | 0.962 | 0.002 | | 0.001 | 0.978 | 0.001 | 0.000 | 0.985 | 0.000 |
| 0m/2m x 75s/150s x L/R | 1 | | 0.048 | 0.828 | 0.048 | 0.005 | 0.942 | 0.005 | | 0.053 | 0.819 | 0.053 | 0.002 | 0.963 | 0.002 | | 0.013 | 0.909 | 0.013 | 0.001 | 0.972 | 0.001 |
| C/E x 0m/2m x 75s/150s x L/R | 1 | | 0.306 | 0.582 | 0.306 | 0.001 | 0.974 | 0.001 | | 0.013 | 0.910 | 0.013 | 0.001 | 0.975 | 0.001 | | 0.000 | 0.988 | 0.000 | 0.000 | 0.999 | 0.000 |
| Group | 80 | | | | | | | | | | | | | | | | | | | | | |
| A1/A2 | 1 | | 0.101 | 0.751 | 0.101 | 1.903 | 0.172 | 1.903 | | 22.651 | <.0001 | 22.651 | 0.024 | 0.877 | 0.024 | | 0.761 | 0.386 | 0.761 | 14.476 | 0.0003 | 14.476 |
| A1/A2 x C/E | 1 | | 0.002 | 0.963 | 0.002 | 1.778 | 0.186 | 1.778 | | 10.307 | 0.002 | 10.307 | 0.092 | 0.762 | 0.092 | | 6.277 | 0.014 | 6.277 | 1.987 | 0.163 | 1.987 |
| A1/A2 x 0m/2m | 1 | | 5.924 | 0.017 | 5.924 | 1.088 | 0.300 | 1.088 | | 9.680 | 0.003 | 9.680 | 4.560 | 0.036 | 4.560 | | 11.208 | 0.001 | 11.208 | 2.154 | 0.146 | 2.154 |
| A1/A2 x 75s/150s | 1 | | 1.279 | 0.262 | 1.279 | 0.155 | 0.695 | 0.155 | | 1.578 | 0.213 | 1.578 | 0.476 | 0.492 | 0.476 | | 0.809 | 0.371 | 0.809 | 0.023 | 0.879 | 0.023 |
| A1/A2 x L/R | 1 | | 0.223 | 0.638 | 0.223 | 0.001 | 0.975 | 0.001 | | 0.002 | 0.967 | 0.002 | 0.384 | 0.537 | 0.384 | | 2.287 | 0.134 | 2.287 | 1.835 | 0.179 | 1.835 |
| A1/A2 x C/E x 0m/2m | 1 | | 4.516 | 0.037 | 4.516 | 0.056 | 0.814 | 0.056 | | 0.183 | 0.670 | 0.183 | 0.613 | 0.436 | 0.613 | | 0.187 | 0.667 | 0.187 | 0.119 | 0.731 | 0.119 |
| A1/A2 x C/E x 75s/150s | 1 | | 0.426 | 0.516 | 0.426 | 0.796 | 0.375 | 0.796 | | 0.020 | 0.888 | 0.020 | 0.208 | 0.650 | 0.208 | | 0.013 | 0.909 | 0.013 | 0.096 | 0.757 | 0.096 |
| A1/A2 x C/E x L/R | 1 | | 0.616 | 0.435 | 0.616 | 0.102 | 0.750 | 0.102 | | 0.022 | 0.883 | 0.022 | 0.271 | 0.604 | 0.271 | | 0.254 | 0.615 | 0.254 | 0.143 | 0.707 | 0.143 |
| A1/A2 x 0m/2m x 75s/150s | 1 | | 0.093 | 0.761 | 0.093 | 0.437 | 0.510 | 0.437 | | 0.033 | 0.857 | 0.033 | 0.430 | 0.514 | 0.430 | | 0.244 | 0.623 | 0.244 | 0.059 | 0.808 | 0.059 |
| A1/A2 x 0m/2m x L/R | 1 | | 0.572 | 0.452 | 0.572 | 0.396 | 0.531 | 0.396 | | 0.404 | 0.527 | 0.404 | 0.732 | 0.395 | 0.732 | | 0.031 | 0.860 | 0.031 | 0.346 | 0.558 | 0.346 |
| A1/A2 x 75s/150s x L/R | 1 | | 0.002 | 0.961 | 0.002 | 0.652 | 0.422 | 0.652 | | 0.119 | 0.731 | 0.119 | 0.379 | 0.540 | 0.379 | | 0.078 | 0.781 | 0.078 | 0.088 | 0.768 | 0.088 |
| A1/A2 x C/E x 0m/2m x 75s/150s | 1 | | 0.888 | 0.349 | 0.888 | 0.269 | 0.605 | 0.269 | | 0.048 | 0.828 | 0.048 | 0.315 | 0.576 | 0.315 | | 0.029 | 0.866 | 0.029 | 0.013 | 0.910 | 0.013 |
| A1/A2 x C/E x 0m/2m x L/R | 1 | | 0.515 | 0.475 | 0.515 | 0.163 | 0.687 | 0.163 | | 0.022 | 0.883 | 0.022 | 0.161 | 0.689 | 0.161 | | 0.114 | 0.737 | 0.114 | 0.012 | 0.915 | 0.012 |
| A1/A2 x C/E x 75s/150s x L/R | 1 | | 0.597 | 0.442 | 0.597 | 0.704 | 0.404 | 0.704 | | 0.065 | 0.799 | 0.065 | 0.548 | 0.461 | 0.548 | | 0.020 | 0.887 | 0.020 | 0.013 | 0.909 | 0.013 |
| A1/A2 x 0m/2m x 75s/150s x L/R | 1 | | 1.879 | 0.174 | 1.879 | 0.551 | 0.460 | 0.551 | | 0.026 | 0.872 | 0.026 | 0.416 | 0.521 | 0.416 | | 0.728 | 0.396 | 0.728 | 1.018 | 0.316 | 1.018 |
| A1/A2 x C/E x 0m/2m x 75s/150s x L/R | 1 | | 0.000 | 0.992 | 0.000 | 0.000 | 0.983 | 0.000 | | 0.138 | 0.711 | 0.138 | 0.131 | 0.718 | 0.131 | | 0.175 | 0.677 | 0.175 | 0.060 | 0.807 | 0.060 |
| A1/A2 x Group | 80 | | | | | | | | | | | | | | | | | | | | | |
| 5bins | 4 | LPC | 6.003 | 0.0001 | 24.010 | 3.613 | 0.007 | 14.450 | LiHP | 8.099 | <.0001 | 32.397 | 1.122 | 0.346 | 4.490 | LvHP | 7.880 | <.0001 | 31.518 | 7.438 | <.0001 | 29.750 |
| 5bins x C/E | 4 | | 0.430 | 0.787 | 1.722 | 3.681 | 0.006 | 14.724 | | 4.788 | 0.001 | 19.150 | 1.457 | 0.215 | 5.828 | | 5.866 | 0.0001 | 23.463 | 3.259 | 0.012 | 13.036 |
| 5bins x 0m/2m | 4 | | 0.669 | 0.614 | 2.678 | 4.653 | 0.001 | 18.613 | | 0.721 | 0.578 | 2.882 | 2.983 | 0.019 | 11.930 | | 2.208 | 0.068 | 8.833 | 0.832 | 0.506 | 3.326 |
| 5bins x 75s/150s | 4 | | 0.881 | 0.476 | 3.523 | 1.286 | 0.275 | 5.143 | | 0.826 | 0.509 | 3.304 | 0.439 | 0.780 | 1.756 | | 1.639 | 0.164 | 6.557 | 1.340 | 0.255 | 5.360 |
| 5bins x L/R | 4 | | 0.564 | 0.689 | 2.256 | 0.237 | 0.918 | 0.947 | | 1.121 | 0.347 | 4.483 | 0.728 | 0.573 | 2.912 | | 0.221 | 0.927 | 0.884 | 0.257 | 0.905 | 1.028 |
| 5bins x C/E x 0m/2m | 4 | | 1.531 | 0.193 | 6.124 | 0.231 | 0.921 | 0.923 | | 0.192 | 0.942 | 0.770 | 0.990 | 0.413 | 3.958 | | 1.063 | 0.375 | 4.251 | 0.550 | 0.699 | 2.201 |
| 5bins x C/E x 75s/150s | 4 | | 1.735 | 0.142 | 6.938 | 0.867 | 0.484 | 3.466 | | 0.497 | 0.738 | 1.987 | 0.891 | 0.470 | 3.563 | | 0.382 | 0.821 | 1.529 | 1.016 | 0.399 | 4.063 |
| 5bins x C/E x L/R | 4 | | 2.390 | 0.051 | 9.561 | 1.541 | 0.190 | 6.163 | | 0.787 | 0.534 | 3.147 | 0.972 | 0.423 | 3.890 | | 0.426 | 0.790 | 1.702 | 0.176 | 0.951 | 0.704 |
| 5bins x 0m/2m x 75s/150s | 4 | | 0.451 | 0.771 | 1.805 | 1.644 | 0.163 | 6.577 | | 1.205 | 0.309 | 4.819 | 1.455 | 0.216 | 5.820 | | 0.923 | 0.451 | 3.692 | 0.860 | 0.488 | 3.441 |
| 5bins x 0m/2m x L/R | 4 | | 2.469 | 0.045 | 9.877 | 1.157 | 0.330 | 4.627 | | 0.459 | 0.766 | 1.836 | 0.524 | 0.718 | 2.095 | | 1.448 | 0.218 | 5.792 | 1.404 | 0.232 | 5.618 |
| 5bins x 75s/150s x L/R | 4 | | 1.150 | 0.333 | 4.601 | 1.537 | 0.191 | 6.148 | | 0.502 | 0.734 | 2.009 | 1.137 | 0.339 | 4.548 | | 1.005 | 0.405 | 4.020 | 0.738 | 0.566 | 2.953 |
| 5bins x C/E x 0m/2m x 75s/150s | 4 | | 0.339 | 0.851 | 1.357 | 1.660 | 0.159 | 6.639 | | 0.372 | 0.829 | 1.488 | 1.918 | 0.107 | 7.672 | | 0.531 | 0.713 | 2.123 | 0.243 | 0.914 | 0.971 |
| 5bins x C/E x 0m/2m x L/R | 4 | | 0.801 | 0.525 | 3.204 | 0.451 | 0.771 | 1.806 | | 0.154 | 0.961 | 0.616 | 0.664 | 0.617 | 2.658 | | 0.134 | 0.970 | 0.538 | 0.046 | 0.996 | 0.186 |
| 5bins x C/E x 75s/150s x L/R | 4 | | 0.236 | 0.918 | 0.944 | 0.566 | 0.688 | 2.264 | | 0.538 | 0.708 | 2.154 | 0.904 | 0.462 | 3.614 | | 0.859 | 0.489 | 3.434 | 0.488 | 0.745 | 1.951 |
| 5bins x 0m/2m x 75s/150s x L/R | 4 | | 0.841 | 0.500 | 3.366 | 0.563 | 0.690 | 2.253 | | 0.093 | 0.985 | 0.372 | 1.017 | 0.398 | 4.069 | | 0.558 | 0.693 | 2.234 | 0.847 | 0.496 | 3.387 |
| 5bins x C/E x 0m/2m x 75s/150s x L/R | 4 | | 1.562 | 0.184 | 6.249 | 0.137 | 0.968 | 0.550 | | 0.256 | 0.906 | 1.022 | 0.697 | 0.594 | 2.789 | | 0.457 | 0.767 | 1.828 | 0.192 | 0.943 | 0.767 |
| 5bins x Group | 320 | | | | | | | | | | | | | | | | | | | | | |
| A1/A2 x 5bins | 4 | | 3.800 | 0.005 | 15.198 | 1.168 | 0.325 | 4.673 | | 42.929 | <.0001 | ###### | 9.205 | <.0001 | 36.819 | | 42.901 | <.0001 | 171.603 | 9.894 | <.0001 | 39.578 |
| A1/A2 x 5bins x C/E | 4 | | 1.049 | 0.382 | 4.198 | 2.845 | 0.024 | 11.380 | | 6.093 | <.0001 | 24.370 | 2.282 | 0.060 | 9.128 | | 4.890 | 0.001 | 19.560 | 2.473 | 0.044 | 9.892 |
| A1/A2 x 5bins x 0m/2m | 4 | | 4.134 | 0.003 | 16.536 | 1.630 | 0.167 | 6.518 | | 0.157 | 0.960 | 0.629 | 3.461 | 0.009 | 13.845 | | 1.101 | 0.356 | 4.403 | 0.742 | 0.564 | 2.966 |
| A1/A2 x 5bins x 75s/150s | 4 | | 0.638 | 0.636 | 2.553 | 0.354 | 0.841 | 1.415 | | 0.189 | 0.944 | 0.754 | 2.049 | 0.087 | 8.197 | | 1.144 | 0.336 | 4.577 | 1.340 | 0.255 | 5.360 |
| A1/A2 x 5bins x L/R | 4 | | 0.611 | 0.655 | 2.445 | 1.238 | 0.295 | 4.950 | | 0.606 | 0.659 | 2.424 | 0.715 | 0.582 | 2.859 | | 0.395 | 0.812 | 1.580 | 0.566 | 0.688 | 2.262 |
| A1/A2 x 5bins x C/E x 0m/2m | 4 | | 0.180 | 0.948 | 0.722 | 1.909 | 0.109 | 7.637 | | 0.304 | 0.875 | 1.217 | 0.927 | 0.449 | 3.706 | | 0.254 | 0.907 | 1.014 | 0.323 | 0.863 | 1.292 |
| A1/A2 x 5bins x C/E x 75s/150s | 4 | | 0.203 | 0.937 | 0.812 | 0.268 | 0.898 | 1.072 | | 0.231 | 0.921 | 0.922 | 1.275 | 0.279 | 5.102 | | 0.293 | 0.882 | 1.173 | 0.523 | 0.719 | 2.093 |
| A1/A2 x 5bins x C/E x L/R | 4 | | 0.341 | 0.850 | 1.363 | 0.622 | 0.647 | 2.487 | | 0.317 | 0.866 | 1.269 | 0.688 | 0.601 | 2.753 | | 0.147 | 0.964 | 0.589 | 0.142 | 0.967 | 0.566 |
| A1/A2 x 5bins x 0m/2m x 75s/150s | 4 | | 0.098 | 0.983 | 0.393 | 1.599 | 0.174 | 6.397 | | 0.583 | 0.675 | 2.331 | 1.157 | 0.330 | 4.627 | | 0.110 | 0.979 | 0.442 | 0.175 | 0.951 | 0.699 |
| A1/A2 x 5bins x 0m/2m x L/R | 4 | | 0.455 | 0.768 | 1.822 | 0.267 | 0.899 | 1.069 | | 0.177 | 0.950 | 0.706 | 0.763 | 0.550 | 3.053 | | 0.731 | 0.572 | 2.922 | 0.715 | 0.582 | 2.858 |
| A1/A2 x 5bins x 75s/150s x L/R | 4 | | 0.558 | 0.694 | 2.230 | 0.249 | 0.910 | 0.996 | | 0.214 | 0.930 | 0.857 | 0.418 | 0.796 | 1.672 | | 0.464 | 0.762 | 1.855 | 0.500 | 0.736 | 2.001 |
| A1/A2 x 5bins x C/E x 0m/2m x 75s/150s | 4 | | 1.094 | 0.360 | 4.375 | 1.167 | 0.325 | 4.667 | | 0.095 | 0.984 | 0.379 | 0.608 | 0.658 | 2.430 | | 0.539 | 0.708 | 2.154 | 0.126 | 0.973 | 0.506 |
| A1/A2 x 5bins x C/E x 0m/2m x L/R | 4 | | 1.910 | 0.109 | 7.639 | 0.644 | 0.632 | 2.574 | | 0.099 | 0.983 | 0.396 | 0.656 | 0.623 | 2.623 | | 0.289 | 0.885 | 1.157 | 0.878 | 0.477 | 3.511 |
| A1/A2 x 5bins x C/E x 75s/150s x L/R | 4 | | 1.969 | 0.099 | 7.876 | 0.260 | 0.903 | 1.042 | | 0.127 | 0.973 | 0.506 | 0.885 | 0.473 | 3.542 | | 0.081 | 0.988 | 0.326 | 0.188 | 0.945 | 0.752 |
| A1/A2 x 5bins x 0m/2m x 75s/150s x L/R | 4 | | 0.563 | 0.690 | 2.251 | 1.118 | 0.348 | 4.473 | | 0.173 | 0.952 | 0.693 | 0.455 | 0.769 | 1.820 | | 0.458 | 0.767 | 1.831 | 0.652 | 0.626 | 2.610 |
| A1/A2 x 5bins x C/E x 0m/2m x 75s/150s x L/R | 4 | | 0.409 | 0.802 | 1.635 | 1.932 | 0.105 | 7.728 | | 0.568 | 0.686 | 2.271 | 1.143 | 0.336 | 4.570 | | 0.266 | 0.900 | 1.063 | 0.171 | 0.953 | 0.685 |
| A1/A2 x 5bins x Group | 320 | | | | | | | | | | | | | | | | | | | | | |
| C/E | 1 | | 0.435 | 0.511 | 0.435 | 0.045 | 0.832 | 0.045 | | 0.383 | 0.538 | 0.383 | 0.108 | 0.743 | 0.108 | | 0.575 | 0.4505 | 0.575 | 0.092 | 0.762 | 0.092 |
| 0m/2m | 1 | | 2.008 | 0.160 | 2.008 | 0.758 | 0.387 | 0.758 | | 0.008 | 0.930 | 0.008 | 0.005 | 0.942 | 0.005 | | 0.198 | 0.6579 | 0.198 | 0.067 | 0.797 | 0.067 |
| 75s/150s | 1 | | 1.064 | 0.306 | 1.064 | 0.803 | 0.373 | 0.803 | | 0.002 | 0.967 | 0.002 | 0.009 | 0.925 | 0.009 | | 0.001 | 0.9793 | 0.001 | 0.001 | 0.972 | 0.001 |
| L/R | 1 | | 0.468 | 0.496 | 0.468 | 0.064 | 0.801 | 0.064 | | 0.212 | 0.646 | 0.212 | 0.041 | 0.840 | 0.041 | | 0.035 | 0.8515 | 0.035 | 0.001 | 0.978 | 0.001 |
| C/E x 0m/2m | 1 | | 0.986 | 0.324 | 0.986 | 0.193 | 0.661 | 0.193 | | 0.007 | 0.936 | 0.007 | 0.002 | 0.962 | 0.002 | | 0.054 | 0.8169 | 0.054 | 0.023 | 0.881 | 0.023 |
| C/E x 75s/150s | 1 | | 0.141 | 0.708 | 0.141 | 0.035 | 0.853 | 0.035 | | 0.173 | 0.679 | 0.173 | 0.037 | 0.848 | 0.037 | | 1E-05 | 0.9973 | 1.1E-05 | 0.004 | 0.948 | 0.004 |
| C/E x L/R | 1 | | 1.363 | 0.247 | 1.363 | 0.830 | 0.365 | 0.830 | | 0.166 | 0.685 | 0.166 | 0.031 | 0.860 | 0.031 | | 0.003 | 0.959 | 0.003 | 0.012 | 0.914 | 0.012 |
| 0m/2m x 75s/150s | 1 | | 0.634 | 0.428 | 0.634 | 0.015 | 0.903 | 0.015 | | 0.082 | 0.775 | 0.082 | 0.056 | 0.814 | 0.056 | | 0.000 | 0.983 | 0.000 | 0.000 | 0.997 | 0.000 |
| 0m/2m x L/R | 1 | | 0.924 | 0.339 | 0.924 | 0.296 | 0.588 | 0.296 | | 0.003 | 0.959 | 0.003 | 0.000 | 0.992 | 0.000 | | 0.008 | 0.929 | 0.008 | 0.003 | 0.956 | 0.003 |
| 75s/150s x L/R | 1 | | 0.093 | 0.761 | 0.093 | 0.009 | 0.923 | 0.009 | | 0.085 | 0.772 | 0.085 | 0.032 | 0.859 | 0.032 | | 0.003 | 0.953 | 0.003 | 0.000 | 0.983 | 0.000 |
| C/E x 0m/2m x 75s/150s | 1 | | 0.694 | 0.407 | 0.694 | 0.078 | 0.781 | 0.078 | | 0.186 | 0.668 | 0.186 | 0.092 | 0.762 | 0.092 | | 0.005 | 0.946 | 0.005 | 0.000 | 0.986 | 0.000 |
| C/E x 0m/2m x L/R | 1 | | 0.719 | 0.399 | 0.719 | 0.136 | 0.714 | 0.136 | | 0.010 | 0.919 | 0.010 | 0.002 | 0.968 | 0.002 | | 0.001 | 0.979 | 0.001 | 0.002 | 0.963 | 0.002 |
| C/E x 75s/150s x L/R | 1 | | 0.837 | 0.363 | 0.837 | 0.245 | 0.622 | 0.245 | | 0.170 | 0.681 | 0.170 | 0.035 | 0.852 | 0.035 | | 0.000 | 0.998 | 0.000 | 0.000 | 0.989 | 0.000 |
| 0m/2m x 75s/150s x L/R | 1 | | 0.432 | 0.513 | 0.432 | 0.181 | 0.671 | 0.181 | | 0.007 | 0.934 | 0.007 | 0.001 | 0.980 | 0.001 | | 0.013 | 0.911 | 0.013 | 0.001 | 0.974 | 0.001 |
| C/E x 0m/2m x 75s/150s x L/R | 1 | | 0.392 | 0.533 | 0.392 | 0.039 | 0.845 | 0.039 | | 0.030 | 0.864 | 0.030 | 0.005 | 0.943 | 0.005 | | 0.004 | 0.950 | 0.004 | 0.005 | 0.942 | 0.005 |
| Group | 80 | | | | | | | | | | | | | | | | | | | | | |
| A1/A2 | 1 | | 0.591 | 0.444 | 0.591 | 0.230 | 0.633 | 0.230 | | 3.272 | 0.074 | 3.272 | 1.477 | 0.228 | 1.477 | | 11.142 | 0.001 | 11.142 | 23.198 | <.0001 | 23.198 |
| A1/A2 x C/E | 1 | | 1.183 | 0.280 | 1.183 | 2.995 | 0.087 | 2.995 | | 16.560 | 0.000 | 16.560 | 8.762 | 0.004 | 8.762 | | 9.351 | 0.003 | 9.351 | 3.333 | 0.072 | 3.333 |
| A1/A2 x 0m/2m | 1 | | 1.888 | 0.173 | 1.888 | 0.863 | 0.356 | 0.863 | | 5.336 | 0.024 | 5.336 | 0.474 | 0.493 | 0.474 | | 24.276 | <.0001 | 24.276 | 0.381 | 0.539 | 0.381 |
| A1/A2 x 75s/150s | 1 | | 0.715 | 0.400 | 0.715 | 0.001 | 0.976 | 0.001 | | 0.125 | 0.725 | 0.125 | 0.136 | 0.713 | 0.136 | | 0.121 | 0.729 | 0.121 | 0.089 | 0.767 | 0.089 |
| A1/A2 x L/R | 1 | | 1.195 | 0.278 | 1.195 | 2.542 | 0.115 | 2.542 | | 0.196 | 0.659 | 0.196 | 0.043 | 0.837 | 0.043 | | 2.354 | 0.129 | 2.354 | 3.649 | 0.060 | 3.649 |
| A1/A2 x C/E x 0m/2m | 1 | | 0.381 | 0.539 | 0.381 | 0.030 | 0.862 | 0.030 | | 0.023 | 0.880 | 0.023 | 1.896 | 0.172 | 1.896 | | 0.307 | 0.581 | 0.307 | 2.496 | 0.118 | 2.496 |
| A1/A2 x C/E x 75s/150s | 1 | | 1.693 | 0.197 | 1.693 | 0.666 | 0.417 | 0.666 | | 0.006 | 0.938 | 0.006 | 0.001 | 0.982 | 0.001 | | 0.044 | 0.834 | 0.044 | 1.084 | 0.301 | 1.084 |
| A1/A2 x C/E x L/R | 1 | | 0.345 | 0.559 | 0.345 | 0.179 | 0.673 | 0.179 | | 0.091 | 0.763 | 0.091 | 0.822 | 0.367 | 0.822 | | 0.138 | 0.712 | 0.138 | 0.622 | 0.433 | 0.622 |
| A1/A2 x 0m/2m x 75s/150s | 1 | | 1.632 | 0.205 | 1.632 | 1.777 | 0.186 | 1.777 | | 2.029 | 0.158 | 2.029 | 0.017 | 0.897 | 0.017 | | 0.190 | 0.665 | 0.190 | 0.550 | 0.460 | 0.550 |
| A1/A2 x 0m/2m x L/R | 1 | | 0.415 | 0.521 | 0.415 | 0.098 | 0.755 | 0.098 | | 0.106 | 0.746 | 0.106 | 0.138 | 0.711 | 0.138 | | 0.073 | 0.788 | 0.073 | 0.928 | 0.338 | 0.928 |
| A1/A2 x 75s/150s x L/R | 1 | | 1.436 | 0.234 | 1.436 | 0.630 | 0.430 | 0.630 | | 0.223 | 0.638 | 0.223 | 1.042 | 0.310 | 1.042 | | 0.002 | 0.963 | 0.002 | 0.039 | 0.843 | 0.039 |
| A1/A2 x C/E x 0m/2m x 75s/150s | 1 | | 0.507 | 0.478 | 0.507 | 0.491 | 0.486 | 0.491 | | 0.269 | 0.606 | 0.269 | 0.191 | 0.663 | 0.191 | | 0.084 | 0.772 | 0.084 | 0.237 | 0.628 | 0.237 |
| A1/A2 x C/E x 0m/2m x L/R | 1 | | 1.300 | 0.258 | 1.300 | 3.344 | 0.071 | 3.344 | | 0.339 | 0.562 | 0.339 | 1.485 | 0.227 | 1.485 | | 0.002 | 0.964 | 0.002 | 0.363 | 0.549 | 0.363 |
| A1/A2 x C/E x 75s/150s x L/R | 1 | | 0.902 | 0.345 | 0.902 | 0.005 | 0.943 | 0.005 | | 0.318 | 0.574 | 0.318 | 2.043 | 0.157 | 2.043 | | 0.030 | 0.864 | 0.030 | 0.002 | 0.962 | 0.002 |
| A1/A2 x 0m/2m x 75s/150s x L/R | 1 | | 0.451 | 0.504 | 0.451 | 0.227 | 0.635 | 0.227 | | 0.044 | 0.834 | 0.044 | 0.027 | 0.870 | 0.027 | | 0.000 | 0.990 | 0.000 | 0.098 | 0.755 | 0.098 |
| A1/A2 x C/E x 0m/2m x 75s/150s x L/R | 1 | | 1.991 | 0.162 | 1.991 | 2.119 | 0.149 | 2.119 | | 0.294 | 0.589 | 0.294 | 0.005 | 0.942 | 0.005 | | 0.003 | 0.955 | 0.003 | 0.090 | 0.765 | 0.090 |
| A1/A2 x Group | 80 | | | | | | | | | | | | | | | | | | | | | |
| 5bins | 4 | RPC | 0.662 | 0.619 | 2.648 | 0.582 | 0.676 | 2.329 | RiHP | 8.870 | <.0001 | 35.480 | 4.663 | 0.001 | 18.651 | RvHP | 17.071 | <.0001 | 68.283 | 12.344 | <.0001 | 49.375 |
| 5bins x C/E | 4 | | 1.511 | 0.199 | 6.046 | 2.911 | 0.022 | 11.646 | | 0.847 | 0.497 | 3.386 | 2.354 | 0.054 | 9.414 | | 2.579 | 0.037 | 10.316 | 2.893 | 0.022 | 11.573 |
| 5bins x 0m/2m | 4 | | 1.442 | 0.220 | 5.769 | 2.518 | 0.041 | 10.071 | | 1.592 | 0.176 | 6.370 | 2.325 | 0.056 | 9.299 | | 0.960 | 0.430 | 3.839 | 1.987 | 0.096 | 7.949 |
| 5bins x 75s/150s | 4 | | 0.690 | 0.599 | 2.761 | 0.336 | 0.854 | 1.342 | | 0.822 | 0.512 | 3.287 | 2.146 | 0.075 | 8.585 | | 0.482 | 0.749 | 1.930 | 0.287 | 0.886 | 1.148 |
| 5bins x L/R | 4 | | 1.233 | 0.297 | 4.932 | 2.376 | 0.052 | 9.504 | | 0.799 | 0.527 | 3.195 | 1.786 | 0.131 | 7.144 | | 1.685 | 0.153 | 6.741 | 1.039 | 0.387 | 4.157 |
| 5bins x C/E x 0m/2m | 4 | | 0.926 | 0.449 | 3.705 | 0.512 | 0.727 | 2.050 | | 0.344 | 0.848 | 1.378 | 1.097 | 0.358 | 4.387 | | 0.657 | 0.622 | 2.629 | 1.239 | 0.294 | 4.957 |
| 5bins x C/E x 75s/150s | 4 | | 1.142 | 0.337 | 4.569 | 1.109 | 0.352 | 4.437 | | 0.700 | 0.593 | 2.799 | 0.304 | 0.875 | 1.217 | | 0.509 | 0.729 | 2.036 | 0.344 | 0.848 | 1.377 |
| 5bins x C/E x L/R | 4 | | 1.149 | 0.333 | 4.597 | 0.632 | 0.640 | 2.527 | | 0.335 | 0.854 | 1.341 | 0.748 | 0.560 | 2.993 | | 0.654 | 0.624 | 2.617 | 0.370 | 0.830 | 1.479 |
| 5bins x 0m/2m x 75s/150s | 4 | | 1.128 | 0.343 | 4.511 | 0.730 | 0.572 | 2.921 | | 0.788 | 0.534 | 3.151 | 0.677 | 0.608 | 2.708 | | 0.478 | 0.752 | 1.913 | 0.606 | 0.659 | 2.422 |
| 5bins x 0m/2m x L/R | 4 | | 0.650 | 0.628 | 2.598 | 0.517 | 0.723 | 2.070 | | 0.795 | 0.529 | 3.181 | 0.841 | 0.500 | 3.366 | | 1.051 | 0.381 | 4.206 | 0.585 | 0.673 | 2.342 |
| 5bins x 75s/150s x L/R | 4 | | 0.910 | 0.458 | 3.640 | 1.084 | 0.365 | 4.335 | | 0.683 | 0.604 | 2.732 | 0.252 | 0.909 | 1.007 | | 0.119 | 0.976 | 0.477 | 0.303 | 0.876 | 1.210 |
| 5bins x C/E x 0m/2m x 75s/150s | 4 | | 0.571 | 0.684 | 2.282 | 0.307 | 0.873 | 1.230 | | 0.700 | 0.593 | 2.800 | 0.209 | 0.933 | 0.836 | | 0.559 | 0.692 | 2.237 | 0.669 | 0.614 | 2.677 |
| 5bins x C/E x 0m/2m x L/R | 4 | | 1.664 | 0.158 | 6.658 | 1.483 | 0.207 | 5.930 | | 1.370 | 0.244 | 5.479 | 0.706 | 0.588 | 2.826 | | 0.263 | 0.902 | 1.052 | 1.068 | 0.372 | 4.274 |
| 5bins x C/E x 75s/150s x L/R | 4 | | 0.603 | 0.661 | 2.411 | 0.517 | 0.723 | 2.070 | | 0.380 | 0.823 | 1.520 | 0.473 | 0.756 | 1.893 | | 0.740 | 0.565 | 2.960 | 0.278 | 0.892 | 1.111 |
| 5bins x 0m/2m x 75s/150s x L/R | 4 | | 0.498 | 0.737 | 1.991 | 0.305 | 0.875 | 1.219 | | 0.156 | 0.960 | 0.623 | 0.333 | 0.856 | 1.332 | | 0.806 | 0.522 | 3.226 | 0.372 | 0.829 | 1.487 |
| 5bins x C/E x 0m/2m x 75s/150s x L/R | 4 | | 1.131 | 0.342 | 4.523 | 0.605 | 0.659 | 2.420 | | 0.146 | 0.965 | 0.582 | 0.171 | 0.953 | 0.686 | | 0.211 | 0.932 | 0.845 | 0.918 | 0.454 | 3.671 |
| 5bins x Group | 320 | | | | | | | | | | | | | | | | | | | | | |
| A1/A2 x 5bins | 4 | | 1.666 | 0.158 | 6.665 | 1.120 | 0.347 | 4.480 | | 12.265 | <.0001 | 49.061 | 24.108 | <.0001 | 96.433 | | 35.322 | <.0001 | 141.289 | 6.799 | <.0001 | 27.195 |
| A1/A2 x 5bins x C/E | 4 | | 0.752 | 0.557 | 3.009 | 0.693 | 0.597 | 2.771 | | 1.758 | 0.137 | 7.032 | 1.788 | 0.131 | 7.151 | | 3.936 | 0.004 | 15.744 | 2.658 | 0.033 | 10.632 |
| A1/A2 x 5bins x 0m/2m | 4 | | 0.901 | 0.464 | 3.603 | 0.697 | 0.595 | 2.788 | | 0.859 | 0.489 | 3.437 | 2.984 | 0.019 | 11.936 | | 2.199 | 0.069 | 8.794 | 3.482 | 0.008 | 13.926 |
| A1/A2 x 5bins x 75s/150s | 4 | | 1.491 | 0.205 | 5.966 | 0.677 | 0.609 | 2.706 | | 0.120 | 0.975 | 0.481 | 0.663 | 0.618 | 2.652 | | 0.343 | 0.849 | 1.372 | 0.200 | 0.938 | 0.801 |
| A1/A2 x 5bins x L/R | 4 | | 0.509 | 0.729 | 2.037 | 0.345 | 0.848 | 1.380 | | 0.674 | 0.610 | 2.697 | 0.631 | 0.641 | 2.524 | | 0.572 | 0.683 | 2.288 | 0.784 | 0.536 | 3.135 |
| A1/A2 x 5bins x C/E x 0m/2m | 4 | | 1.223 | 0.301 | 4.891 | 1.708 | 0.148 | 6.834 | | 0.177 | 0.950 | 0.707 | 0.816 | 0.516 | 3.265 | | 0.221 | 0.926 | 0.885 | 2.350 | 0.054 | 9.400 |
| A1/A2 x 5bins x C/E x 75s/150s | 4 | | 0.537 | 0.709 | 2.146 | 0.392 | 0.815 | 1.567 | | 0.248 | 0.911 | 0.993 | 0.516 | 0.724 | 2.064 | | 0.725 | 0.576 | 2.898 | 0.456 | 0.768 | 1.823 |
| A1/A2 x 5bins x C/E x L/R | 4 | | 1.433 | 0.223 | 5.730 | 1.689 | 0.152 | 6.755 | | 0.228 | 0.923 | 0.910 | 0.026 | 0.999 | 0.106 | | 0.160 | 0.959 | 0.638 | 0.641 | 0.634 | 2.563 |
| A1/A2 x 5bins x 0m/2m x 75s/150s | 4 | | 0.955 | 0.432 | 3.821 | 0.639 | 0.635 | 2.555 | | 0.716 | 0.581 | 2.865 | 0.316 | 0.867 | 1.263 | | 0.264 | 0.901 | 1.056 | 0.815 | 0.517 | 3.258 |
| A1/A2 x 5bins x 0m/2m x L/R | 4 | | 1.066 | 0.374 | 4.263 | 1.586 | 0.178 | 6.344 | | 0.170 | 0.954 | 0.678 | 0.479 | 0.751 | 1.917 | | 0.222 | 0.926 | 0.886 | 0.244 | 0.913 | 0.976 |
| A1/A2 x 5bins x 75s/150s x L/R | 4 | | 0.553 | 0.697 | 2.212 | 0.485 | 0.747 | 1.939 | | 0.147 | 0.964 | 0.590 | 0.018 | 0.999 | 0.072 | | 0.197 | 0.940 | 0.786 | 0.254 | 0.907 | 1.015 |
| A1/A2 x 5bins x C/E x 0m/2m x 75s/150s | 4 | | 1.034 | 0.390 | 4.135 | 1.169 | 0.325 | 4.675 | | 0.195 | 0.941 | 0.781 | 0.386 | 0.819 | 1.542 | | 0.108 | 0.980 | 0.431 | 1.026 | 0.394 | 4.103 |
| A1/A2 x 5bins x C/E x 0m/2m x L/R | 4 | | 1.029 | 0.392 | 4.117 | 1.106 | 0.354 | 4.422 | | 0.431 | 0.786 | 1.724 | 0.091 | 0.985 | 0.366 | | 0.184 | 0.947 | 0.737 | 0.572 | 0.683 | 2.289 |
| A1/A2 x 5bins x C/E x 75s/150s x L/R | 4 | | 1.121 | 0.346 | 4.485 | 0.771 | 0.545 | 3.084 | | 0.223 | 0.926 | 0.892 | 0.476 | 0.753 | 1.904 | | 0.206 | 0.935 | 0.824 | 0.322 | 0.864 | 1.286 |
| A1/A2 x 5bins x 0m/2m x 75s/150s x L/R | 4 | | 1.115 | 0.349 | 4.460 | 1.339 | 0.255 | 5.358 | | 0.307 | 0.873 | 1.229 | 0.189 | 0.944 | 0.756 | | 0.037 | 0.997 | 0.149 | 0.160 | 0.958 | 0.642 |
| A1/A2 x 5bins x C/E x 0m/2m x 75s/150s x L/R | 4 | | 0.815 | 0.5162 | 3.261 | 0.977 | 0.420 | 3.910 | | 0.291 | 0.8835 | 1.165 | 0.248 | 0.911 | 0.994 | | 0.093 | 0.985 | 0.372 | 0.744 | 0.563 | 2.977 |
| A1/A2 x 5bins x Group | 320 | | | | | | | | | | | | | | | | | | | | | |

Cells are shaded for p < 0.05: CE | 0m2m | 75s/750s | L/R | bins/period

**Table S5. ANOVA table of coherence in both hemispheres during the 10-s period surrounding arm entry.** Coherence in the 8-10 Hz and 51-100 Hz bands across brain regions was analyzed using the same repeated-measures ANOVA as in Table S4. See Table S4 for details on factors, significance criteria, and abbreviations.

| Coherence around T-maze arm entry | df | LPC_LiHP 8-10 Hz | | | LPC_LiHP 51-100 Hz | | | LPC_LvHP 8-10 Hz | | | LPC_LvHP 51-100 Hz | | | RPC_RiHP 8-10 Hz | | | RPC_RiHP 51-100 Hz | | | RPC_RvHP 8-10 Hz | | | RPC_RvHP 51-100 Hz | | |
|---|---|---|---|---|---|---|---|---|---|---|---|---|---|---|---|---|---|---|---|---|---|---|---|---|---|
| | | F-value | p-value | Lam-bda | F-value | p-value | Lam-bda | F-value | p-value | Lam-bda | F-value | p-value | Lam-bda | F-value | p-value | Lam-bda | F-value | p-value | Lam-bda | F-value | p-value | Lam-bda | F-value | p-value | Lam-bda |
| C/E | 1 | 2.524 | 0.116 | 2.524 | 4.206 | 0.044 | 4.206 | 2.943 | 0.090 | 2.943 | 48.806 | <.0001 | 48.806 | 0.575 | 0.451 | 0.575 | 1.771 | 0.187 | 1.771 | 5.795 | 0.018 | 5.795 | 5.308 | 0.024 | 5.308 |
| 0m/2m | 1 | 0.409 | 0.524 | 0.409 | 0.083 | 0.774 | 0.083 | 0.078 | 0.780 | 0.078 | 0.851 | 0.359 | 0.851 | 0.202 | 0.654 | 0.202 | 0.009 | 0.926 | 0.009 | 0.007 | 0.935 | 0.007 | 0.057 | 0.811 | 0.057 |
| 75s/150s | 1 | 0.002 | 0.964 | 0.002 | 0.259 | 0.612 | 0.259 | 0.067 | 0.797 | 0.067 | 1.104 | 0.297 | 1.104 | 0.004 | 0.950 | 0.004 | 0.0002 | 0.9888 | 0.0002 | 0.071 | 0.790 | 0.071 | 0.096 | 0.757 | 0.096 |
| L/R | 1 | 0.056 | 0.814 | 0.056 | 0.192 | 0.662 | 0.192 | 0.025 | 0.874 | 0.025 | 0.229 | 0.634 | 0.229 | 0.0001 | 0.991 | 0.0001 | 0.001 | 0.9701 | 0.001 | 0.002 | 0.964 | 0.002 | 0.055 | 0.815 | 0.055 |
| C/E x 0m/2m | 1 | 0.029 | 0.865 | 0.029 | 0.0001 | 0.992 | 0.0001 | 0.002 | 0.962 | 0.002 | 0.322 | 0.572 | 0.322 | 0.0005 | 0.983 | 0.0005 | 7E-06 | 0.998 | 7E-06 | 0.275 | 0.601 | 0.275 | 0.004 | 0.950 | 0.004 |
| C/E x 75s/150s | 1 | 0.025 | 0.876 | 0.025 | 0.267 | 0.607 | 0.267 | 0.206 | 0.651 | 0.206 | 1.253 | 0.266 | 1.253 | 0.001 | 0.978 | 0.001 | 0.012 | 0.9127 | 0.012 | 0.016 | 0.900 | 0.016 | 0.08 | 0.778 | 0.08 |
| C/E x L/R | 1 | 0.168 | 0.683 | 0.168 | 0.108 | 0.744 | 0.108 | 0.057 | 0.812 | 0.057 | 0.195 | 0.660 | 0.195 | 0.006 | 0.939 | 0.006 | 4E-05 | 0.9951 | 4E-05 | 0.039 | 0.844 | 0.039 | 0.035 | 0.851 | 0.035 |
| 0m/2m x 75s/150s | 1 | 0.012 | 0.912 | 0.012 | 0.025 | 0.875 | 0.025 | 0.015 | 0.902 | 0.015 | 0.119 | 0.731 | 0.119 | 0.006 | 0.939 | 0.006 | 0.012 | 0.912 | 0.012 | 0.065 | 0.800 | 0.065 | 0.053 | 0.818 | 0.053 |
| 0m/2m x L/R | 1 | 0.071 | 0.791 | 0.071 | 0.218 | 0.642 | 0.218 | 0.131 | 0.718 | 0.131 | 2.007 | 0.161 | 2.007 | 0.057 | 0.812 | 0.057 | 0.154 | 0.696 | 0.154 | 0.031 | 0.861 | 0.031 | 0.261 | 0.611 | 0.261 |
| 75s/150s x L/R | 1 | 0.019 | 0.892 | 0.019 | 0.065 | 0.800 | 0.065 | 0.034 | 0.855 | 0.034 | 1.061 | 0.306 | 1.061 | 0.03 | 0.864 | 0.03 | 0.047 | 0.830 | 0.047 | 0.005 | 0.947 | 0.005 | 0.115 | 0.736 | 0.115 |
| C/E x 0m/2m x 75s/150s | 1 | 0.004 | 0.947 | 0.004 | 0.0005 | 0.983 | 0.0005 | 0.001 | 0.979 | 0.001 | 0.33 | 0.567 | 0.33 | 0.003 | 0.956 | 0.003 | 0.009 | 0.925 | 0.009 | 0.162 | 0.689 | 0.162 | 0.081 | 0.776 | 0.081 |
| C/E x 0m/2m x L/R | 1 | 0.214 | 0.645 | 0.214 | 0.418 | 0.520 | 0.418 | 0.089 | 0.766 | 0.089 | 2.161 | 0.146 | 2.161 | 0.007 | 0.932 | 0.007 | 0.134 | 0.715 | 0.134 | 0.067 | 0.797 | 0.067 | 0.22 | 0.640 | 0.22 |
| C/E x 75s/150s x L/R | 1 | 0.023 | 0.880 | 0.023 | 0.024 | 0.876 | 0.024 | 2E-05 | 0.997 | 2E-05 | 1.237 | 0.270 | 1.237 | 2E-05 | 0.997 | 2E-05 | 0.017 | 0.8963 | 0.017 | 0.016 | 0.898 | 0.016 | 0.125 | 0.724 | 0.125 |
| 0m/2m x 75s/150s x L/R | 1 | 0.383 | 0.538 | 0.383 | 0.293 | 0.590 | 0.293 | 0.003 | 0.954 | 0.003 | 0.381 | 0.539 | 0.381 | 0.001 | 0.981 | 0.001 | 0.001 | 0.9757 | 0.001 | 0.08 | 0.778 | 0.08 | 0.001 | 0.973 | 0.001 |
| C/E x 0m/2m x 75s/150s x L/R | 1 | 0.035 | 0.853 | 0.035 | 0.017 | 0.895 | 0.017 | 0.011 | 0.916 | 0.011 | 0.303 | 0.584 | 0.303 | 6E-06 | 0.998 | 6E-06 | 0.0003 | 0.986 | 0.0003 | 0.106 | 0.745 | 0.106 | 0.0001 | 0.991 | 0.0001 |
| Group | 80 | | | | | | | | | | | | | | | | | | | | | | | | |
| A1/A2 | 1 | 0.268 | 0.606 | 0.268 | 3.184 | 0.078 | 3.184 | 0.016 | 0.901 | 0.016 | 0.001 | 0.974 | 0.001 | 1.18 | 0.281 | 1.18 | 16.966 | <.0001 | 16.966 | 0.97 | 0.328 | 0.97 | 11.431 | 0.001 | 11.431 |
| A1/A2 x C/E | 1 | 8.094 | 0.006 | 8.094 | 0.163 | 0.688 | 0.163 | 14.01 | 0.0003 | 14.01 | 0.039 | 0.843 | 0.039 | 0.256 | 0.614 | 0.256 | 6.971 | 0.010 | 6.971 | 2.593 | 0.111 | 2.593 | 4.87 | 0.030 | 4.87 |
| A1/A2 x 0m/2m | 1 | 2.045 | 0.157 | 2.045 | 2.627 | 0.109 | 2.627 | 0.104 | 0.748 | 0.104 | 4.939 | 0.029 | 4.939 | 3.789 | 0.055 | 3.789 | 4.276 | 0.042 | 4.276 | 4.315 | 0.041 | 4.315 | 3.07 | 0.084 | 3.07 |
| A1/A2 x 75s/150s | 1 | 0.007 | 0.933 | 0.007 | 0.287 | 0.594 | 0.287 | 0.009 | 0.926 | 0.009 | 0.324 | 0.571 | 0.324 | 1.965 | 0.165 | 1.965 | 0.69 | 0.409 | 0.69 | 0.017 | 0.897 | 0.017 | 0.397 | 0.531 | 0.397 |
| A1/A2 x L/R | 1 | 0.019 | 0.890 | 0.019 | 0.559 | 0.457 | 0.559 | 2.115 | 0.150 | 2.115 | 2.349 | 0.129 | 2.349 | 3.79 | 0.055 | 3.79 | 2.34 | 0.130 | 2.34 | 4.509 | 0.037 | 4.509 | 1.494 | 0.225 | 1.494 |
| A1/A2 x C/E x 0m/2m | 1 | 1.311 | 0.256 | 1.311 | 0.534 | 0.467 | 0.534 | 0.317 | 0.575 | 0.317 | 2.058 | 0.155 | 2.058 | 0.517 | 0.474 | 0.517 | 0.029 | 0.866 | 0.029 | 0.351 | 0.555 | 0.351 | 0.037 | 0.848 | 0.037 |
| A1/A2 x C/E x 75s/150s | 1 | 0.134 | 0.715 | 0.134 | 2.044 | 0.157 | 2.044 | 0.333 | 0.565 | 0.333 | 0.22 | 0.641 | 0.22 | 0.035 | 0.853 | 0.035 | 1.832 | 0.180 | 1.832 | 0.28 | 0.598 | 0.28 | 0.558 | 0.457 | 0.558 |
| A1/A2 x C/E x L/R | 1 | 0.161 | 0.690 | 0.161 | 0.0002 | 0.990 | 0.0002 | 2.964 | 0.089 | 2.964 | 2.021 | 0.159 | 2.021 | 0.596 | 0.443 | 0.596 | 0.729 | 0.396 | 0.729 | 0.056 | 0.814 | 0.056 | 0.188 | 0.666 | 0.188 |
| A1/A2 x 0m/2m x 75s/150s | 1 | 0.41 | 0.524 | 0.41 | 1.035 | 0.312 | 1.035 | 0.925 | 0.339 | 0.925 | 0.026 | 0.873 | 0.026 | 0.7 | 0.405 | 0.7 | 0.261 | 0.611 | 0.261 | 0.173 | 0.679 | 0.173 | 0.065 | 0.800 | 0.065 |
| A1/A2 x 0m/2m x L/R | 1 | 1.27 | 0.263 | 1.27 | 1.421 | 0.237 | 1.421 | 0.239 | 0.626 | 0.239 | 1.27 | 0.263 | 1.27 | 1.928 | 0.169 | 1.928 | 0.39 | 0.534 | 0.39 | 0.306 | 0.582 | 0.306 | 0.112 | 0.738 | 0.112 |
| A1/A2 x 75s/150s x L/R | 1 | 0.422 | 0.518 | 0.422 | 0.698 | 0.406 | 0.698 | 0.004 | 0.951 | 0.004 | 0.197 | 0.659 | 0.197 | 0.422 | 0.518 | 0.422 | 0.104 | 0.748 | 0.104 | 0.78 | 0.380 | 0.78 | 0.534 | 0.467 | 0.534 |
| A1/A2 x C/E x 0m/2m x 75s/150s | 1 | 0.484 | 0.489 | 0.484 | 0.078 | 0.781 | 0.078 | 1.325 | 0.253 | 1.325 | 1.166 | 0.284 | 1.166 | 0.361 | 0.550 | 0.361 | 2.081 | 0.153 | 2.081 | 0.023 | 0.880 | 0.023 | 0.348 | 0.557 | 0.348 |
| A1/A2 x C/E x 0m/2m x L/R | 1 | 1.592 | 0.211 | 1.592 | 1.174 | 0.282 | 1.174 | 0.349 | 0.556 | 0.349 | 0.085 | 0.771 | 0.085 | 0.831 | 0.365 | 0.831 | 0.108 | 0.743 | 0.108 | 0.204 | 0.653 | 0.204 | 0.02 | 0.888 | 0.02 |
| A1/A2 x C/E x 75s/150s x L/R | 1 | 0.556 | 0.458 | 0.556 | 2.483 | 0.119 | 2.483 | 0.238 | 0.627 | 0.238 | 0.139 | 0.710 | 0.139 | 0.046 | 0.830 | 0.046 | 0.068 | 0.795 | 0.068 | 0.647 | 0.424 | 0.647 | 0.345 | 0.558 | 0.345 |
| A1/A2 x 0m/2m x 75s/150s x L/R | 1 | 0.558 | 0.457 | 0.558 | 0.229 | 0.633 | 0.229 | 2.065 | 0.155 | 2.065 | 0.354 | 0.554 | 0.354 | 0.04 | 0.841 | 0.04 | 0.389 | 0.534 | 0.389 | 0.106 | 0.746 | 0.106 | 0.458 | 0.501 | 0.458 |
| A1/A2 x C/E x 0m/2m x 75s/150s x L/R | 1 | 0.14 | 0.709 | 0.14 | 0.009 | 0.927 | 0.009 | 0.516 | 0.475 | 0.516 | 0.206 | 0.651 | 0.206 | 0.103 | 0.749 | 0.103 | 0.008 | 0.930 | 0.008 | 0.087 | 0.769 | 0.087 | 0.258 | 0.613 | 0.258 |
| A1/A2 x Group | 80 | | | | | | | | | | | | | | | | | | | | | | | | |
| 5bins | 4 | 6.228 | <.0001 | 24.914 | 1.311 | 0.266 | 5.246 | 4.811 | 0.001 | 19.245 | 0.8 | 0.526 | 3.2 | 1.402 | 0.233 | 5.607 | 2.394 | 0.051 | 9.576 | 8.101 | <.0001 | 32.403 | 1.208 | 0.307 | 4.833 |
| 5bins x C/E | 4 | 3.273 | 0.012 | 13.092 | 1.882 | 0.113 | 7.527 | 3.421 | 0.009 | 13.685 | 1.168 | 0.325 | 4.672 | 3.147 | 0.015 | 12.588 | 5.67 | 0.0002 | 22.679 | 0.541 | 0.706 | 2.163 | 4.465 | 0.002 | 17.859 |
| 5bins x 0m/2m | 4 | 5.534 | 0.0003 | 22.137 | 1.711 | 0.147 | 6.846 | 2.31 | 0.058 | 9.24 | 0.656 | 0.623 | 2.623 | 0.976 | 0.421 | 3.903 | 0.825 | 0.510 | 3.3 | 2.662 | 0.033 | 10.649 | 1.738 | 0.141 | 6.95 |
| 5bins x 75s/150s | 4 | 1.154 | 0.331 | 4.616 | 1.56 | 0.185 | 6.24 | 0.783 | 0.537 | 3.133 | 1.428 | 0.225 | 5.71 | 0.254 | 0.907 | 1.018 | 0.144 | 0.965 | 0.578 | 0.485 | 0.747 | 1.94 | 0.567 | 0.687 | 2.267 |
| 5bins x L/R | 4 | 1.216 | 0.304 | 4.864 | 0.129 | 0.972 | 0.516 | 0.648 | 0.628 | 2.593 | 0.487 | 0.746 | 1.946 | 1.467 | 0.212 | 5.867 | 1.218 | 0.303 | 4.874 | 0.609 | 0.657 | 2.435 | 2.067 | 0.085 | 8.266 |
| 5bins x C/E x 0m/2m | 4 | 1.255 | 0.288 | 5.022 | 1.442 | 0.220 | 5.769 | 0.146 | 0.965 | 0.585 | 1.167 | 0.325 | 4.668 | 1.318 | 0.263 | 5.271 | 1.404 | 0.232 | 5.617 | 0.885 | 0.473 | 3.542 | 1.73 | 0.143 | 6.922 |
| 5bins x C/E x 75s/150s | 4 | 2.502 | 0.042 | 10.01 | 1.576 | 0.181 | 6.302 | 1.51 | 0.199 | 6.04 | 0.84 | 0.500 | 3.362 | 0.584 | 0.675 | 2.335 | 0.786 | 0.535 | 3.143 | 0.188 | 0.945 | 0.753 | 0.686 | 0.602 | 2.746 |
| 5bins x C/E x L/R | 4 | 0.504 | 0.733 | 2.016 | 0.57 | 0.685 | 2.279 | 0.318 | 0.866 | 1.273 | 0.339 | 0.852 | 1.354 | 0.962 | 0.429 | 3.848 | 0.393 | 0.814 | 1.57 | 0.106 | 0.980 | 0.426 | 0.796 | 0.529 | 3.182 |
| 5bins x 0m/2m x 75s/150s | 4 | 1.183 | 0.318 | 4.734 | 0.98 | 0.419 | 3.92 | 0.191 | 0.943 | 0.765 | 1.896 | 0.111 | 7.584 | 0.756 | 0.555 | 3.023 | 0.404 | 0.806 | 1.615 | 0.792 | 0.531 | 3.168 | 0.292 | 0.883 | 1.169 |
| 5bins x 0m/2m x L/R | 4 | 1.399 | 0.234 | 5.596 | 2.781 | 0.027 | 11.125 | 0.631 | 0.641 | 2.524 | 1.808 | 0.127 | 7.232 | 0.237 | 0.917 | 0.949 | 1.044 | 0.385 | 4.176 | 2.495 | 0.043 | 9.978 | 1.22 | 0.302 | 4.88 |
| 5bins x 75s/150s x L/R | 4 | 0.261 | 0.903 | 1.042 | 1.302 | 0.269 | 5.21 | 0.211 | 0.932 | 0.843 | 3.025 | 0.018 | 12.1 | 0.947 | 0.437 | 3.787 | 0.44 | 0.780 | 1.761 | 0.174 | 0.951 | 0.698 | 0.258 | 0.905 | 1.031 |
| 5bins x C/E x 0m/2m x 75s/150s | 4 | 0.975 | 0.421 | 3.9 | 1.658 | 0.160 | 6.633 | 0.348 | 0.846 | 1.39 | 0.742 | 0.564 | 2.97 | 0.48 | 0.751 | 1.919 | 0.374 | 0.827 | 1.498 | 1.048 | 0.382 | 4.194 | 1.17 | 0.324 | 4.682 |
| 5bins x C/E x 0m/2m x L/R | 4 | 0.634 | 0.638 | 2.538 | 0.194 | 0.941 | 0.777 | 0.344 | 0.848 | 1.378 | 1.909 | 0.109 | 7.636 | 1.909 | 0.109 | 7.634 | 2.597 | 0.036 | 10.387 | 1.066 | 0.374 | 4.263 | 1.603 | 0.173 | 6.411 |
| 5bins x C/E x 75s/150s x L/R | 4 | 0.271 | 0.897 | 1.084 | 0.431 | 0.786 | 1.724 | 0.723 | 0.577 | 2.894 | 2.565 | 0.038 | 10.259 | 1.032 | 0.391 | 4.128 | 0.296 | 0.881 | 1.184 | 0.585 | 0.674 | 2.341 | 0.563 | 0.690 | 2.253 |
| 5bins x 0m/2m x 75s/150s x L/R | 4 | 0.799 | 0.527 | 3.194 | 1.647 | 0.162 | 6.587 | 0.649 | 0.628 | 2.596 | 0.899 | 0.465 | 3.596 | 0.931 | 0.446 | 3.725 | 0.448 | 0.774 | 1.792 | 0.556 | 0.695 | 2.226 | 0.261 | 0.903 | 1.042 |
| 5bins x C/E x 0m/2m x 75s/150s x L/R | 4 | 0.326 | 0.861 | 1.304 | 0.442 | 0.779 | 1.767 | 0.78 | 0.539 | 3.12 | 1.504 | 0.201 | 6.015 | 0.402 | 0.808 | 1.606 | 0.514 | 0.726 | 2.056 | 0.301 | 0.877 | 1.205 | 0.291 | 0.884 | 1.165 |
| 5bins x Group | 320 | | | | | | | | | | | | | | | | | | | | | | | | |
| A1/A2 x 5bins | 4 | 4.447 | 0.002 | 17.787 | 1.685 | 0.153 | 6.741 | 12.72 | <.0001 | 50.88 | 2.828 | 0.025 | 11.313 | 10.108 | <.0001 | 40.432 | 2.302 | 0.059 | 9.208 | 26.184 | <.0001 | 104.74 | 4.588 | 0.001 | 18.352 |
| A1/A2 x 5bins x C/E | 4 | 10.292 | <.0001 | 41.17 | 0.643 | 0.632 | 2.571 | 9.648 | <.0001 | 38.594 | 0.496 | 0.739 | 1.982 | 4.775 | 0.001 | 19.102 | 0.954 | 0.433 | 3.816 | 3.521 | 0.008 | 14.085 | 1.893 | 0.111 | 7.572 |
| A1/A2 x 5bins x 0m/2m | 4 | 5.437 | 0.0003 | 21.75 | 1.267 | 0.283 | 5.067 | 0.383 | 0.821 | 1.531 | 2.065 | 0.085 | 8.261 | 1.743 | 0.140 | 6.971 | 2.039 | 0.089 | 8.155 | 1.092 | 0.361 | 4.366 | 1.229 | 0.299 | 4.914 |
| A1/A2 x 5bins x 75s/150s | 4 | 1.044 | 0.385 | 4.176 | 0.906 | 0.460 | 3.626 | 1.818 | 0.125 | 7.274 | 2.534 | 0.040 | 10.135 | 0.716 | 0.582 | 2.864 | 0.93 | 0.447 | 3.72 | 2.012 | 0.093 | 8.048 | 1.282 | 0.277 | 5.126 |
| A1/A2 x 5bins x L/R | 4 | 0.873 | 0.481 | 3.49 | 0.312 | 0.870 | 1.249 | 1.381 | 0.240 | 5.526 | 2.032 | 0.090 | 8.129 | 0.343 | 0.849 | 1.373 | 3.287 | 0.012 | 13.149 | 1.362 | 0.247 | 5.449 | 1.157 | 0.330 | 4.627 |
| A1/A2 x 5bins x C/E x 0m/2m | 4 | 1.044 | 0.385 | 4.175 | 1.426 | 0.225 | 5.702 | 0.38 | 0.823 | 1.521 | 2.349 | 0.054 | 9.396 | 1.265 | 0.284 | 5.061 | 0.112 | 0.978 | 0.447 | 0.165 | 0.956 | 0.659 | 0.155 | 0.961 | 0.621 |
| A1/A2 x 5bins x C/E x 75s/150s | 4 | 0.784 | 0.536 | 3.138 | 1.405 | 0.232 | 5.621 | 2.346 | 0.055 | 9.385 | 1.184 | 0.318 | 4.737 | 0.723 | 0.577 | 2.893 | 0.187 | 0.945 | 0.75 | 0.509 | 0.729 | 2.036 | 0.73 | 0.572 | 2.918 |
| A1/A2 x 5bins x C/E x L/R | 4 | 1.47 | 0.211 | 5.882 | 0.422 | 0.793 | 1.689 | 0.645 | 0.630 | 2.582 | 0.779 | 0.540 | 3.116 | 0.15 | 0.963 | 0.602 | 0.284 | 0.888 | 1.137 | 0.922 | 0.452 | 3.686 | 0.963 | 0.428 | 3.851 |
| A1/A2 x 5bins x 0m/2m x 75s/150s | 4 | 1.705 | 0.149 | 6.819 | 0.525 | 0.717 | 2.1 | 1.142 | 0.337 | 4.566 | 0.783 | 0.537 | 3.133 | 1.121 | 0.347 | 4.484 | 2.366 | 0.053 | 9.466 | 1.013 | 0.401 | 4.051 | 1.057 | 0.378 | 4.226 |
| A1/A2 x 5bins x 0m/2m x L/R | 4 | 0.525 | 0.717 | 2.1 | 0.974 | 0.422 | 3.894 | 1.514 | 0.198 | 6.056 | 1.156 | 0.330 | 4.623 | 0.78 | 0.539 | 3.119 | 1.211 | 0.306 | 4.843 | 0.541 | 0.706 | 2.165 | 0.654 | 0.624 | 2.616 |
| A1/A2 x 5bins x 75s/150s x L/R | 4 | 1.136 | 0.339 | 4.545 | 0.713 | 0.584 | 2.85 | 0.363 | 0.835 | 1.452 | 1.419 | 0.227 | 5.676 | 1.017 | 0.399 | 4.067 | 0.78 | 0.539 | 3.121 | 0.138 | 0.968 | 0.552 | 0.058 | 0.994 | 0.232 |
| A1/A2 x 5bins x C/E x 0m/2m x 75s/150s | 4 | 0.683 | 0.604 | 2.731 | 2.405 | 0.050 | 9.619 | 0.651 | 0.626 | 2.605 | 1.697 | 0.150 | 6.787 | 0.918 | 0.453 | 3.674 | 1.489 | 0.205 | 5.956 | 0.114 | 0.978 | 0.456 | 0.45 | 0.773 | 1.799 |
| A1/A2 x 5bins x C/E x 0m/2m x L/R | 4 | 0.05 | 0.995 | 0.199 | 1.155 | 0.331 | 4.619 | 0.107 | 0.980 | 0.429 | 1.037 | 0.388 | 4.15 | 0.639 | 0.635 | 2.557 | 0.854 | 0.492 | 3.417 | 0.802 | 0.524 | 3.21 | 2.504 | 0.042 | 10.017 |
| A1/A2 x 5bins x C/E x 75s/150s x L/R | 4 | 0.324 | 0.862 | 1.298 | 0.613 | 0.653 | 2.453 | 0.652 | 0.626 | 2.609 | 0.795 | 0.529 | 3.179 | 0.885 | 0.473 | 3.541 | 0.913 | 0.456 | 3.653 | 0.08 | 0.988 | 0.322 | 0.086 | 0.987 | 0.343 |
| A1/A2 x 5bins x 0m/2m x 75s/150s x L/R | 4 | 0.64 | 0.634 | 2.561 | 0.586 | 0.673 | 2.344 | 0.241 | 0.915 | 0.962 | 1.284 | 0.276 | 5.138 | 0.671 | 0.613 | 2.683 | 0.56 | 0.692 | 2.241 | 0.028 | 0.998 | 0.114 | 0.409 | 0.802 | 1.636 |
| A1/A2 x 5bins x C/E x 0m/2m x 75s/150s x L/R | 4 | 0.184 | 0.946 | 0.738 | 1.688 | 0.152 | 6.754 | 1.046 | 0.384 | 4.183 | 0.541 | 0.706 | 2.163 | 0.443 | 0.777 | 1.774 | 0.81 | 0.520 | 3.24 | 0.72 | 0.579 | 2.882 | 0.044 | 0.996 | 0.175 |
| A1/A2 x 5bins x Group | 320 | | | | | | | | | | | | | | | | | | | | | | | | |

**Table S6. ANOVA table of PLV and MI between PC and iHP/vHP in both hemispheres during the 10-s period surrounding arm entry.** PLV and MI between iHP/vHP (8-10 Hz) and PC (51-100 Hz) were analyzed using the same repeated-measures ANOVA as in Table S4. See Table S4 for details on factors, significance criteria, and abbreviations.

**PLV**

| HP 8-10Hz_PC 51-100Hz around T-maze arm entry | df | LiHP 8-10Hz_LPC 51-100Hz | | | LvHP 8-10Hz_LPC 51-100Hz | | | RiHP 8-10Hz_RPC 51-100Hz | | | RvHP 8-10Hz_RPC 51-100Hz | | |
|---|---|---|---|---|---|---|---|---|---|---|---|---|---|
| | | F-value | p-value | Lam-bda | F-value | p-value | Lam-bda | F-value | p-value | Lam-bda | F-value | p-value | Lam-bda |
| C/E | 1 | 69.320 | <.0001 | 69.320 | 63.377 | <.0001 | 63.377 | 79.817 | <.0001 | 79.817 | 72.979 | <.0001 | 72.979 |
| 0m/2m | 1 | 0.016 | 0.899 | 0.016 | 0.499 | 0.482 | 0.499 | 0.051 | 0.822 | 0.051 | 0.074 | 0.786 | 0.074 |
| 75s/150s | 1 | 1.070 | 0.304 | 1.070 | 0.182 | 0.670 | 0.182 | 4.128 | 0.046 | 4.128 | 0.238 | 0.627 | 0.238 |
| L/R | 1 | 0.342 | 0.560 | 0.342 | 0.328 | 0.569 | 0.328 | 0.038 | 0.846 | 0.038 | 2.015 | 0.160 | 2.015 |
| C/E x 0m/2m | 1 | 0.000 | 0.988 | 0.000 | 0.047 | 0.830 | 0.047 | 1.651 | 0.203 | 1.651 | 0.191 | 0.664 | 0.191 |
| C/E x 75s/150s | 1 | 1.421 | 0.237 | 1.421 | 1.491 | 0.226 | 1.491 | 2.917 | 0.092 | 2.917 | 2.353 | 0.129 | 2.353 |
| C/E x L/R | 1 | 0.105 | 0.746 | 0.105 | 0.271 | 0.604 | 0.271 | 0.762 | 0.385 | 0.762 | 1.010 | 0.318 | 1.010 |
| 0m/2m x 75s/150s | 1 | 0.149 | 0.701 | 0.149 | 0.423 | 0.518 | 0.423 | 0.260 | 0.611 | 0.260 | 0.278 | 0.600 | 0.278 |
| 0m/2m x L/R | 1 | 0.932 | 0.337 | 0.932 | 0.002 | 0.968 | 0.002 | 0.716 | 0.400 | 0.716 | 1.165 | 0.284 | 1.165 |
| 75s/150s x L/R | 1 | 0.094 | 0.760 | 0.094 | 0.002 | 0.962 | 0.002 | 0.112 | 0.739 | 0.112 | 0.077 | 0.782 | 0.077 |
| C/E x 0m/2m x 75s/150s | 1 | 0.120 | 0.730 | 0.120 | 1.286 | 0.260 | 1.286 | 0.117 | 0.733 | 0.117 | 1.939 | 0.168 | 1.939 |
| C/E x 0m/2m x L/R | 1 | 1.547 | 0.217 | 1.547 | 0.200 | 0.656 | 0.200 | 1.271 | 0.263 | 1.271 | 0.653 | 0.422 | 0.653 |
| C/E x 75s/150s x L/R | 1 | 0.434 | 0.512 | 0.434 | 0.009 | 0.925 | 0.009 | 0.998 | 0.321 | 0.998 | 1.207 | 0.275 | 1.207 |
| 0m/2m x 75s/150s x L/R | 1 | 0.131 | 0.718 | 0.131 | 0.043 | 0.836 | 0.043 | 0.645 | 0.424 | 0.645 | 0.044 | 0.834 | 0.044 |
| C/E x 0m/2m x 75s/150s x L/R | 1 | 0.991 | 0.322 | 0.991 | 0.067 | 0.796 | 0.067 | 0.691 | 0.408 | 0.691 | 3.076 | 0.083 | 3.076 |
| Group | 80 | | | | | | | | | | | | |
| A1/A2 | 1 | 0.708 | 0.403 | 0.708 | 0.007 | 0.934 | 0.007 | 0.703 | 0.404 | 0.703 | 1.203 | 0.276 | 1.203 |
| A1/A2 x C/E | 1 | 0.809 | 0.371 | 0.809 | 0.440 | 0.509 | 0.440 | 4.000 | 0.049 | 4.000 | 6.184 | 0.015 | 6.184 |
| A1/A2 x 0m/2m | 1 | 0.231 | 0.632 | 0.231 | 0.158 | 0.692 | 0.158 | 0.024 | 0.877 | 0.024 | 4.520 | 0.037 | 4.520 |
| A1/A2 x 75s/150s | 1 | 0.025 | 0.876 | 0.025 | 0.128 | 0.722 | 0.128 | 0.229 | 0.633 | 0.229 | 0.009 | 0.925 | 0.009 |
| A1/A2 x L/R | 1 | 0.408 | 0.525 | 0.408 | 0.000 | 0.996 | 0.000 | 0.064 | 0.802 | 0.064 | 0.701 | 0.405 | 0.701 |
| A1/A2 x C/E x 0m/2m | 1 | 0.769 | 0.383 | 0.769 | 0.856 | 0.358 | 0.856 | 0.306 | 0.582 | 0.306 | 0.013 | 0.909 | 0.013 |
| A1/A2 x C/E x 75s/150s | 1 | 0.646 | 0.424 | 0.646 | 4.281 | 0.042 | 4.281 | 0.800 | 0.374 | 0.800 | 0.005 | 0.943 | 0.005 |
| A1/A2 x C/E x L/R | 1 | 0.034 | 0.854 | 0.034 | 0.075 | 0.784 | 0.075 | 0.753 | 0.388 | 0.753 | 0.283 | 0.596 | 0.283 |
| A1/A2 x 0m/2m x 75s/150s | 1 | 0.425 | 0.516 | 0.425 | 1.230 | 0.271 | 1.230 | 0.830 | 0.365 | 0.830 | 0.117 | 0.733 | 0.117 |
| A1/A2 x 0m/2m x L/R | 1 | 0.759 | 0.386 | 0.759 | 2.223 | 0.140 | 2.223 | 1.440 | 0.234 | 1.440 | 0.797 | 0.375 | 0.797 |
| A1/A2 x 75s/150s x L/R | 1 | 0.007 | 0.936 | 0.007 | 4.043 | 0.048 | 4.043 | 0.153 | 0.697 | 0.153 | 1.201 | 0.277 | 1.201 |
| A1/A2 x C/E x 0m/2m x 75s/150s | 1 | 2.193 | 0.143 | 2.193 | 0.976 | 0.326 | 0.976 | 0.044 | 0.835 | 0.044 | 0.477 | 0.492 | 0.477 |
| A1/A2 x C/E x 0m/2m x L/R | 1 | 0.303 | 0.584 | 0.303 | 0.069 | 0.793 | 0.069 | 0.244 | 0.623 | 0.244 | 0.062 | 0.803 | 0.062 |
| A1/A2 x C/E x 75s/150s x L/R | 1 | 0.514 | 0.475 | 0.514 | 0.011 | 0.919 | 0.011 | 1.023 | 0.315 | 1.023 | 0.773 | 0.382 | 0.773 |
| A1/A2 x 0m/2m x 75s/150s x L/R | 1 | 1.025 | 0.315 | 1.025 | 6.916 | 0.010 | 6.916 | 7.362 | 0.008 | 7.362 | 1.366 | 0.246 | 1.366 |
| A1/A2 x C/E x 0m/2m x 75s/150s x L/R | 1 | 0.038 | 0.846 | 0.038 | 0.343 | 0.560 | 0.343 | 0.647 | 0.424 | 0.647 | 0.351 | 0.555 | 0.351 |
| A1/A2 x Group | 80 | | | | | | | | | | | | |
| 5bins | 4 | 0.908 | 0.460 | 3.632 | 0.644 | 0.632 | 2.576 | 0.575 | 0.681 | 2.301 | 0.315 | 0.868 | 1.261 |
| 5bins x C/E | 4 | 0.575 | 0.681 | 2.299 | 1.109 | 0.352 | 4.435 | 0.163 | 0.957 | 0.651 | 1.057 | 0.378 | 4.227 |
| 5bins x 0m/2m | 4 | 4.151 | 0.003 | 16.604 | 0.202 | 0.937 | 0.809 | 0.779 | 0.539 | 3.117 | 1.361 | 0.247 | 5.445 |
| 5bins x 75s/150s | 4 | 0.832 | 0.506 | 3.328 | 1.124 | 0.345 | 4.498 | 0.657 | 0.623 | 2.627 | 1.552 | 0.187 | 6.207 |
| 5bins x L/R | 4 | 1.681 | 0.154 | 6.724 | 2.857 | 0.024 | 11.430 | 0.332 | 0.857 | 1.327 | 1.065 | 0.374 | 4.259 |
| 5bins x C/E x 0m/2m | 4 | 1.128 | 0.343 | 4.514 | 0.190 | 0.943 | 0.761 | 1.302 | 0.269 | 5.208 | 1.448 | 0.218 | 5.793 |
| 5bins x C/E x 75s/150s | 4 | 0.690 | 0.600 | 2.758 | 0.541 | 0.706 | 2.165 | 0.412 | 0.800 | 1.647 | 1.693 | 0.151 | 6.770 |
| 5bins x C/E x L/R | 4 | 1.427 | 0.225 | 5.709 | 1.711 | 0.147 | 6.843 | 0.695 | 0.596 | 2.780 | 0.684 | 0.603 | 2.737 |
| 5bins x 0m/2m x 75s/150s | 4 | 0.564 | 0.689 | 2.254 | 0.678 | 0.608 | 2.713 | 3.780 | 0.005 | 15.119 | 0.296 | 0.881 | 1.182 |
| 5bins x 0m/2m x L/R | 4 | 0.684 | 0.604 | 2.735 | 0.337 | 0.853 | 1.350 | 1.561 | 0.185 | 6.243 | 0.503 | 0.733 | 2.013 |
| 5bins x 75s/150s x L/R | 4 | 1.184 | 0.318 | 4.736 | 1.615 | 0.170 | 6.459 | 0.278 | 0.892 | 1.111 | 1.450 | 0.217 | 5.800 |
| 5bins x C/E x 0m/2m x 75s/150s | 4 | 0.846 | 0.497 | 3.383 | 0.731 | 0.571 | 2.925 | 0.233 | 0.920 | 0.933 | 0.159 | 0.959 | 0.635 |
| 5bins x C/E x 0m/2m x L/R | 4 | 1.598 | 0.175 | 6.392 | 0.846 | 0.497 | 3.384 | 0.436 | 0.783 | 1.743 | 0.169 | 0.954 | 0.674 |
| 5bins x C/E x 75s/150s x L/R | 4 | 1.227 | 0.299 | 4.909 | 1.106 | 0.354 | 4.424 | 0.701 | 0.592 | 2.803 | 1.377 | 0.242 | 5.510 |
| 5bins x 0m/2m x 75s/150s x L/R | 4 | 0.487 | 0.745 | 1.949 | 0.784 | 0.536 | 3.135 | 0.580 | 0.677 | 2.320 | 1.110 | 0.352 | 4.442 |
| 5bins x C/E x 0m/2m x 75s/150s x L/R | 4 | 1.377 | 0.242 | 5.510 | 0.402 | 0.807 | 1.609 | 0.248 | 0.911 | 0.991 | 0.322 | 0.863 | 1.287 |
| 5bins x Group | 320 | | | | | | | | | | | | |
| A1/A2 x 5bins | 4 | 1.630 | 0.166 | 6.522 | 0.621 | 0.648 | 2.483 | 0.888 | 0.471 | 3.553 | 0.324 | 0.862 | 1.298 |
| A1/A2 x 5bins x C/E | 4 | 2.519 | 0.041 | 10.074 | 0.548 | 0.701 | 2.190 | 0.869 | 0.483 | 3.475 | 0.950 | 0.435 | 3.801 |
| A1/A2 x 5bins x 0m/2m | 4 | 0.441 | 0.779 | 1.766 | 1.080 | 0.367 | 4.319 | 0.615 | 0.652 | 2.460 | 1.217 | 0.303 | 4.869 |
| A1/A2 x 5bins x 75s/150s | 4 | 0.833 | 0.505 | 3.332 | 0.253 | 0.908 | 1.012 | 1.119 | 0.348 | 4.474 | 2.808 | 0.026 | 11.230 |
| A1/A2 x 5bins x L/R | 4 | 0.481 | 0.750 | 1.922 | 1.856 | 0.118 | 7.423 | 0.923 | 0.451 | 3.691 | 1.215 | 0.304 | 4.861 |
| A1/A2 x 5bins x C/E x 0m/2m | 4 | 0.324 | 0.862 | 1.295 | 0.886 | 0.473 | 3.543 | 1.049 | 0.382 | 4.198 | 1.780 | 0.133 | 7.120 |
| A1/A2 x 5bins x C/E x 75s/150s | 4 | 1.560 | 0.185 | 6.239 | 0.964 | 0.427 | 3.857 | 0.872 | 0.481 | 3.487 | 0.606 | 0.659 | 2.423 |
| A1/A2 x 5bins x C/E x L/R | 4 | 1.497 | 0.203 | 5.988 | 1.020 | 0.397 | 4.079 | 1.136 | 0.339 | 4.545 | 1.963 | 0.100 | 7.851 |
| A1/A2 x 5bins x 0m/2m x 75s/150s | 4 | 1.324 | 0.261 | 5.297 | 1.893 | 0.111 | 7.573 | 1.293 | 0.273 | 5.173 | 1.400 | 0.234 | 5.599 |
| A1/A2 x 5bins x 0m/2m x L/R | 4 | 0.411 | 0.800 | 1.646 | 1.280 | 0.278 | 5.119 | 0.330 | 0.858 | 1.320 | 1.227 | 0.299 | 4.909 |
| A1/A2 x 5bins x 75s/150s x L/R | 4 | 0.762 | 0.551 | 3.048 | 0.197 | 0.940 | 0.789 | 1.208 | 0.307 | 4.832 | 0.723 | 0.577 | 2.892 |
| A1/A2 x 5bins x C/E x 0m/2m x 75s/150s | 4 | 1.570 | 0.182 | 6.281 | 1.393 | 0.236 | 5.572 | 1.276 | 0.279 | 5.104 | 2.056 | 0.086 | 8.224 |
| A1/A2 x 5bins x C/E x 0m/2m x L/R | 4 | 1.096 | 0.359 | 4.383 | 0.891 | 0.470 | 3.563 | 1.274 | 0.280 | 5.096 | 0.334 | 0.855 | 1.337 |
| A1/A2 x 5bins x C/E x 75s/150s x L/R | 4 | 1.131 | 0.342 | 4.524 | 0.365 | 0.834 | 1.460 | 0.979 | 0.419 | 3.916 | 1.158 | 0.330 | 4.630 |
| A1/A2 x 5bins x 0m/2m x 75s/150s x L/R | 4 | 0.844 | 0.498 | 3.374 | 0.896 | 0.467 | 3.583 | 0.867 | 0.484 | 3.470 | 1.502 | 0.201 | 6.007 |
| A1/A2 x 5bins x C/E x 0m/2m x 75s/150s x L/R | 4 | 1.587 | 0.178 | 6.348 | 1.153 | 0.332 | 4.611 | 3.080 | 0.016 | 12.318 | 0.957 | 0.431 | 3.829 |
| A1/A2 x 5bins x Group | 320 | | | | | | | | | | | | |

**MI**

| HP 8-10Hz_PC 51-100Hz around T-maze arm entry | LiHP 8-10Hz_LPC 51-100Hz | | | LvHP 8-10Hz_LPC 51-100Hz | | | RiHP 8-10Hz_RPC 51-100Hz | | | RvHP 8-10Hz_RPC 51-100Hz | | |
|---|---|---|---|---|---|---|---|---|---|---|---|---|
| | F-value | p-value | Lam-bda | F-value | p-value | Lam-bda | F-value | p-value | Lam-bda | F-value | p-value | Lam-bda |
| C/E | 11.688 | 0.001 | 11.688 | 9.868 | 0.002 | 9.868 | 34.321 | <.0001 | 34.321 | 36.993 | <.0001 | 36.993 |
| 0m/2m | 0.076 | 0.783 | 0.076 | 0.063 | 0.803 | 0.063 | 0.031 | 0.861 | 0.031 | 0.190 | 0.664 | 0.190 |
| 75s/150s | 1.083 | 0.301 | 1.083 | 1.120 | 0.293 | 1.120 | 3.289 | 0.074 | 3.289 | 2.956 | 0.089 | 2.956 |
| L/R | 0.972 | 0.327 | 0.972 | 1.543 | 0.218 | 1.543 | 6.914 | 0.010 | 6.914 | 8.045 | 0.006 | 8.045 |
| C/E x 0m/2m | 0.014 | 0.905 | 0.014 | 0.114 | 0.737 | 0.114 | 0.001 | 0.981 | 0.001 | 0.137 | 0.712 | 0.137 |
| C/E x 75s/150s | 1.418 | 0.237 | 1.418 | 1.075 | 0.303 | 1.075 | 3.715 | 0.058 | 3.715 | 3.686 | 0.058 | 3.686 |
| C/E x L/R | 0.198 | 0.658 | 0.198 | 0.417 | 0.520 | 0.417 | 2.563 | 0.113 | 2.563 | 5.001 | 0.028 | 5.001 |
| 0m/2m x 75s/150s | 0.746 | 0.390 | 0.746 | 0.762 | 0.385 | 0.762 | 0.056 | 0.813 | 0.056 | 0.108 | 0.743 | 0.108 |
| 0m/2m x L/R | 0.774 | 0.382 | 0.774 | 0.705 | 0.404 | 0.705 | 1.638 | 0.204 | 1.638 | 2.432 | 0.123 | 2.432 |
| 75s/150s x L/R | 1.489 | 0.226 | 1.489 | 1.785 | 0.185 | 1.785 | 1.882 | 0.174 | 1.882 | 2.622 | 0.109 | 2.622 |
| C/E x 0m/2m x 75s/150s | 0.862 | 0.356 | 0.862 | 1.124 | 0.292 | 1.124 | 0.006 | 0.940 | 0.006 | 0.256 | 0.614 | 0.256 |
| C/E x 0m/2m x L/R | 0.384 | 0.537 | 0.384 | 0.382 | 0.538 | 0.382 | 0.831 | 0.365 | 0.831 | 1.228 | 0.271 | 1.228 |
| C/E x 75s/150s x L/R | 0.734 | 0.394 | 0.734 | 0.979 | 0.325 | 0.979 | 0.580 | 0.449 | 0.580 | 1.722 | 0.193 | 1.722 |
| 0m/2m x 75s/150s x L/R | 0.216 | 0.644 | 0.216 | 0.200 | 0.656 | 0.200 | 0.114 | 0.736 | 0.114 | 0.201 | 0.655 | 0.201 |
| C/E x 0m/2m x 75s/150s x L/R | 0.004 | 0.947 | 0.004 | 0.006 | 0.938 | 0.006 | 0.038 | 0.845 | 0.038 | 0.289 | 0.592 | 0.289 |
| Group | | | | | | | | | | | | |
| A1/A2 | 12.861 | 0.001 | 12.861 | 10.893 | 0.001 | 10.893 | 10.035 | 0.002 | 10.035 | 6.934 | 0.010 | 6.934 |
| A1/A2 x C/E | 4.354 | 0.040 | 4.354 | 2.003 | 0.161 | 2.003 | 2.892 | 0.093 | 2.892 | 3.228 | 0.076 | 3.228 |
| A1/A2 x 0m/2m | 1.564 | 0.215 | 1.564 | 1.240 | 0.269 | 1.240 | 0.408 | 0.525 | 0.408 | 1.618 | 0.207 | 1.618 |
| A1/A2 x 75s/150s | 1.009 | 0.318 | 1.009 | 1.239 | 0.269 | 1.239 | 0.800 | 0.374 | 0.800 | 0.825 | 0.367 | 0.825 |
| A1/A2 x L/R | 3.257 | 0.075 | 3.257 | 8.049 | 0.006 | 8.049 | 18.519 | <.0001 | 18.519 | 14.942 | 0.0002 | 14.942 |
| A1/A2 x C/E x 0m/2m | 0.979 | 0.325 | 0.979 | 1.865 | 0.176 | 1.865 | 0.604 | 0.440 | 0.604 | 0.109 | 0.742 | 0.109 |
| A1/A2 x C/E x 75s/150s | 0.444 | 0.507 | 0.444 | 0.829 | 0.365 | 0.829 | 0.878 | 0.352 | 0.878 | 1.265 | 0.264 | 1.265 |
| A1/A2 x C/E x L/R | 0.561 | 0.456 | 0.561 | 1.901 | 0.172 | 1.901 | 7.732 | 0.007 | 7.732 | 7.089 | 0.009 | 7.089 |
| A1/A2 x 0m/2m x 75s/150s | 0.318 | 0.574 | 0.318 | 0.030 | 0.862 | 0.030 | 0.158 | 0.692 | 0.158 | 0.144 | 0.706 | 0.144 |
| A1/A2 x 0m/2m x L/R | 0.003 | 0.954 | 0.003 | 0.549 | 0.461 | 0.549 | 0.122 | 0.728 | 0.122 | 0.000 | 0.993 | 0.000 |
| A1/A2 x 75s/150s x L/R | 1.543 | 0.218 | 1.543 | 2.207 | 0.141 | 2.207 | 1.983 | 0.163 | 1.983 | 0.353 | 0.554 | 0.353 |
| A1/A2 x C/E x 0m/2m x 75s/150s | 0.208 | 0.650 | 0.208 | 0.697 | 0.406 | 0.697 | 0.483 | 0.489 | 0.483 | 0.115 | 0.735 | 0.115 |
| A1/A2 x C/E x 0m/2m x L/R | 0.195 | 0.660 | 0.195 | 0.238 | 0.627 | 0.238 | 0.001 | 0.978 | 0.001 | 0.006 | 0.940 | 0.006 |
| A1/A2 x C/E x 75s/150s x L/R | 0.182 | 0.670 | 0.182 | 0.219 | 0.641 | 0.219 | 1.819 | 0.181 | 1.819 | 0.416 | 0.521 | 0.416 |
| A1/A2 x 0m/2m x 75s/150s x L/R | 0.052 | 0.821 | 0.052 | 0.003 | 0.960 | 0.003 | 0.028 | 0.868 | 0.028 | 0.001 | 0.972 | 0.001 |
| A1/A2 x C/E x 0m/2m x 75s/150s x L/R | 0.058 | 0.810 | 0.058 | 0.358 | 0.551 | 0.358 | 0.505 | 0.479 | 0.505 | 0.032 | 0.858 | 0.032 |
| A1/A2 x Group | | | | | | | | | | | | |
| 5bins | 0.831 | 0.506 | 3.324 | 1.696 | 0.151 | 6.784 | 3.146 | 0.015 | 12.582 | 2.965 | 0.020 | 11.862 |
| 5bins x C/E | 0.398 | 0.810 | 1.592 | 0.677 | 0.608 | 2.710 | 3.025 | 0.018 | 12.101 | 2.315 | 0.057 | 9.261 |
| 5bins x 0m/2m | 1.207 | 0.308 | 4.827 | 1.074 | 0.369 | 4.297 | 2.909 | 0.022 | 11.637 | 1.650 | 0.161 | 6.601 |
| 5bins x 75s/150s | 0.475 | 0.754 | 1.900 | 0.438 | 0.781 | 1.753 | 1.741 | 0.141 | 6.965 | 0.771 | 0.545 | 3.083 |
| 5bins x L/R | 4.714 | 0.001 | 18.857 | 3.160 | 0.014 | 12.639 | 4.959 | 0.001 | 19.835 | 3.517 | 0.008 | 14.069 |
| 5bins x C/E x 0m/2m | 1.703 | 0.149 | 6.814 | 1.053 | 0.380 | 4.210 | 1.856 | 0.118 | 7.423 | 1.332 | 0.258 | 5.328 |
| 5bins x C/E x 75s/150s | 0.545 | 0.703 | 2.178 | 0.393 | 0.814 | 1.571 | 1.140 | 0.338 | 4.559 | 0.504 | 0.733 | 2.018 |
| 5bins x C/E x L/R | 2.838 | 0.025 | 11.353 | 1.982 | 0.097 | 7.929 | 2.119 | 0.078 | 8.475 | 1.142 | 0.337 | 4.570 |
| 5bins x 0m/2m x 75s/150s | 1.104 | 0.355 | 4.417 | 1.367 | 0.245 | 5.469 | 2.596 | 0.036 | 10.385 | 1.291 | 0.273 | 5.165 |
| 5bins x 0m/2m x L/R | 0.850 | 0.495 | 3.398 | 0.657 | 0.622 | 2.627 | 0.635 | 0.638 | 2.539 | 0.591 | 0.670 | 2.362 |
| 5bins x 75s/150s x L/R | 0.562 | 0.690 | 2.248 | 0.664 | 0.617 | 2.657 | 1.825 | 0.124 | 7.302 | 1.261 | 0.285 | 5.046 |
| 5bins x C/E x 0m/2m x 75s/150s | 1.249 | 0.290 | 4.994 | 1.638 | 0.165 | 6.550 | 1.338 | 0.256 | 5.352 | 1.381 | 0.240 | 5.524 |
| 5bins x C/E x 0m/2m x L/R | 0.609 | 0.657 | 2.436 | 0.353 | 0.842 | 1.412 | 0.179 | 0.949 | 0.715 | 0.156 | 0.960 | 0.623 |
| 5bins x C/E x 75s/150s x L/R | 0.455 | 0.769 | 1.819 | 0.272 | 0.896 | 1.087 | 1.489 | 0.205 | 5.957 | 0.980 | 0.419 | 3.919 |
| 5bins x 0m/2m x 75s/150s x L/R | 0.315 | 0.868 | 1.258 | 0.299 | 0.879 | 1.195 | 0.583 | 0.675 | 2.331 | 0.512 | 0.727 | 2.047 |
| 5bins x C/E x 0m/2m x 75s/150s x L/R | 0.212 | 0.932 | 0.849 | 0.075 | 0.990 | 0.301 | 0.265 | 0.901 | 1.059 | 0.331 | 0.857 | 1.325 |
| 5bins x Group | | | | | | | | | | | | |
| A1/A2 x 5bins | 1.493 | 0.204 | 5.970 | 2.435 | 0.047 | 9.741 | 1.549 | 0.188 | 6.197 | 2.539 | 0.040 | 10.155 |
| A1/A2 x 5bins x C/E | 0.686 | 0.602 | 2.745 | 1.441 | 0.220 | 5.763 | 0.252 | 0.908 | 1.008 | 1.015 | 0.400 | 4.061 |
| A1/A2 x 5bins x 0m/2m | 1.540 | 0.190 | 6.160 | 1.230 | 0.298 | 4.919 | 2.168 | 0.073 | 8.670 | 2.042 | 0.088 | 8.167 |
| A1/A2 x 5bins x 75s/150s | 0.761 | 0.551 | 3.045 | 1.040 | 0.387 | 4.160 | 0.263 | 0.902 | 1.051 | 0.463 | 0.763 | 1.852 |
| A1/A2 x 5bins x L/R | 3.228 | 0.013 | 12.912 | 3.149 | 0.015 | 12.598 | 4.001 | 0.004 | 16.005 | 6.008 | 0.0001 | 24.032 |
| A1/A2 x 5bins x C/E x 0m/2m | 1.295 | 0.272 | 5.179 | 0.807 | 0.521 | 3.230 | 0.775 | 0.542 | 3.099 | 1.181 | 0.319 | 4.724 |
| A1/A2 x 5bins x C/E x 75s/150s | 0.333 | 0.856 | 1.333 | 0.366 | 0.833 | 1.463 | 0.190 | 0.944 | 0.760 | 0.376 | 0.826 | 1.504 |
| A1/A2 x 5bins x C/E x L/R | 2.072 | 0.084 | 8.286 | 1.830 | 0.123 | 7.319 | 1.559 | 0.185 | 6.236 | 2.821 | 0.025 | 11.285 |
| A1/A2 x 5bins x 0m/2m x 75s/150s | 0.461 | 0.765 | 1.843 | 0.339 | 0.852 | 1.355 | 0.752 | 0.557 | 3.009 | 0.610 | 0.655 | 2.442 |
| A1/A2 x 5bins x 0m/2m x L/R | 1.343 | 0.254 | 5.371 | 0.701 | 0.592 | 2.803 | 2.361 | 0.053 | 9.443 | 1.326 | 0.260 | 5.304 |
| A1/A2 x 5bins x 75s/150s x L/R | 0.891 | 0.470 | 3.564 | 1.127 | 0.344 | 4.508 | 1.352 | 0.251 | 5.408 | 2.117 | 0.079 | 8.468 |
| A1/A2 x 5bins x C/E x 0m/2m x 75s/150s | 0.798 | 0.527 | 3.193 | 0.777 | 0.541 | 3.108 | 0.226 | 0.924 | 0.905 | 0.414 | 0.799 | 1.655 |
| A1/A2 x 5bins x C/E x 0m/2m x L/R | 1.350 | 0.251 | 5.402 | 0.994 | 0.411 | 3.977 | 2.584 | 0.037 | 10.335 | 1.505 | 0.201 | 6.020 |
| A1/A2 x 5bins x C/E x 75s/150s x L/R | 0.744 | 0.563 | 2.975 | 0.830 | 0.507 | 3.319 | 1.252 | 0.289 | 5.009 | 1.903 | 0.110 | 7.610 |
| A1/A2 x 5bins x 0m/2m x 75s/150s x L/R | 0.589 | 0.671 | 2.355 | 0.354 | 0.841 | 1.417 | 2.785 | 0.027 | 11.138 | 1.788 | 0.131 | 7.150 |
| A1/A2 x 5bins x C/E x 0m/2m x 75s/150s x L/R | 0.364 | 0.834 | 1.457 | 0.368 | 0.831 | 1.474 | 1.889 | 0.112 | 7.557 | 1.545 | 0.189 | 6.180 |
| A1/A2 x 5bins x Group | | | | | | | | | | | | |
